# Information propagation through enzyme-free catalytic templating of DNA dimerization with weak product inhibition

**DOI:** 10.1101/2023.08.23.554302

**Authors:** Javier Cabello Garcia, Rakesh Mukherjee, Wooli Bae, Guy-Bart V. Stan, Thomas E. Ouldridge

## Abstract

Information propagation by sequence-specific, template-catalyzed molecular assembly is the source of the biochemical complexity of living systems. Templating allows the production of thousands of sequence-defined proteins from only 20 distinct building blocks. By contrast, exploitation of this powerful chemical motif is rare in non-biological contexts, particularly in enzyme-free environments, where even the template-catalyzed formation of dimers is a significant challenge. The main obstacle is product inhibition: the tendency of products to bind to their templates more strongly than individual monomers, preventing the effective catalytic templating of longer polymers. Here we present a rationally designed enzyme-free system in which a DNA template catalyzes, with weak competitive product inhibition, the production of sequence-specific DNA dimers. We demonstrate the selective templating of 9 different dimers with high specificity and catalytic turnover. Most importantly, our mechanism demonstrates a rational design principle for engineering information propagation by molecular templating of longer polymers.

## Introduction

Molecular templating is an extraordinarily powerful chemical process. As an example, most chemical complexity in biology arises from well-established templating processes, such as RNA transcription, protein translation, and DNA replication, where sequence information is efficiently copied from a copolymer template into a newly produced daughter copolymer. ^1^ Using these processes, cell machinery can produce tens of thousands of distinct proteins from 20 amino acids.^2^ The role of templates as information encoders is vital for producing these distinct sequence-specific protein polymers. A small set of monomer units, such as amino acids, reacting in the absence of a template would result in the formation of a heterogeneous population of random protein sequences, as they cannot encode in their chemical interactions enough information to direct the assembly of individual proteins from such a diverse catalog of possible products.^3^

Although templating reactions in the cells are governed by enzyme-catalyzed reactions, there has been wide interest in rationally engineering enzyme-free templating mechanisms to assemble specific molecules.^4^ Many researchers seek to use templating to enhance reactions that have an otherwise low yield.^5,6^ Others pursue templating as a pathway to synthesize new complex sequence-controlled polymers, ^7,8^ or even use biological polymers, like DNA, as an easily synthesized template for directing combinatorial screenings to discover new materials and molecules with therapeutic potential.^9,10^ More ambitiously, biologically-relevant polymers are used as templates to understand the origin of life or engineer synthetic life.^11–15^ When designing enzyme-free templated systems, one of the biggest challenges, rather than efficient monomer recognition, is producing templates that act effectively as a catalyst.

To ensure a reliable copying system, the reaction of monomers must be slow in solution but occur rapidly and with high turnover in the presence of the catalyst template. To achieve this high turnover, the assembled products must be efficiently released from the template to ensure the reusability of the template.^11,16^ If templates were not reusable, a new, highly-specific template would need to be assembled for each product macromolecule; creating the template itself would become a self-assembly challenge of similar magnitude to the assembly of the product, defeating the purpose of templating.^17,18^

The tendency of products to remain bound to catalysts is known as product inhibition and is observed across all types of catalysts.^19^ If the product/catalyst complex has a lifetime similar to the substrate/catalyst complex, it will tend to “compete” with the monomers for binding. In extreme cases, if product binding is irreversible, it will prevent catalysis altogether.

Product inhibition is a particular challenge for templated assembly of dimers and longer molecules. After polymerization of the monomers, the resultant product is typically bound more strongly due to the cooperative template interaction of the now interconnected monomers. Indeed, in simple models, the free-energy change of dissociation increases linearly with polymer length.^20^ This cooperative effect results in stronger inhibition as the polymer length increases.

As a result of cooperative product inhibition, the construction of enzyme-free templating catalysts has seen limited progress. Several dimer templating systems have been demonstrated, with varying degrees of product inhibition and catalytic efficiency. ^13,15,21,22^ These systems are, however, not generalizable to longer templates. They lack a mechanism for overcoming product inhibition whilst ensuring that the weakly-binding, partially-formed products do not prematurely detach from the template.^17^ Alternatively, many groups have circumvented product inhibition by cycling external conditions to first favor growth on the template and then separation of product and template. ^23,24,24–26^ Others have created environments with a non-chemical supply of energy – temperature gradients ^14^ or mechanical agitation^27^ – that allow growth and separation. Although life may have originated in a similar non-autonomous setting, with replication driven by external conditions, this requirement limits the autonomy of the templating process and can compromise the use of templates for chemical synthesis, lacking the versatility of extant autonomous, chemically-driven biological processes.

In this work, we implement, solely from DNA, sequence-specific catalytic templating of DNA dimerisation with low product inhibition. To do so, we employ a mechanism that diverts free energy from the binding of monomers to weaken the bonds of the reacting monomers with the template. Previous work in this direction has had limited efficacy,^28^ involved templates that are not reusable catalysts, ^29,30^ or has not incorporated sequence specificity. ^31^ Leveraging the handhold-mediated strand displacement (HMSD) motif introduced in Cabello-Garcia *et al*.,^29^ we demonstrate that a DNA-based template can perform sequence-specific catalysis of the formation of one out of nine different dimers in competition, with high turnover and low product inhibition. Moreover, the design is, in principle, generalizable to the templated copying of longer templates, further increasing the potential of DNA as a tool to efficiently explore the vast chemical space of sequence-defined polymers.

## Results

### Principle of HMSD-based catalysts

The proposed system consists of two sequential physical DNA reactions: toehold-mediated strand displacement (TMSD) and handhold-mediated strand displacement (HMSD). TMSD (Fig. 1a) is central to dynamic DNA nanotechnology.^32^ The reaction involves three nucleic acid strands - an invader *I*, an incumbent *C*, and a target *R*. Initially, *R* and *C* form a duplex separate from *I*. However, *R* is typically longer than *C* and presents a single-stranded “toehold” overhang. *I* is complementary to the whole of *R*, and can thus bind to the toehold and then displace *C* from its binding with *R*. Toeholds act as recognition domains for the displacement, with longer toeholds increasing the probability of successful displacement, resulting in an exponential increase in displacement rate with toehold length up to a plateau at around six-to-seven nucleotides.^33^

**Figure 1.**
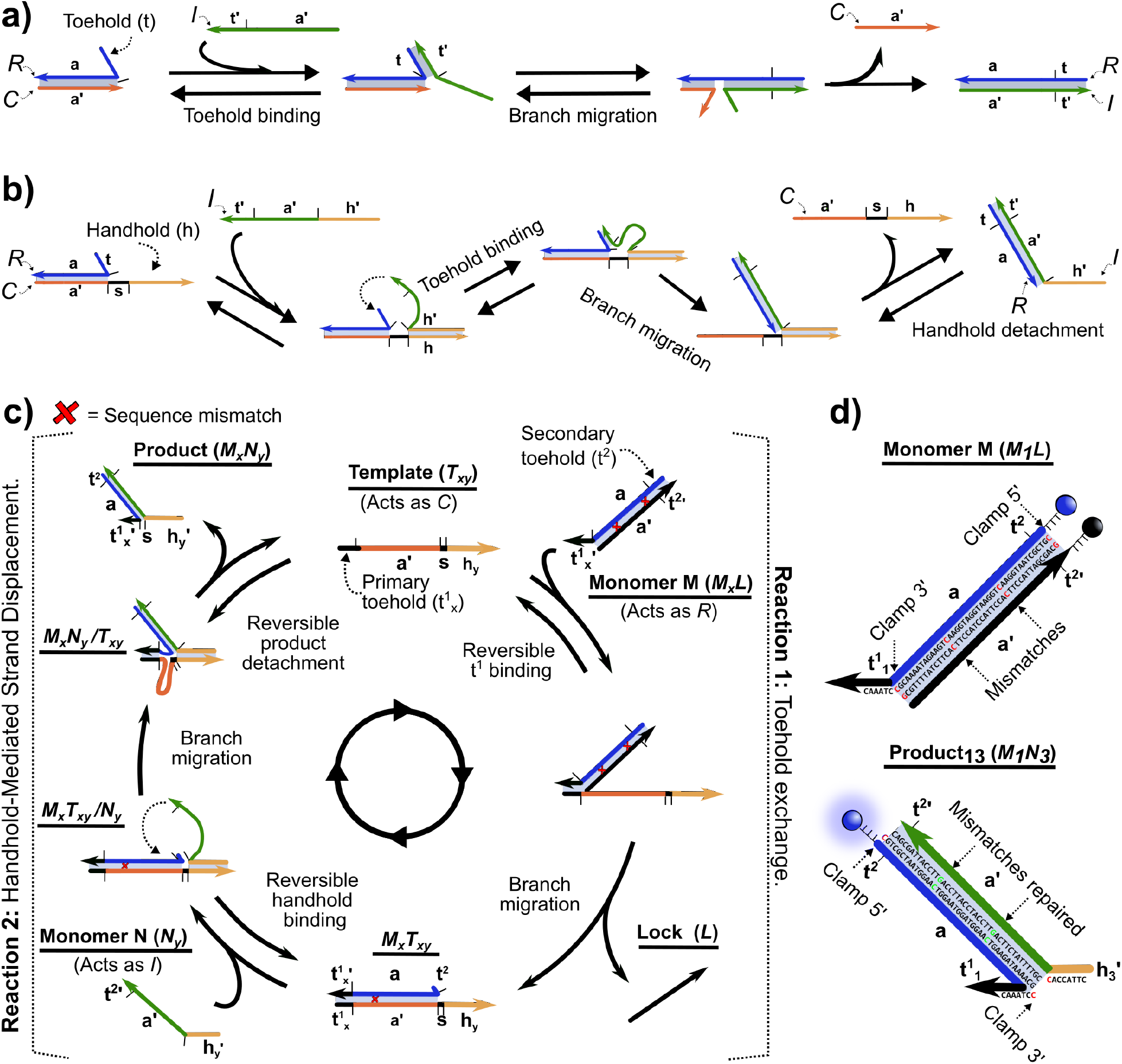
DNA strand displacement topologies, catalysis mechanism of the template, and system design. DNA strands are represented by domains (contiguous sequences of nucleotides considered to hybridize as one entity). Domains are labeled with a lowercase letter and an apostrophe (^*′*^) indicating sequence complementarity, *e*.*g. a*^*′*^ binds to *a*. **a**, Toehold-mediated strand displacement (TMSD). Binding to the toehold domain (*t*) in the target DNA strand (*R*) mediates displacement of the incumbent (*C*) by the invader (*I*). After displacement, the toehold is cooperatively sequestered in duplex *IR*. **b**, Handhold-mediated strand displacement (HMSD). When *I* binds to the handhold (*h*) domain in *C*, the effective concentration of *I* increases in the vicinity of *R*, enhancing displacement. The reversible nature of handhold binding allows *IR* to detach. **c**, DNA-based catalytic templating system. DNA monomers (*M*_*x*_*L* and *N*_*y*_) can dimerize after binding to a DNA template (*T*_*xy*_) via TMSD and HMSD, respectively. Dimerization between the monomers weakens the interaction with *T*_*xy*_, allowing *M*_*x*_*N*_*y*_ to detach and for *T*_*xy*_ to undergo another dimerization cycle. **d**, *M*_*x*_*L* and *M*_*x*_*N*_*y*_ duplexes. The dimerization domain (*a*) is initially hidden by *L*, inhibiting any direct reaction in the absence of *T*_*xy*_. The edges of the *M*_*x*_*L* duplex have additional bases – “clamps” – suppressing any leak reactions. Two mismatched base pairs in *M*_*x*_*L*’s *a* ensure that dimerization is thermodynamically favored.

HMSD (Fig. 1b) is a recently-proposed motif that adds new functionality to strand displacement networks. It operates similarly to TMSD, but the initial recognition domain (the “handhold”) is in *C* rather than *R*. This change in topology is ideal for templating; the binding of *I* to *R* can be templated by the recognition between *C* and *I*, just as recognition between DNA and RNA nucleotides templates the polymerization of RNA during transcription. Furthermore, the binding of *I* to *R* disrupts the binding of *R* to *C*; the displacement process rips apart the *CR* duplex allowing the *IR* duplex to spontaneously detach.^29^

In our proposed templating system, shown in Fig. 1c, our monomer units are two pools of DNA strands, labeled *M*_*x*_ and *N*_*y*_, homologous to *R* and *I*. Both *M*_*x*_ and *N*_*y*_ monomers possess a long “dimerization” domain and a short “recognition” domain. Here, *x, y* = 1, 2, 3… depicts the sequence identity of the monomer’s recognition domains (toehold in *M*_*x*_ and handhold in *N*_*y*_), which are recognized by a template *T*_*xy*_. The “dimerization” domains are *x, y*-independent, and complementary, so any arbitrary *M*_*x*_*N*_*y*_ dimer can form with roughly equal stability. Monomer dimerization when free in solution is suppressed by a “lock” strand *L*, complementary to the dimerization domain on *M*_*x*_. Instead, dimerization is catalyzed by a template *T*_*xy*_ containing recognition sequence domains complementary to those on both *M*_*x*_ and *N*_*y*_, and a dimerization domain complementary to the dimerization domain on *M*_*x*_. As a result, as illustrated in Fig. 1c, *T*_*xy*_ can first displace *L* from *M*_*x*_ via TMSD, and then *N*_*y*_ can displace *T*_*xy*_ from its duplex with *M*_*x*_ via HMSD, releasing the dimer *M*_*x*_*N*_*y*_ and completing catalysis. The HMSD process is facilitated by a short “secondary” toehold on *M*_*x*_ revealed during the first TMSD step.

Since *M*_*x*_*N*_*y*_ detaches from *T*_*xy*_, the overall reaction simply replaces a duplex in the *M*_*x*_*L* complex with an identical one in *M*_*x*_*N*_*y*_, with no extra-base pairs formed in the process. As stated, the process would not have any thermodynamic drive pushing it toward dimerization.

Therefore, to make the *M*_*x*_*N*_*y*_ products more stable than the reactants, without substantially increasing the rate of template-free dimerization, we incorporate two mismatched base pairs in the *M*_*x*_*L* duplexes.^34^ Each of these sequence mismatches destabilizes the *M*_*x*_*L* duplex by around 9 *k*_B_*T* relative to the *M*_*x*_*N*_*y*_ product.^20^ One of the mismatches is eliminated during the TMSD reaction, and the second during HMSD (Fig. 1d).

### Optimization of system design for catalytic turnover

The programmability of DNA-based engineering allows a systematic base-by-base optimization of the dimerization mechanism. We considered four key features of the system: the lengths of the primary toehold, handhold, and secondary toehold sequence domains and the length and structure of the dimerization domain. We considered a set of system designs using primary toeholds ranging in length from 4-8 nucleotides (nt) and handholds from 6-10 nt. Based on Ref.,^29^ we introduced a secondary toehold of 2 nt, as it ensures fast HMSD without introducing a long toehold that could trigger TMSD on its own.

Additionally, the final design of our system uses a dimerization domain as illustrated in Fig. 1d. This design has “clamp” base pairs in *M*_*x*_*L*, not present in *M*_*x*_*N*_*y*_, to reduce the spontaneous displacement of *L* from *M*_*x*_*L* by *N*_*y*_. We collect the results for an alternative design without clamps in Supplementary Results 1 (Supplementay Figures 20 and 21); this alternative design was slightly faster but suffered from a significant dimerization rate in the absence of *T*_*xy*_. As we henceforth only consider variation in the primary toehold and handhold lengths, we will use the notation *u*t/*v*h to refer to a system with a primary toehold of *u* nt and a handhold of *v* nt.

To compare the different systems with variable toehold and handhold lengths, we rank their performance attending to their initial turnover frequency (TOF), estimated from the initial rate of their dimerization reaction per the amount of *T*_*xy*_ catalyst present in the reaction solution. ^35^ The TOF indicates how effective the template catalyst *T* is at binding the substrates – via TMSD – converting them into a product – via HMSD – and then releasing that product. The experiments consisted of the injection of 10 nm *N*_3_ into a pre-equilibrated mixture of 10 nm *M*_1_*L* and 1 nm *T*_13_ (we focus on *M*_1_ and *N*_3_ when optimizing the system design). Turnover of the *M*_1_*L* duplex is tracked by fluorescence emission, as the *M*_1_*L* is labeled with a fluorophore-quencher pair, and the initially quenched signal is recovered once *L* is released from the duplex (Fig. 2a). We provide results for the kinetics of individual TMSD and HMSD steps in Supplementary Note 6.1.

**Figure 2.**
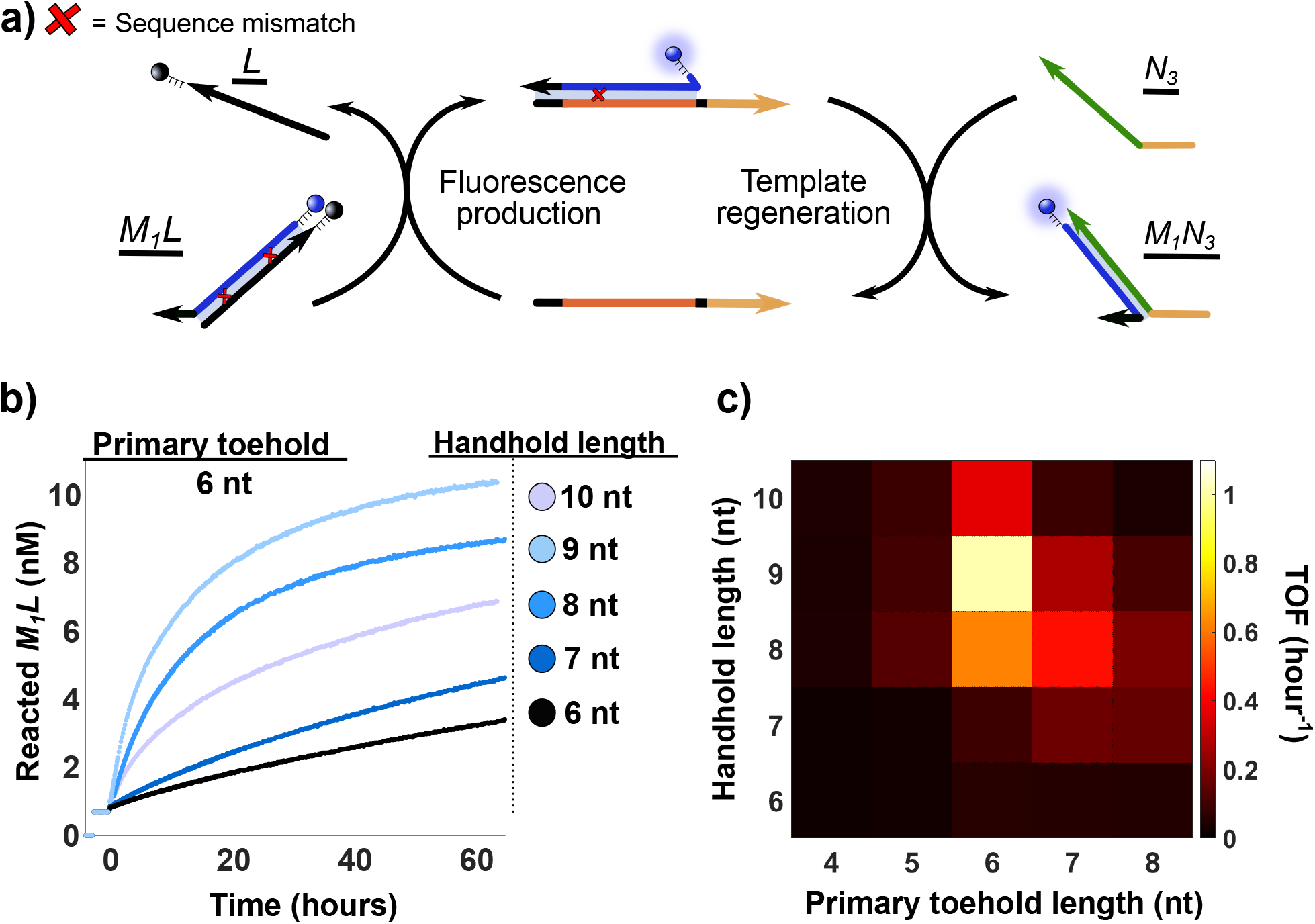
Initial turnover frequency (TOF) is optimized for toeholds and handholds of moderate length. **a**, Experimental setup. A small concentration of template (1 nm) is combined with a larger pool of *M*_1_*L* and *N*_3_ monomers (10 nm) for a range of primary toehold and handhold lengths. The catalytic turnover of *M*_1_*L* is reported by an increase in fluorescence signal. **b**, Example trajectories showing the concentration of reacted *M*_1_*L* over time, for a range of handhold lenghts and a primary toehold of 6 nt. Increasing handhold length above 9 nt results in a decrease of the *M*_1_*L* catalytic turnover due to increased product inhibition. These results illustrate how the overall reaction rate is a balance between displacement and *M*_*x*_*N*_*y*_ detachment from *T*_*xy*_. The concentration of reacted *M*_1_*L* is inferred from the fluorescence data as outlined in Supplementary Note 4. **c**, Initial rate of reaction per unit of template (TOF) for each primary toehold and handhold condition. An optimum is obtained for a system with a primary toehold of 6 nt and a handhold of 9 nt (6t/9h) (1.01 *±* 0.03 hour ^-1^) followed by condition 6t/8h (0.622 *±* 0.009 hour ^-1^).

As illustrated by the kinetics shown in Fig. 2b, 1 nm of *T*_*xy*_ is capable of triggering the transformation of high concentrations of *M*_1_*L* relative to the amount of template, indicating multiple rounds of catalytic turnover. The measured TOF, under the conditions of the experiment – far from a rate-saturating monomer concentration – reaches its maximum for 6t/9h at 1.01 *±* 0.03 molecules turned over per hour per molecule of template. Here, recognition domains are long enough to encourage binding to the template and high displacement rates but not so long that release of the product is slow, as happens for 6t/10h in Fig. 2b. It is notable from Fig. 2c that an increase in the primary toehold length tends to decrease the optimal handhold length and vice-versa, indicating the importance of minimizing the cooperative binding of *M* and *N* to *T* once they form product *MN*.

### Characterisation of the system’s resistance to product inhibition

The initial TOF metric ignores the effect of competitive product inhibition from the rebinding of products in the environment. To test the inhibition resistance of the different designs, we ran the same experiments but with variable concentrations of pre-annealed *M*_1_*N*_3_ already present in the reaction mix. The results for primary toeholds of length 5-7 nt and handholds of 8-10 nt are plotted in Fig. 3a. From the initial TOFs of the resultant kinetics, we estimated the product concentration at which the reaction’s initial rate is halved (IC_50_) as a metric to compare product inhibition.^36^ The results demonstrate the expected inverse correlation between product inhibition resistance and domain lengths, with estimated IC_50_’s of 11 nm for 6t/8h, 4 nm for 6t/9h and 2 nm for 6t/10h (Fig. 3b). We, therefore, select 6t/8h as our optimal design due to its balance between a high TOF and an IC_50_ similar to the monomer concentrations, which indicates relatively weak, non-cooperative binding to the template by the product. A screening for all *u*t/*v*h conditions and their extracted TOF values are shown in Supplementary Fig. 15 and Supplementary Table 16, respectively.

**Figure 3.**
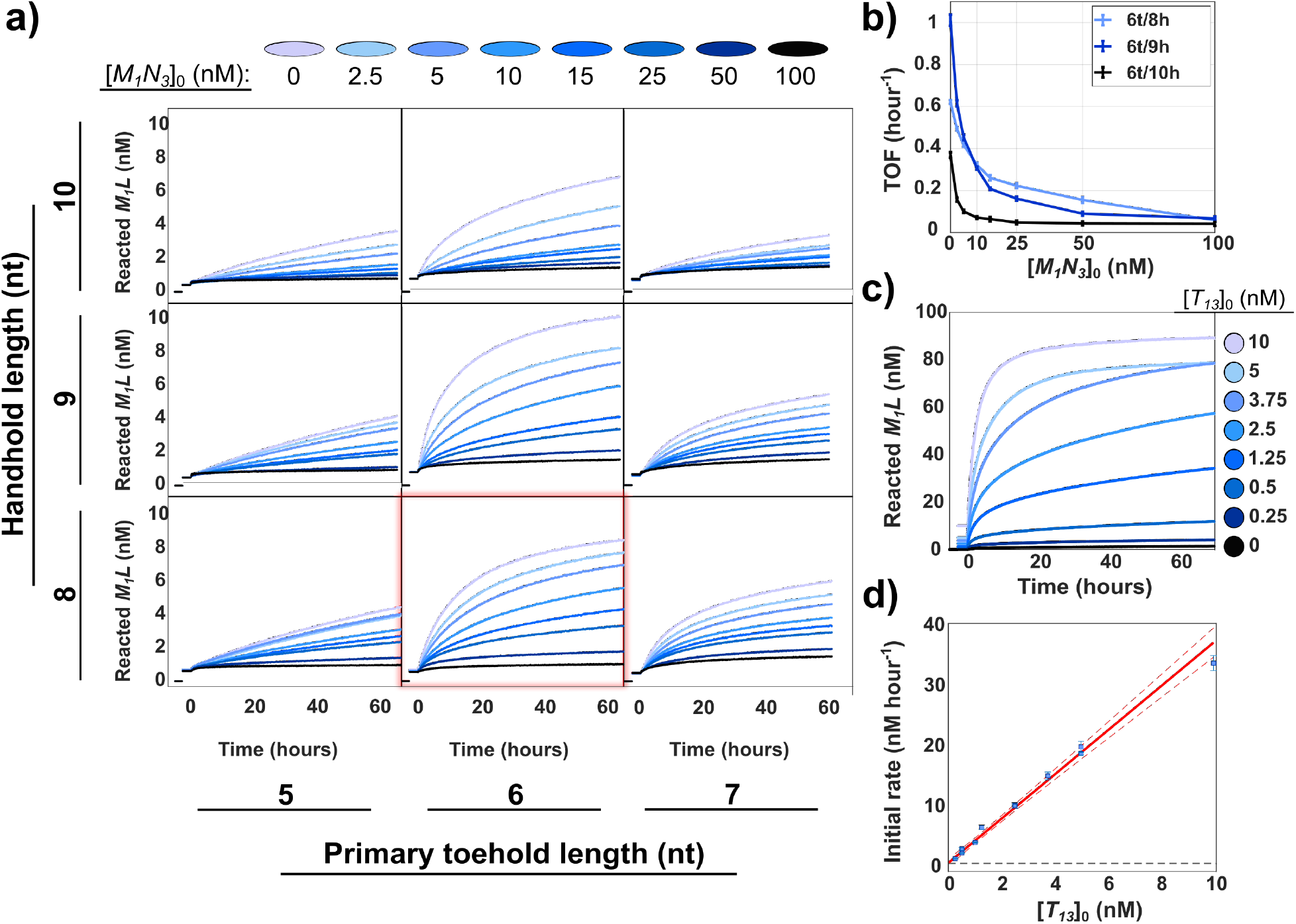
The optimal design of HMSD-based catalyst experiences only moderate competitive product inhibition and achieves high turnover. **a**, Reacted monomer concentration [*M*_1_*L*] in a system with 10 nm *N*_3_, 10nm *M*_1_*L*, 1 nm *T*, and an initial non-fluorescent pool of products *M*_1_*N*_3_ at a range of concentrations [*M*_1_*N*_3_]_0_. The condition 6t/8h, considered as optimal, is highlighted in red. **b**, Initial turnover frequency (TOF) at different [*M*_1_*N*_3_]_0_ conditions for 6t/8h, 6t/9h, and 6t/10h, obtained from the kinetics depicted in panel (a). Data reported with a 95% confidence interval. Although 6t/9h has a higher TOF in the absence of [*M*_1_*N*_3_], 6t/8h combines rapid growth with a higher resistance to rate reduction at high [*M*_1_*N*_3_]_0_. **c**, Turnover of *M*_1_*L* as inferred from fluorescence data, in experiments with 100 nm *M*_1_*L*, 100 nm *N*_3_, and variable concentrations of the template [*T*_13_]_0_ (6t/8h). A large proportion of *M*_1_*L* is observed to react, even for a concentration of [*T*_13_]_0_ 400 times lower than the number of monomers, reaching turnovers above 20 products per template. Template-free leak reactions are essentially negligible (0.32 ± 0.06 M^-1^ ± s^-1^) even compared to the lowest template concentration regimes. **d**, Initial rates of reactions from (c) and an additional set of replica experiments (Supplementary Note 6.2) plotted with a 95% interval as a function of template concentration [*T*_13_]_0_. ‘Red line’: linear fit of the system TOF (3.6 ± 0.3 hour^-1^; ‘red dashed line’: 95% confidence interval of the fit; ‘black dashed line’: untemplated rate for monomers at 100 nm = 0.012 *±* 0.002 nm hour^-1^.

To test whether the inhibition experienced by 6t/8h is strictly competitive (arising from the competition between *M*_1_*N*_3_ and *M*_1_*L* for binding to the template), we increased the initial concentration of monomers and products by an order of magnitude and catalyzed the reaction with 2.5, 5, or 10 nm of template. The three tested conditions replicate the same behavior, with its IC_50_ being independent of [*T*_*xy*_]_0_ and comparable to the concentration of the monomer (*≈* 100 nm, Supplementary Fig. 16, Supplementary Table 17).

However, a catalyst must not only have a high reaction rate, but must also be able to complete several catalytic cycles. We report catalytic turnover of the 6t/8h design using a very large monomer:template in Fig. 3c. Single templates can catalyze the assembly of at least 20-25 dimers per template. Moreover, the underlying leak rate due in the absence of a template is very slow. The template-free control is consistent, with a reaction rate of 0.32*±* 0.06, M^*−*1^ s^*−*1^ (Supplementary Table 14, Supplementary Figure 12), slower than previous measurements of toehold-free strand displacement.^33^ The dimer production signal due to the presence of templates far exceeds this leak reaction, even when templates are only present at a ratio of 1:400 with the monomers. In addition, these experiments confirm that the reaction initial rate is proportional to the template concentration (Fitted TOF for 100 nm monomers regime = 3.6 *±* 0.3 hour^-1^; Fig. 3d).

### Sequence-specific copying by templated dimerization

To demonstrate that the optimized 6t/8h design can perform information propagation by sequence-specific templating, we now consider mixtures with three distinct species per monomer type: *M*_*x*_, *N*_*y*_ with *x, y* = 1, 2, 3. Monomers of the same type share the same dimerization domain but have different template recognition domain sequences (Strand sequences are shown in Supplementary Note 1). Thus, there are 9 possible templated *M*_*x*_*N*_*y*_ of similar stability, associated with 9 catalytic templates *T*_*xy*_. If each *T*_*xy*_ templates the formation of only the dimer *N*_*x*_*M*_*y*_ from a mixture of monomers, it will have successfully copied the information in its sequence. The characterization of the 9 separate templated reactions in isolation is given in Supplementary Note 6.2.

We evaluate the success of the specific-sequence templating reaction by measuring the real-time kinetics of *M*_*x*_*L* turnover and by gel electrophoresis. To identify the possible *M*_*x*_*N*_*y*_ dimers, we label both *M*_*x*_ and *N*_*y*_ with fluorophores, as shown in Fig. 4a. As this labeling alone cannot unmistakably distinguish all 9 complexes, we also give each monomer different lengths for the poly(T) linkers connecting the fluorophores to the monomers. The different linkers allow dimers to be identified during gel electrophoresis by a combination of their migration speed during gel electrophoresis and Förster resonance energy transfer (FRET) between the fluorophores in the dimer.

**Figure 4.**
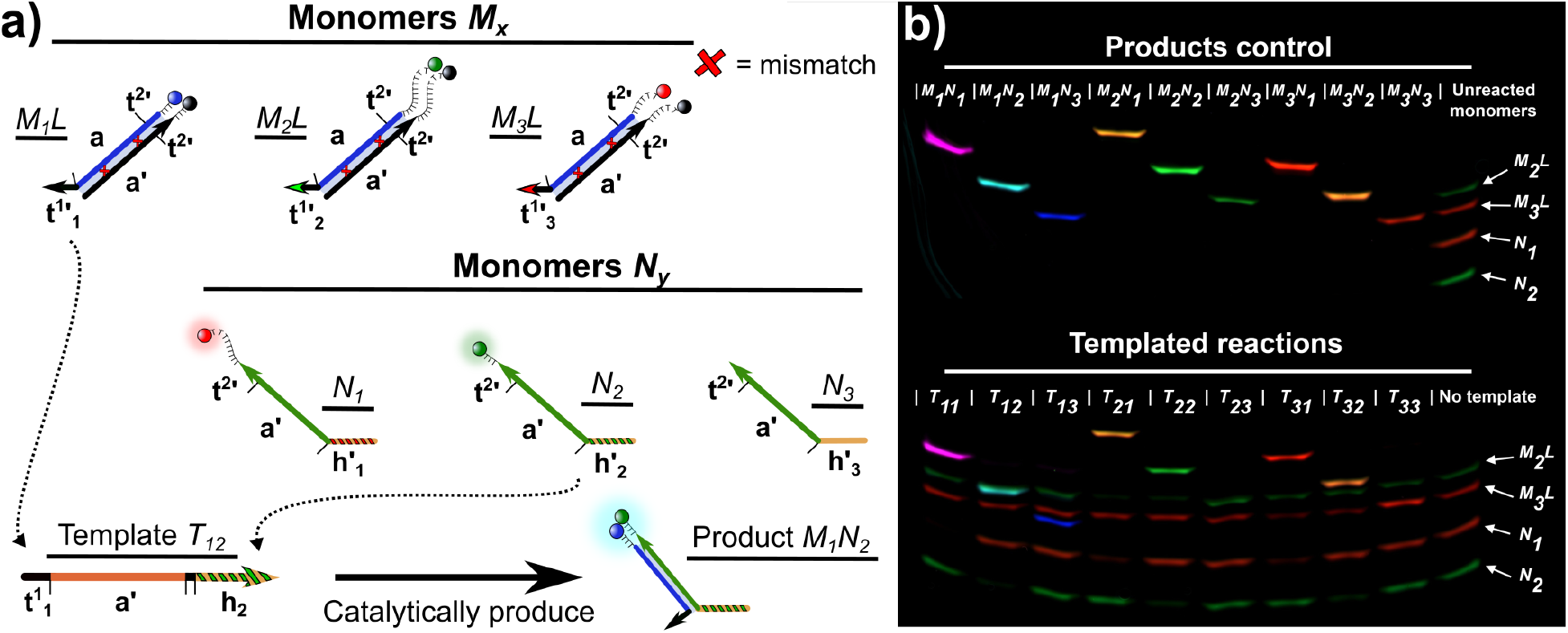
Information propagation by sequence-specific catalytic dimerization. **a**, Design of monomers to demonstrate accurate information propagation in catalytic dimerization. We consider three types of *M*_*x*_, differentiated by their primary toehold, and three types of *N*_*y*_, differentiated by their handholds. Fluorescent labeling, using poly(T) linkers of variable length, allows the identification of all *M*_*x*_*N*_*y*_ complexes through gel electrophoresis. Templates *T*_*xy*_ are intended to selectively template the formation of *M*_*x*_*N*_*y*_ from a pool of all six monomer species. **b**, Fluorescent scan of gel electrophoresis demonstrating sequence-specific templating. Products control: Signal produced by each possible *M*_*x*_*N*_*y*_ dimer produced by annealing 75 nm of each monomer. Templated reactions: Reaction mixture in which a low concentration of a single *T*_*xy*_ (5 nm) is combined with 100 nm of each *M*_*x*_*L* monomer and 75 nm of each *N*_*y*_ monomer. Observed products and signal from unreacted monomers in each well after 40 hours of reaction is consistent with the intended *M*_*x*_*N*_*y*_ production. False colors: blue, Alexa 488; green, Alexa 546; red, Alexa 647; cyan, FRET 488/546; yellow, FRET 546/648; purple, FRET 488/648)

Fig. 4b and Fig. 5 illustrate the results of experiments in which a single *T*_*xy*_ is mixed with all six monomer species. In Fig. 4b), we show polyacrylamide gel electrophoresis (PAGE) analysis of the system after 40 hours of reaction; the gels demonstrate that information is copied from template to product with a high degree of accuracy. For each *T*_*xy*_, the expected *M*_*x*_*N*_*y*_ band is visible, and its constituent monomer bands fade, with little evidence of unintended product formation.

**Figure 5.**
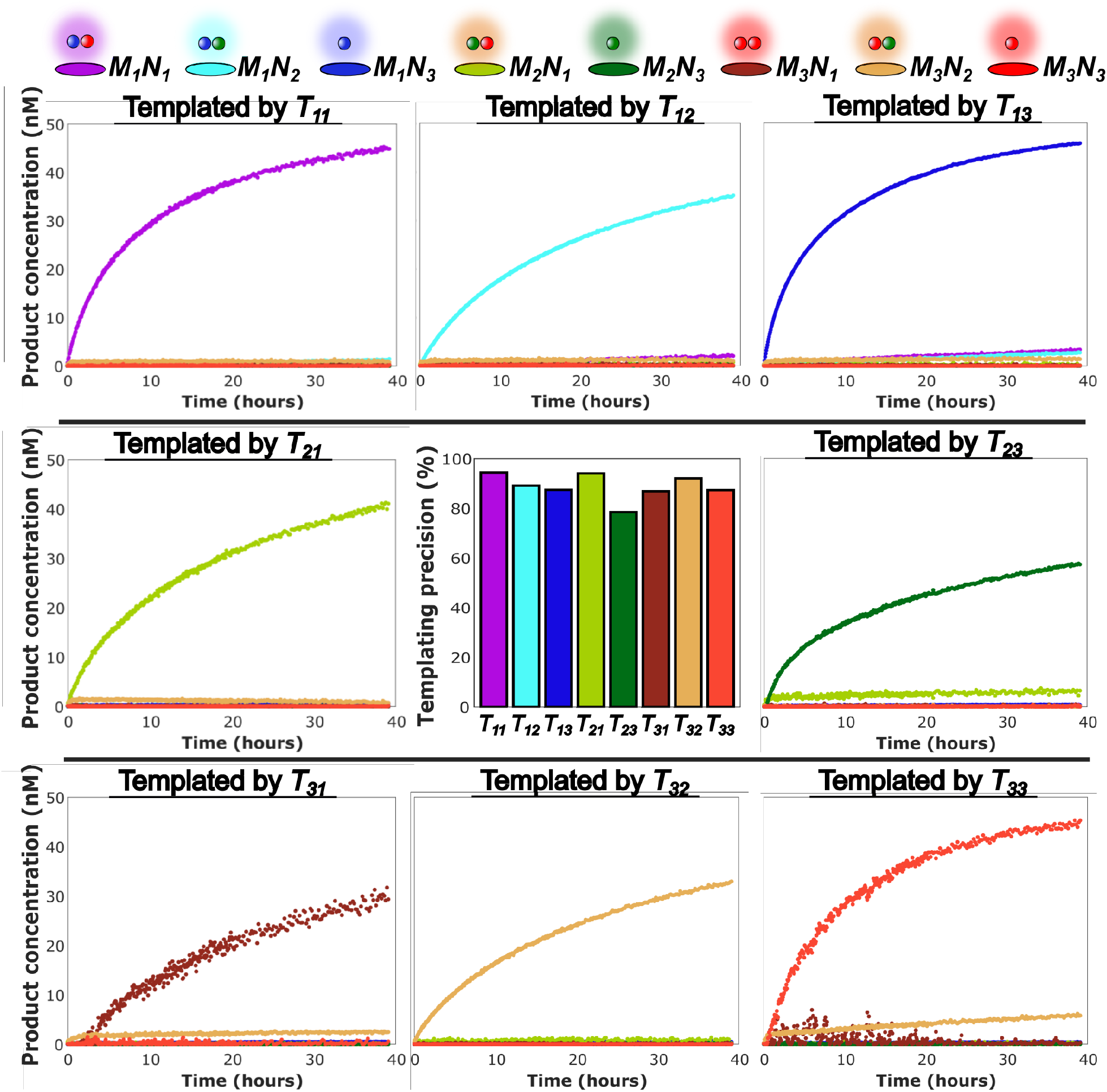
Real-time kinetics of dimer formation for sequence-specific dimerization. A mixture of six monomers and a specific template *T*_*xy*_ were mixed together to react. We plot inferred concentrations of 8 of the 9 possible dimers (*M*_2_*N*_2_ is not distinguishable from its constituent monomers via fluorescence alone) for all templates except *T*_22_. Each dimer is represented by the same color in each plot, indicated by the key at the top of the figure. The central panel shows, for each template, an estimate of the percentage of correct dimer formation after 24 hours.

Gel results are supported by the inferred concentrations from the real-time kinetics experiment (Fig. 5). We inferred the concentrations of all products – except *M*_2_*N*_2_ due to its low signal-to-noise ratio – by deconvoluting fluorescence signals as described in Supplementary Note 6.2. A quantitative analysis of the kinetic data suggests an accuracy between 80-95% for each template intended product, despite the challenge of inferring product concentrations through fluorescence alone (Supplementary Results 2, Supplementary Tables 19-21). Assuming the production of *M*_2_*N*_2_ is similar to other species, as is suggested by the PAGE data, we estimate that the mutual information between template and product is 2.45 bits, out of a maximum of 3.17 bits for perfect copying. Results for different reactant concentrations are collected in Supplementary Results 2.

## Conclusions

Using our DNA-based system, based on the principle of channeling the free energy of dimerization into disrupting binding to a template, we have demonstrated: i) weak competitive product inhibition that requires a product concentration approximately equal to the monomer concentration to halve the reaction rate; ii) a catalytic reaction around 1500 times faster than its leak rate under the conditions considered (5 nm *T*_13_; 100 nm monomers) iii) a turnover of at least 20-25 reactions per template; and iv) information propagation by highly-specific molecular templating, with an accuracy of around 90% when selecting a single product from nine alternatives.

Comparing the performance of our DNA-based system to other synthetic, sequence-specific templating mechanisms is not straightforward since the capabilities of those systems are often couched in the language of autocatalysis and self-replication. However, early biomolecular replication experiments^12,21,22,37^ typically show catalytic rates only a few times faster than spontaneous reactions, significant product inhibition at low product concentrations, or limited turnover per catalyst. More recent work^13,15,28^ has demonstrated improvements along various axes of this performance space. But unlike previous designs, our HMSD-templating system is, in principle, extendable to longer templates. Crucially, the binding of *M*_*x*_ to the template is stable (Supplementary Results 4, Supplementary Figure 29) until that binding is disrupted by dimerization. Our template for dimerization can therefore be extended by adding more sites that look like the binding site of *M*_*x*_, and copies could grow while remaining template-attached until they reach a final truncated binding site analogous to the binding site for monomer *N*_*y*_ in this work (see sketch in Supplementary Note 7, Supplementary Figure 19). Recent theoretical work has demonstrated that a system of this kind can produce copies of longer templates.^17^

Our work opens up at least three directions for further investigation. First, generalize the mechanism to longer templates. Factors not considered in the theory of Juritz *et al*.^17^ may impede this feat. However, recent work has shown reliable (but not sequence-specific) templating of tetramers using catalyzed decarboxylative aldol-type reactions using a related approach.^31^ Second, adapt the HMSD motif to self-replication. In the current work, copies carry the same information as templates but are not functional templates themselves, prohibiting self-replication. Finally, although the non-covalently bound dimers produced in our system are long-lived, a natural direction would be to combine the mechanism with covalent bond formation, either between the DNA strands themselves or using DNA-based recognition to assemble other molecules, opening new horizons for sequence-controlled polymer synthesis and drug discovery.

## Methods

### DNA sequence design

DNA sequences that minimize undesired interactions during HMSD were designed with bespoke scripts using the NUPACK server (http://www.nupack.org).^38,39^ All strands were purchased from Integrated DNA Technologies (IDT) with HPLC purification and normalized at 100 μm in LabReady buffer. All sequences are listed by function in Supplementary Note 1, Supplementary Tables 1 to 4.

### *M*_*x*_*L* duplex preparation

*M*_*x*_*L* duplexes were annealed at a concentration of 2 μm of *M*_*x*_ with a 10% excess of its corresponding *L* to ensure the sequestration of every *M*_*x*_. Annealing was performed in 100 μL of experimental buffer (TAE 1X and 1 m NaCl, pH 8.3) and annealed by heating to 95°C for 4 min and cooling to 20°C at a rate of 1°C/min.

### Bulk Fluorescence Spectroscopy

Bulk fluorescence assays were carried out in a Clariostar Microplate reader (BMG LABTECH) using flat *μ*Clear bottom 96-well plates (Greiner) and reading from the bottom. Experimental protocols were based on those previously described in Ref.^29^ Each experiment consisted of the system’s kinetics and a set of complementary measurements. These complementary measurements quantified a fluorescence baseline and estimated the concentration of each species in the system from the fluorescence signal measured after sequentially triggering the reaction of all the species. More detailed protocols for the different experiments performed in this work are provided in Supplementary Note 3 (Supplementary Tables 7-12 and Supplementary Figures 5-8). Unprocessed fluorescence signals are provided in a data upload on Zenodo (https://doi.org/10.5281/zenodo.8256587).

### Tests for optimizing the dimerization mechanism

The main tests on the kinetics of the dimerization mechanism were performed using the monomers *M*_1_ and *N*_3_, and the template *T*_13_. Unless stated otherwise, kinetic results were obtained by tracking the fluorescence of the labeled strand *M*_1_ at 25°C in a 200 μL volume. Typically, this fluorescence would increase due to the displacement of the quencher-bearing lock strand *L* in the presence of *T*_13_ and *N*_3_. Kinetics were recorded after injecting 50 μL of the reaction triggering species in 150 μL of experimental buffer containing the rest of the reactant species (pump speed: 430 μL s^*−*1^). The final mixture was shaken for 3 s (double-orbital, at 400 rpm). Injected and reacting volumes were previously preheated to the experiment temperature. Simultaneously, we recorded the fluorescence of a positive control, (*M*_1_*N*_3_) and a negative control (experimental buffer). These controls were used to correct the measured fluorescence due to temperature and volume changes. The samples were contained in Eppendorf Lobind tubes, and the plate reader’s injector system was passivated by incubating with BSA 5% for 30 min to maximize concentration reproducibility during the assays.^40^

Strand *M*_1_ was labeled with Alexa Fluor^®^ 488 (excitation: 488/14 nm; emission: 535/30 nm) and strand *L* with FQ IowaBlack quencher. Every tested system was assayed at least three times, including modifications of either *T*_13_ or *N*_3_ concentrations to extract further information from the reaction kinetics in different regimes. The fluorescence signal was averaged for 100 flashes in a spiral area scan per data point. For experiments that lasted under 1 hour, the kinetics were just averaged for 20 flashes per data point. Data from further experiments conducted during optimization, including assays on individual reaction substeps, is reported in Supplementary Note 6 (Supplementary Tables 13-18 and Supplementary Figures 10-18).

Once the basic design had been optimized through tests of variants of *M*_1_, *N*_3_ and *T*_13_, we also measured the dimerization kinetics of other combinations of *M*_*x*_, *N*_*y*_ and *T*_*xy*_ for this optimized design. The results of these experiments are reported in Supplementary Note 6 (Supplementary Tables 13, 14 and 18 and Supplementary Figures 11, 12, 17 and 18).

The additional monomer species used during these assays were labeled in the following way: *M*_2_*L*: Alexa Fluor^®^ 546 (excitation: 540/20 nm; emission: 590/30 nm) and Black Hole quencher^®^-2; *M*_3_*L*: Alexa Fluor^®^ 647 (excitation: 625/30 nm; emission: 680/30 nm) and RQ IowaBlack quencher; *N*_1_: Alexa Fluor^®^ 647; *N*_2_: Alexa Fluor^®^ 546. To monitor as many species as possible, the experiments recorded these fluorescence signals and FRET resulting from the combinations of Alexa Fluor^®^ 488/546 (excitation: 488/14 nm; emission: 590/30 nm), Alexa Fluor^®^ 488/647 (excitation: 488/14 nm; emission: 670/30 nm and Alexa Fluor^®^ 546/647 (excitation: 540/20 nm; emission: 680/30 nm).

### Kinetics of sequence-specific copying by templated dimerization

The templating of specific products required more sophisticated bulk fluorescence assays. The experiment reported in Fig. 5 consisted of ten wells loaded with an intended concentration of 100 nm of each monomer *M* (*M*_1_*L, M*_2_*L* and *M*_3_*L*). Nine of these wells also contained 5 nm of one of the nine tested templates, with the tenth containing no template to track the mechanism’s leak reaction. The reaction in each of these wells was triggered with 50 μL of a solution of the *N* monomers (*N*_1_, *N*_2_ and *N*_3_), each of them at an intended concentration of 75 nm. The experiment also included another set of ten wells. Nine of them contained positive controls for each dimer, formed by annealing 75 nm of each combination of *M* and *N* strands. The last well was an only-buffer blank control. Raw fluorescent data for this experiment is present in Supplementary Fig. 8.

Sequence-specific copying experiments were initially performed using different concentrations of the monomers and templates; see Supplementary Results 2 (Supplementary Figures 22-26). The results are qualitatively similar, though it was observed an apparent slow timescale conversion of *M*_1_*N*_3_ into *M*_1_*N*_1_ and *M*_1_*N*_2_ when *M*_*x*_*L* was not initially present in excess of *N*_*y*_. It appears that empty templates can also catalyze this interconversion reaction on slower timescales than the templating of assembly, reducing the apparent accuracy. This mechanism is further discussed in Supplementary Results 3 (Supplementary Figures 27 and 28).

### Polyacrylamide Gel Electrophoresis (PAGE)

After recording the kinetics of sequence-specific copying by templated dimerization, aliquots from the nine template conditions and the no template control were loaded into a polyacrylamide gel. A second gel was produced as a reference, using the dimers assembled in the kinetics positive control and a freshly made mixture of all unreacted monomers (*M*_*x*_*L* at 100 nm and *N*_*y*_ at 75 nm) in the experimental buffer. The gel used was a pre-cast Novex™ 10% 37.5:1 acrylamide: bisacrylamide gel in TBE 1X (Invitrogen™). The samples were mixed with native gel loading dye solution 10x (Invitrogen™), and 15 μL of the mixture was loaded in each well. The gels were run in an X-Cell SureLock™ electrophoresis chamber, using TBE 1X + 50 mm NaCl as running buffer, and a program of 10 V/cm for 30 minutes and 15 V/cm for 90 minutes. The tank was kept in an ice bath during the electrophoresis to avoid sample heating. To avoid band distortions due to the difference in ionic strength between buffer (50 mm NaCl) and samples (900 mm NaCl), the loaded samples were left in the wells for at least 30 minutes before starting the electrophoresis. ^41^

The gels were imaged with a Typhoon™ gel scanner (Amersham™). The false colors reported in Fig. 4b correspond to the following fluorescence measurements: blue (Excitation: 488 nm; emission: 525/20 nm), green (excitation: 532 nm; emission: 570/20 nm), red (excitation: 635 nm; emission: 670/30 nm), cyan (excitation: 488 nm; emission: 570/20 nm), yellow (excitation: 532 nm; emission: 670/30 nm) and purple (excitation: 488 nm; emission: 670/30 nm). The results reported in Fig. 4b and Supplementary Figures 23 and 25 are an overlay of these six scans. Raw data for each scan is contained on Zenodo (https://doi.org/10.5281/zenodo.8256587).

### Calibration of Fluorescent Signals

Fluorescence calibrations were made for all the fluorescence species used in this work. Calibrations aimed to estimate the units of fluorescence produced per nm of each complex containing a fluorophore-labeled species to quantify its concentration during the experiments. Calibration curves ranged from 15 nm to 150 nm, in 200 μL volumes, from stock solutions normalised at 100 μm. Additional calibrations tested the variation of fluorescence of *M*_1_ when bound to the template. The calibration protocol and results are given in Supplementary Note 2 (Supplementary Figures 1-4, Supplementary Tables 5 and 6).

## Data Processing

The data from bulk fluorescence experiments were corrected with each experiment’s positive and negative controls and transformed from fluorescence units to concentrations of the relevant species using fluorescence calibrations. The transformation procedures are described in detail in Supplementary Note 4 (Supplementary Figure 9), and the scripts are available at the Zenodo upload (https://doi.org/10.5281/zenodo.8256587).

## Data Fitting

Simple models of reaction kinetics (Supplementary Note 5) were used to fit reaction rate constants to the processed data to give more information about system performance. In particular, we estimated the following quantities: the rate constant for binding of *M*_*x*_*L* to *T*_*xy*_ (or the displacement of *L* from *M*_*x*_ by *T*_*xy*_); the rate constant for spontaneous leak reactions between *M*_*x*_*L* and *N*_*y*_, the rate constant of the HMSD substep for *M*_1_*N*_3_ and the initial reaction rate for catalytic dimerization (TOF) for all templates used.

All fits were performed with MATLAB R2019a Optimization Toolbox. Supplementary Note 6 contains a detailed description of the fitting procedures, with the resultant fits tabulated in Supplementary Tables 13-18 and Illustrated in Supplementary Figures 10-18.

## Acknowledgement

TEO was supported by a Royal Society University Research Fellowship, JCG by a Royal Society PhD studentship, and G.-B.S. by a Royal Academy of Engineering Chair in Emerging Technology for Engineering Biology (CiET1819*\*5). This work is part of a project that has received funding from the European Research Council (ERC) under the European Union’s Horizon 2020 research and innovation programme (Grant agreement 851910). This research was supported by EPSRC grant EP/P02596X/1.

## Supporting Information Available

Further results, and information on experimental methods and data processing, can be obtained from the Supplementary Information. Raw data, figures, and the scripts used to process that data and generate the results plotted here, are available to download from Zenodo (https://doi.org/10.5281/zenodo.8256587).

