## Supplementary Notes for "Information propagation through enzyme-free catalytic templating of DNA dimerization with weak product inhibition"

### Supplementary tables


1 Template strands ($T_{xy}$) . . . 8

2 Monomer M ($M_x$) and its lock strands ($L_x$) . . . 9

3 Monomer N ($N_y$) strands . . . 9

4 Strands used in each figure of this publication . . . 10

5 Calibration matrix 1: multi-channel coefficient of fluorescence (fluorescence emission per nM) of the species present in sequence-specific copying experiments at 25° C . . . 15

6 Calibration matrix 2: multi-channel coefficient of fluorescence (fluorescence emission per nM) of the species present in sequence-specific copying experiments at 25° C . . . 16

7 Detailed experimental procedure for $M_x$-binding characterisation . . . 20

8 Detailed experimental procedure for 1-fluorescence channel monomers leak reaction characterisation . . . 21

9 Detailed experimental procedure to measure dimerization kinetics . . . 23

10 Detailed experimental procedure to test product inhibition . . . 24

11 Detailed experimental procedure for dimerization kinetics with labeled $N_y$. . . 26

12 Detailed experimental procedure for sequence-specific copying by templated dimerization. . . . 27

13 Fitted TMSD rate constant $k_t$. . . . 42

14 Fitted leak reaction rate $k_{\text{leak}}$. . . . 45

15 Fitted $k_h$. . . . 47

16 Initial rates of monomer turnover by $T_{13}$ in the presence of $M_1N_3$ (10 nM-scale). 50

17 Initial rates of monomer turnover by $T_{13}$ in the presence of $M_1N_3$ (100 nM-scale). 52

18 Initial rates for monomer turnover by $T_{xy}$ used during specific dimerization assays. . . . 54

19 Templating precision of first specific-dimerization iteration. . . . 69

20 Templating precision of second specific-dimerization iteration. . . . . . . . . . 69

21 Templating precision of third specific-dimerization iteration. . . . . . . . . . . 69

### Supplementary figures


1 Calibration: $M_1$ strand alone (Alexa Fluor® 488) at 25° C. Gain 1261 . . . . 13
2 Calibration: $M_1L$ complex. (Quenched Alexa Fluor® 488) at 25° C. Detector Gain 1261 . . . . . . . . . . . . . . . . . . . . . . . . . . . . . . . . . . 13
3 Calibration: $T_{13}M_1$ complex (Alexa Fluor® 488) at 25° C. Detector Gain 1261 14
4 Calibration: $M_1N_3$ complex (Alexa Fluor® 488) at 25° C. Detector Gain 1261 14
5 Example of $M_xL$ binding characterisation. . . . . . . . . . . . . . . . . . . . 20
6 Example of dimerization experiments with $N_3$ (single fluorescent channel). . 23
7 Example of dimerization experiments with labeled $N_y$ (multichannel fluorescence experiment). . . . . . . . . . . . . . . . . . . . . . . . . . . . . . . 26
8 Raw data showing the kinetics of the sequence-specific copying by templated dimerization. . . . . . . . . . . . . . . . . . . . . . . . . . . . . . . . . 28
9 Example of the result of fluorescence data processing for single fluorophore experiments. . . . . . . . . . . . . . . . . . . . . . . . . . . . . . . . . 34
10 Fits of $k_t$ for different primary toehold lengths $t$. . . . . . . . . . . . . . . . 43
11 Fits of $k_t$ for different competing templates. . . . . . . . . . . . . . . . . . 44
12 Fits of $k_{\mathrm{leak}}$ for different combination of monomers. . . . . . . . . . . . . 46
13 Fits of $k_h$ for different handhold lengths. . . . . . . . . . . . . . . . . . . . 47
14 Primary toehold length doesn't affect the HMSD kinetics. . . . . . . . . . . . 48
15 Kinetics of 10 nM-scale inhibition tests. . . . . . . . . . . . . . . . . . . . . 51
16 Kinetics of 100 nM-scale inhibition tests. . . . . . . . . . . . . . . . . . . . 53
17 Kinetics of the $T_{xy}$-mediated reaction in isolation. . . . . . . . . . . . . . . 55
18 Linear fits of sequenze-specific dimerization templates' initial rates to obtain their TOF. . . . . . . . . . . . . . . . . . . . . . . . . . . . . . . . . . . 56
19 Outline design of a system that exploits HMSD to copy a longer template . . 58
20 Performance of the clamp-free version of the dimerization system . . . . . . 61
21 Exploring reaction kinetics for the original version of the displacement domain 62

22 Kinetics of first iteration of sequence-specific dimerization experiment . . . . 64
23 PAGE of first iteration of sequence-specific dimerization experiment . . . . . 65
24 Kinetics of second iteration of sequence-specific dimerization experiment . . 66
25 PAGE of second iteration of sequence-specific dimerization experiment . . . 67
26 Negative controls for specific dimerization experiments. . . . . . . . . . . . . 67
27 Template-mediated interconversion of products . . . . . . . . . . . . . . . . . . 71
28 Secondary toehold mediated TMSD reaction without handhold complmentarity 73
29 Evidence of the effective irreversibility of the displacement of $L$ from $M_1L$ by $T_{13}$ . . . . . . . . . . . . . . . . . . . . . . . . . . . . . . . . . . . . . 74

### 1 Supplementary Note 1: DNA sequences

All sequences used are collected in Supplementary Tables 1 to 3, and the strands used for each figure are listed in Supplementary Table 4. Strands are named after the sequence and length of their primary toehold ($t^1$) and handhold domain (h). Template strands ($T_{xy}$ ($u$t/$v$h) can bind by the $t^1$ domain to the pool of monomer $M$ duplexes ($M_xL$), and by the h domain the pool of complementary monomer $N$ strands ($N_y$).

$T_{xy}$, $M_x$ and $N_y$ strands have, by default, toehold domains of length 6nt and handhold domains of length 8nt. To test variants of t and h, we used the following:

- To test toeholds longer than 6, we used $M_1$ with toeholds of length 8 and $T_{13}$ with with toeholds of length 7 and 8.
- To test handholds longer than 8, we used $N_3$ with handholds of length 10 and $T_{13}$ with with handholds of length 9 and 10.
- To test shorter handholds or toeholds, we used $M_x$ and $N_y$ strands with toehold domains of length 6nt and handhold domains of length 8nt, and truncated versions of $T_{xy}$.

Sequences are colored by domain using the following colors:

**Red**: Mismatches whose repair drives the dimerisation reaction.

**Grey**: $L_x$ dimerization domain that binds to $M_x$. Complementary to blue. The same sequence as orange and green, but contains two mismatches with $M_x$.

**Orange**: $T_{xy}$ dimerization domain that binds to $M_x$, and is displaced during HMSD by $N_y$'s green dimerization domain. Same sequence as gray and green, but contains one mismatch with $M_x$.

**Green**: $N_y$ dimerization domain that binds to $M_x$ in order to create the dimer. Same sequence as gray and orange, but it does not contain any mismatch with $M_x$.

**Blue**: $M_x$ strands' dimerization domain. Complementary to gray, orange, and green.

**Gold**: Handhold domain of $T_{xy}$ strands and handhold-complementary domains in $N_y$ strands. Variable length and sequence in $T_{xy}$ and $N_y$ strands .

**Purple**: Toehold recognition domain of $T_{xy}$ strands and recognition-complementary toehold domains in $M_x$ strands. These domains constitute the primary toehold. Variable length and sequence in $T_{xy}$ and $M_x$ strands.

**Light blue**: Secondary toehold ($t^2$) domain and its complement. Always the same 2 nt sequence. Present in $M_x$; the complement is present in $L_x$ and $N_y$ strands

**Black:** 2 nt spacer domains between orange and gold domains in $T_{xy}$ strands; poly-T spacers; and clamp nucleotides in $M_x$, $L_x$ and $T_{xy}$ strands.

Supplementary Table 1: Template strands ($T_{xy}$).

| | Sequence | Size (nt) |
|---|---|---|
| $T_{11}$ | 5' - GTT TAG **G**CG TTT TAT CTT CA**C** TTC CAT CCA TTC CAG TTC CAT TAG CGT TGA TGA GGA - 3' | 57 |
| $T_{12}$ | 5' - GTT TAG **G**CG TTT TAT CTT CA**C** TTC CAT CCA TTC CAG TTC CAT TAG CGT TGA GTA CTG - 3' | 57 |
| $T_{13}$ (6t/8h) | 5' - GTT TAG **G**CG TTT TAT CTT CA**C** TTC CAT CCA TTC CAG TTC CAT TAG CGT TGT GGT AAG - 3' | 57 |
| $T_{21}$ | 5' - TTG TCA **G**CG TTT TAT CTT CA**C** TTC CAT CCA TTC CAG TTC CAT TAG CGT TGA TGA GGA - 3' | 57 |
| $T_{22}$ | 5' - TTG TCA **G**CG TTT TAT CTT CA**C** TTC CAT CCA TTC CAG TTC CAT TAG CGT TGA GTA CTG - 3' | 57 |
| $T_{23}$ | 5' - TTG TCA **G**CG TTT TAT CTT CA**C** TTC CAT CCA TTC CAG TTC CAT TAG CGT TGT GGT AAG - 3' | 57 |
| $T_{31}$ | 5' - ATT CTT **G**CG TTT TAT CTT CA**C** TTC CAT CCA TTC CAG TTC CAT TAG CGT TGA TGA GGA - 3' | 57 |
| $T_{32}$ | 5' - ATT CTT **G**CG TTT TAT CTT CA**C** TTC CAT CCA TTC CAG TTC CAT TAG CGT TGA GTA CTG - 3' | 57 |
| $T_{33}$ | 5' - ATT CTT **G**CG TTT TAT CTT CA**C** TTC CAT CCA TTC CAG TTC CAT TAG CGT TGT GGT AAG - 3' | 57 |
| $T_{13}$ (4t/6h) | 5' - TTA G**G**C GTT TTA TCT TCA **C**TT CCA TCC ATT CCA GTT CCA TTA GCG TTG TGG TA - 3' | 53 |
| $T_{13}$ (4t/7h) | 5' - TTA G**G**C GTT TTA TCT TCA **C**TT CCA TCC ATT CCA GTT CCA TTA GCG TTG TGG TAA - 3' | 54 |
| $T_{13}$ (4t/8h) | 5' - TTA G**G**C GTT TTA TCT TCA **C**TT CCA TCC ATT CCA GTT CCA TTA GCG TTG TGG TAA G - 3' | 55 |
| $T_{13}$ (4t/9h) | 5' - TTA G**G**C GTT TTA TCT TCA **C**TT CCA TCC ATT CCA GTT CCA TTA GCG TTG TGG TAA GA - 3' | 56 |
| $T_{13}$ (4t/10h) | 5' - TTA G**G**C GTT TTA TCT TCA **C**TT CCA TCC ATT CCA GTT CCA TTA GCG TTG TGG TAA GAG - 3' | 57 |
| $T_{13}$ (5t/6h) | 5' - TTT AG**G** CGT TTT ATC TTC A**C**T TCC ATC CAT TCC AGT TCC ATT AGC GTT GTG GTA - 3' | 54 |
| $T_{13}$ (5t/7h) | 5' - TTT AG**G** CGT TTT ATC TTC A**C**T TCC ATC CAT TCC AGT TCC ATT AGC GTT GTG GTA A - 3' | 55 |
| $T_{13}$ (5t/8h) | 5' - TTT AG**G** CGT TTT ATC TTC A**C**T TCC ATC CAT TCC AGT TCC ATT AGC GTT GTG GTA AG - 3' | 56 |
| $T_{13}$ (5t/9h) | 5' - TTT AG**G** CGT TTT ATC TTC A**C**T TCC ATC CAT TCC AGT TCC ATT AGC GTT GTG GTA AGA - 3' | 57 |
| $T_{13}$ (5t/10h) | 5' - TTT AG**G** CGT TTT ATC TTC A**C**T TCC ATC CAT TCC AGT TCC ATT AGC GTT GTG GTA AGA G - 3' | 58 |
| $T_{13}$ (6t/6h) | 5' - GTT TAG **G**CG TTT TAT CTT CA**C** TTC CAT CCA TTC CAG TTC CAT TAG CGT TGT GGT A - 3' | 55 |
| $T_{13}$ (6t/7h) | 5' - GTT TAG **G**CG TTT TAT CTT CA**C** TTC CAT CCA TTC CAG TTC CAT TAG CGT TGT GGT AA - 3' | 56 |
| $T_{13}$ (6t/9h) | 5' - GTT TAG **G**CG TTT TAT CTT CA**C** TTC CAT CCA TTC CAG TTC CAT TAG CGT TGT GGT AAG A - 3' | 57 |
| $T_{13}$ (6t/10h) | 5' - GTT TAG **G**CG TTT TAT CTT CA**C** TTC CAT CCA TTC CAG TTC CAT TAG CGT TGT GGT AAG AG - 3' | 58 |
| $T_{13}$ (7t/6h) | 5' - TGT TTA G**G**C GTT TTA TCT TCA **C**TT CCA TCC ATT CCA GTT CCA TTA GCG TTG TGG TA - 3' | 56 |
| $T_{13}$ (7t/7h) | 5' - TGT TTA G**G**C GTT TTA TCT TCA **C**TT CCA TCC ATT CCA GTT CCA TTA GCG TTG TGG TAA - 3' | 57 |
| $T_{13}$ (7t/8h) | 5' - TGT TTA G**G**C GTT TTA TCT TCA **C**TT CCA TCC ATT CCA GTT CCA TTA GCG TTG TGG TAA G - 3' | 58 |
| $T_{13}$ (7t/9h) | 5' - TGT TTA G**G**C GTT TTA TCT TCA **C**TT CCA TCC ATT CCA GTT CCA TTA GCG TTG TGG TAA GA - 3' | 59 |
| $T_{13}$ (7t/10h) | 5' - TGT TTA G**G**C GTT TTA TCT TCA **C**TT CCA TCC ATT CCA GTT CCA TTA GCG TTG TGG TAA GAG- 3' | 60 |
| $T_{13}$ (8t/6h) | 5' - TTG TTT AG**G** CGT TTT ATC TTC A**C**T TCC ATC CAT TCC AGT TCC ATT AGC GTT GTG GTA - 3' | 57 |
| $T_{13}$ (8t/7h) | 5' - TTG TTT AG**G** CGT TTT ATC TTC A**C**T TCC ATC CAT TCC AGT TCC ATT AGC GTT GTG GTA A - 3' | 58 |
| $T_{13}$ (8t/8h) | 5' - TTG TTT AG**G** CGT TTT ATC TTC A**C**T TCC ATC CAT TCC AGT TCC ATT AGC GTT GTG GTA AG - 3' | 59 |
| $T_{13}$ (8t/9h) | 5' - TTG TTT AG**G** CGT TTT ATC TTC A**C**T TCC ATC CAT TCC AGT TCC ATT AGC GTT GTG GTA AGA - 3' | 60 |
| $T_{13}$ (8t/10h) | 5' - TTG TTT AG**G** CGT TTT ATC TTC A**C**T TCC ATC CAT TCC AGT TCC ATT AGC GTT GTG GTA AGA G - 3' | 61 |
| $T_{13}$ v.1 | 5' - GGA AAC A CG CAT CAT A **C** T CAA GTC AAA GTC AAG TCA TAT TCA GGG - 3' | 45 |

Supplementary Table 2: Monomer M ($M_x$) and its lock strands ($L_x$).

| | Sequence | Size (nt) |
|---|---|---|
| $M_1$ | 5' - **AlexaFluor®488** – ttt **C**GT CGC TAA TGG AAC TGG AAT GGA TGG AAC TGA AGA TAA AAC G**C**C TAA AC - 3' | 53 |
| $M_2$ | 5' - **AlexaFluor®546** – ttt ttt ttt ttt **C**GT CGC TAA TGG AAC TGG AAT GGA TGG AAC TGA AGA TAA AAC G**C**T GAC AA - 3' | 62 |
| $M_3$ | 5' - **AlexaFluor®647** – ttt ttt **C**GT CGC TAA TGG AAC TGG AAT GGA TGG AAC TGA AGA TAA AAC G**C**A AGA AT - 3' | 56 |
| $M_1$ (Unlab.) | 5' - ttt **C**GT CGC TAA TGG AAC TGG AAT GGA TGG AAC TGA AGA TAA AAC G**C**C TAA AC - 3' | 53 |
| $M_1$ (8t) | 5' - **AlexaFluor®488** – ttt **C**GT CGC TAA TGG AAC TGG AAT GGA TGG AAC TGA AGA TAA AAC G **C**C TAA ACA A - 3' | 55 |
| $M_1$ (8t Unlab.) | 5' - ttt **C**GT CGC TAA TGG AAC TGG AAT GGA TGG AAC TGA AGA TAA AAC G**C**C TAA ACA A - 3' | 55 |
| $M_1$ v.1 | 5' - **AlexaFluor®488** – ttt CCG ACT TGA CTT TGA CTT GAC TAT GAT GCGTGT TTC - 3' | 38 |
| $L_1$ | 5' - **G**CG TTT TAT CTT CA**C** TTC CAT CCA TTC CA**C** TTC CAT TAG CGA C**G** ttt – **IowaBlack®FQ** - 3' | 47 |
| $L_2$ | 5' - **G**CG TTT TAT CTT CA**C** TTC CAT CCA TTC CA**C** TTC CAT TAG CGA C**G** ttt ttt ttt ttt – **BlackHoleQuencher®2** - 3' | 53 |
| $L_3$ | 5' - **G**CG TTT TAT CTT CA**C** TTC CAT CCA TTC CAC TTC CAT TAG CGA C**G** ttt ttt – **IowaBlack®RQ** - 3' | 50 |
| $L_1$ v.1 | 5' - CGC ATC ATA**C** TC AAG TCA AA**C** TCA AGT CGG ttt – **BlackHoleQuencher®1** - 3' | 33 |

Supplementary Table 3: Monomer N ($N_y$) strands.

| | Sequence | Size (nt) |
|---|---|---|
| $N_1$ | 5' - TCC TCA TCC GTT TTA TCT TCA GTT CCA TCC ATT CCA GTT CCA TTA GCG AC ttt ttt ttt – **AlexaFluor®647** - 3' | 59 |
| $N_2$ | 5' - CAG TAC TCC GTT TTA TCT TCA GTT CCA TCC ATT CCA GTT CCA TTA GCG AC ttt – **AlexaFluor®546** - 3' | 53 |
| $N_3$ | 5' - CTT ACC ACC GTT TTA TCT TCA GTT CCA TCC ATT CCA GTT CCA TTA GCG AC - 3' | 50 |
| $N_3$ (10h) | 5' - CTC TTA CCA CCG TTT TAT CTT CAG TTC CAT CCA TTC CAG TTC CAT TAG CGA C - 3' | 52 |
| $N_3$ v.1 | 5' - CCC TGA ATC GCA TCA TAG TCA AGT CAA AGT CAA GTCGG - 3' | 38 |
| $N$ v.1 (0h) | 5' - C GCA TCA TAG TCA AGT CAA AGT CAA GTCGG - 3' | 30 |

**Supplementary Table 4: Strands used in each figure of this publication.**

| Figure | Strands used |
|---|---|
| **2** | $T_{13}$ (4-8t/6-10h), $M_1$, $M_1$ (8t), $L_1$, $N_3$, $N_3$ (10h) |
| **3a** | $T_{13}$ (5-7t/8-10h), $M_1$, $M_1$ (Unlab.), $M_1$ (8t), $M_1$ (8t Unlab.), $L_1$, $N_3$, $N_3$ (10h) |
| **3c** | $T_{13}$ (6t/8h), $M_1$, $L_1$, $N_3$ |
| **4b** | $T_{11}$, $T_{12}$, $T_{13}$ (6t/8h), $T_{21}$, $T_{22}$, $T_{23}$, $T_{31}$, $T_{32}$, $T_{33}$, $M_1$, $M_2$, $M_3$, $L_1$, $L_2$, $L_3$, $N_1$, $N_2$, $N_3$ |
| **5** | $T_{11}$, $T_{12}$, $T_{13}$ (6t/8h), $T_{21}$, $T_{22}$, $T_{23}$, $T_{31}$, $T_{32}$, $T_{33}$, $M_1$, $M_2$, $M_3$, $L_1$, $L_2$, $L_3$, $N_1$, $N_2$, $N_3$ |
| **Sup.1** | $M_1$ |
| **Sup.2** | $M_1$, $L_1$ |
| **Sup.3** | $M_1$, $L_1$, $T_1$3(6t/8h) |
| **Sup.4** | $M_1$, $L_1$, $T_1$3(6t/8h), $N_3$ |
| **Sup.5** | $M_1$, $L_1$, $T_1$2 |
| **Sup.6** | $M_2$, $L_2$, $T_2$3, $N_3$ |
| **Sup.7** | $M_1$, $L_1$, $T_1$2, $N_2$ |
| **Sup.8** | $T_{11}$, $T_{12}$, $T_{13}$ (6t/8h), $T_{21}$, $T_{22}$, $T_{23}$, $T_{31}$, $T_{32}$, $T_{33}$, $M_1$, $M_2$, $M_3$, $L_1$, $L_2$, $L_3$, $N_1$, $N_2$, $N_3$ |
| **Sup.9** | $M_3$, $L_3$, $T_{33}$, $N_3$ |
| **Sup.10** | $M_1$, $L_1$, $T_{13}$(4t/8h), $T_{13}$(5t/8h), $T_{13}$(6t/8h), $T_{13}$(7t/8h), $T_{13}$(8t/8h) |
| **Sup.11** | $M_1$, $M_2$, $M_3$, $L_1$, $L_2$, $L_3$, $T_{11}$, $T_{12}$, $T_{13}$(6t/8h), $T_{21}$, $T_{22}$, $T_{23}$, $T_{31}$, $T_{32}$, $T_{33}$ |
| **Sup.12** | $M_1$, $M_2$, $M_3$, $L_1$, $L_2$, $L_3$, $N_1$, $N_2$, $N_3$ |
| **Sup.13** | $M_1$, $L_1$, $N_3$, $T_{13}$(6t/6h), $T_{13}$(6t/7h), $T_{13}$(6t/9h), $T_{13}$(6t/10h) |
| **Sup.14** | $M_1$, $L_1$, $N_3$, $T_{13}$(4t/8h), $T_{13}$(5t/8h), $T_{13}$(6t/8h), $T_{13}$(7t/8h), $T_{13}$(8t/8h) |
| **Sup.15** | $T_{13}$ (4-8t/6-10h), $M_1$, $M_1$ (Unlab.), $M_1$ (8t), $M_1$ (8t Unlab.), $L_1$, $N_3$, $N_3$ (10h) |
| **Sup.16** | $T_{13}$ (6t/8h), $M_1$, $M_1$ (Unlab.), $L_1$, $N_3$ |
| **Sup.17** | $T_{11}$, $T_{12}$, $T_{13}$ (6t/8h), $T_{21}$, $T_{22}$, $T_{23}$, $T_{31}$, $T_{32}$, $T_{33}$, $M_1$, $M_2$, $M_3$, $L_1$, $L_2$, $L_3$, $N_1$, $N_2$, $N_3$ |
| **Sup.20** | $T_{13}$ v.1, $M_1$ v.1, $L_1$ v.1, $N_3$ v.1 |
| **Sup.21a** | $T_{13}$ v.1, $M_1$ v.1, $L_1$ v.1 |
| **Sup.21b** | $T_{13}$, $T_{13}$ v.1, $M_1$, $M_1$ v.1, $L_1$, $L_1$ v.1, $N_3$, $N_3$ v.1, $N_3$ v.1 (0h) |
| **Sup.22** | $T_{11}$, $T_{12}$, $T_{13}$ (6t/8h), $T_{21}$, $T_{22}$, $T_{23}$, $T_{31}$, $T_{32}$, $T_{33}$, $M_1$, $M_2$, $M_3$, $L_1$, $L_2$, $L_3$, $N_1$, $N_2$, $N_3$ |
| **Sup.23** | $T_{11}$, $T_{12}$, $T_{13}$ (6t/8h), $T_{21}$, $T_{22}$, $T_{23}$, $T_{31}$, $T_{32}$, $T_{33}$, $M_1$, $M_2$, $M_3$, $L_1$, $L_2$, $L_3$, $N_1$, $N_2$, $N_3$ |
| **Sup.24** | $T_{11}$, $T_{12}$, $T_{13}$ (6t/8h), $T_{21}$, $T_{22}$, $T_{23}$, $T_{31}$, $T_{32}$, $T_{33}$, $M_1$, $M_2$, $M_3$, $L_1$, $L_2$, $L_3$, $N_1$, $N_2$, $N_3$ |
| **Sup.25** | $T_{11}$, $T_{12}$, $T_{13}$ (6t/8h), $T_{21}$, $T_{22}$, $T_{23}$, $T_{31}$, $T_{32}$, $T_{33}$, $M_1$, $M_2$, $M_3$, $L_1$, $L_2$, $L_3$, $N_1$, $N_2$, $N_3$ |
| **Sup.26** | $M_1$, $M_2$, $M_3$, $L_1$, $L_2$, $L_3$, $N_1$, $N_2$, $N_3$ |
| **Sup.27a** | $T_{11}$, $M_1$, $L_1$, $N_1$, |
| **Sup.27b** | $T_{12}$, $T_{33}$, $M_1$, $N_1$, $N_3$ |
| **Sup.28a** | $T_{12}$,$T_{22}$, $T_{32}$, $M_1$, $M_2$, $M_3$, $L_1$, $L_2$, $L_3$, $N_3$ |
| **Sup.28b** | $T_{12}$, $M_1$, $L_1$, $N_1$ |
| **Sup.29** | $T_{13}$ (6t/8h), $M_1$, $L_1$, $N_3$ |

### 2 Supplementary Note 2: Fluorescent species calibration

This section describes the methods used to obtain calibration curves for converting the raw fluorescence data from the experiments into concentration estimates. It also presents the results of calibration.

In our experiments, we measure fluorescence in a plate reader from the bottom of the plate. Reactions are triggered injecting liquid into the plate, meaning that the volume changes (by up to 50% over the full course of the measurement). Typically, the change in fluorescence measured in response to such a dilution is small, because the path length of fluid in the illumination/detection region increases to compensate for a decrease in concentration.[1] Moreover, adjustment of the focusing height of the plate reader can provide fine adjustment to the relative fluorescence before and after dilution. We selected a focusing height such that the dilution didn't produce a significant fluorescence difference between the volumes of 150 µL and 200 µL most commonly used in our experiments. As a result, the fluorescence signals effectively report the molecular count of certain species in the well. For convenience, however, we will refer to these molecular counts through a multi-step experiment in terms of their concentration as if they were solvated in a 200 µL volume (the volume at which reaction kinetics were measured).

For our experimental setup, the signal measured by the plate reader is assumed to be a linear function of this effective molecular concentration of fluorescent species. The constant term is assumed to be a background offset equivalent to the fluorescence of a buffer-only control. Therefore, after subtracting the median fluorescence of the negative control from all values, a weighted linear regression model was used to fit a proportional expression for the median fluorescence $F$ of each well as a function of molecular concentration $x$, $F(x) = ax$. Here, $a$ is the fluorescent count per nM of the labelled species for the measurement conditions; we describe this quantity as a "coefficient of fluorescence". The only exception to the above was the weakly-fluorescent species $M_1L$, which required an offset to account for a background signal.

Fitting was performed with the MATLAB R2019a Optimization Toolbox.[2] The residuals were weighted with the inverse of the fluorescence reading to account for the Poisson distribution of the photomultiplier counts .[3] The regression coefficient value is reported with 95% confidence bounds. Fitting results and their residuals for the fluorescence of $M_1$ in various complexes are shown in Supplementary Figures 1 to 4.

#### 2.1 Single channel calibration of Alexa Fluor® 488 for complexes involved in $M_1N_3$ formation

To a first approximation, the $M_1L$ complex has very low fluorescence, and all other complexes containing $M_1$ have high fluorescence. However, the fluorescence of $M_1L$ is not zero, and other complexes of $M_1$ show variation in the fluorescence signal due to changes in the sequence environment of the fluorophores. Calibration curves were used to estimate the signal variation between the different hybridization states of $M_1$, and the fluorescence of $M_1L$.

##### 2.1.1 Protocol

1. Three $M_1$ sets of serial dilutions with concentrations ranging from 0 to 150 nM were prepared in a volume of 150 µL. Fluorescence was measured and fitted using linear regression (Figure 1), giving the coefficient of fluorescence of $M_1$.

2. One of the $M_1$ sets of serial dilutions was brought to a volume of 195 µL, with an injection of $L_1$ sufficient to achieve a $[L_1] = 500$ nM. The fluorescence data (Figure 2) allows for the calculation of the coefficient of fluorescence of $M_1L$.

3. That same serial dilution was brought to a volume of 200 µL with an injection of $T_{13}$ sufficient to give $[T_{13}] = 200$ nM. Another set of serial dilutions containing just $M_1$ was brought to the same volume using $T_{13}$ ($[T_{13}] = 200$ nM) for comparison and to

demonstrate that the displacement of $L_1$ from $M_1L$ by the added $T_{13}$ was complete. The fluorescence data of both replicas (Figure 3) allows for the calculation of the coefficient of fluorescence of $M_1T_{13}$.

4. All three sets of serial dilutions were then brought to 205 µL with an injection of $N_3$, sufficient to give a concentration of $[N_3] = 500$ nM. The three curves (Figure 4) were used to calculate the coefficient of fluorescence of $M_1N_3$. Their consistency is indicative of the fact that a large excess of $N_3$ is sufficient to convert all $M_1$ into $M_1N_3$ complexes, regardless of the presence of excess concentrations of $L$ or $T_{13}$.

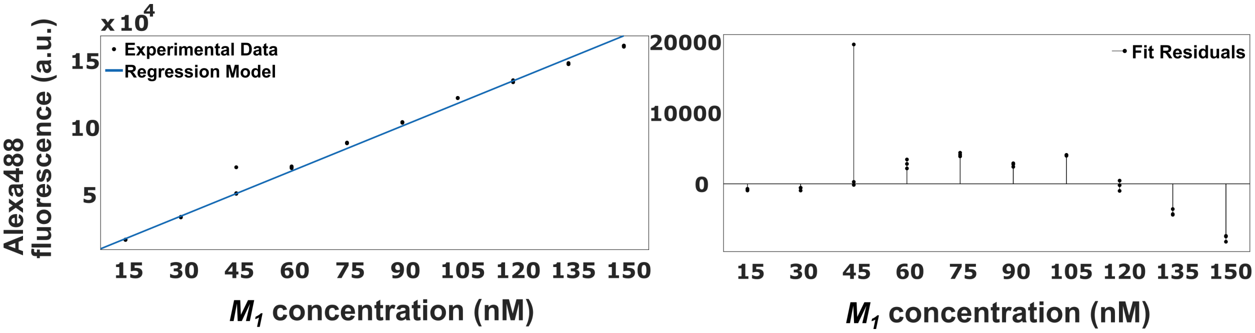


**Supplementary Figure 1: Calibration: $M_1$ strand (Alexa Fluor® 488) at 25° C. Detector Gain 1261.** 3 replicas per point. Left: Regression line. Right: Fit residuals. a = 1123 fluo a.u. nM$^{-1}$ (1106, 1140). Goodness of fit: SSE = 4425. R-square = 0.9959. RMSE = 12.36.

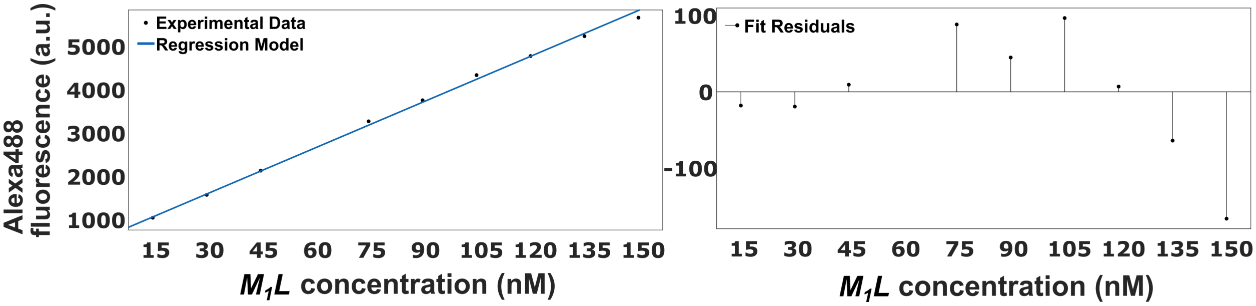


**Supplementary Figure 2: Calibration: $M_1L$ complex (Quenched Alexa Fluor® 488) at 25° C. Detector Gain 1261.** Added $L$ in excess. Only one replica per point. Left: Regression line. Right: Fit residuals. a = 35.24 fluo a.u. nM$^{-1}$ (33.86, 36.63). The calibration line had a non-negligible background (b = 550.2 (449.6, 650.8)). This term is probably due to wrongly annealed duplexes, as it doesn't appear in other calibrations using thermocycler-annealed strands. Therefore, it was ignored in subsequent analysis. Goodness of fit: SSE = 19.28. R-square = 0.9977. Adjusted R-square = 0.9974. RMSE = 1.552.

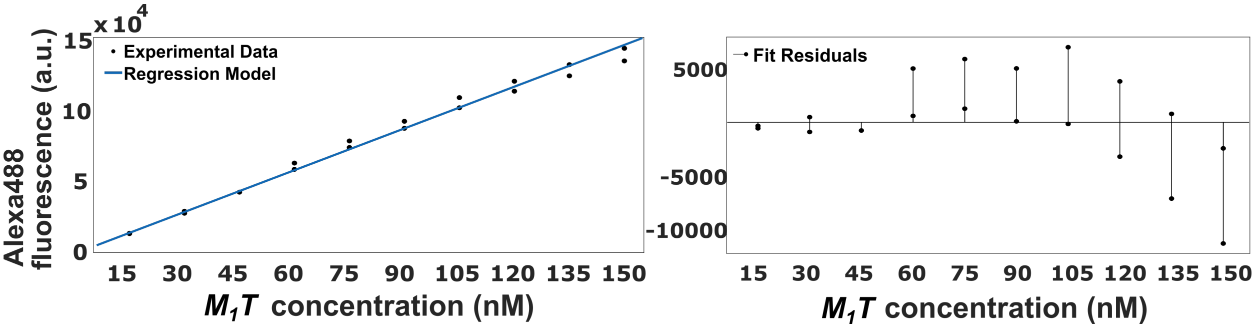


**Supplementary Figure 3: Calibration: $T_{13}M_1$ complex (Alexa Fluor® 488) at 25° C. Detector Gain 1261.** Triggered $M_1L$ with excess of $T_{13}$. Only one replica per point. Left: Regression line. Right: Fit residuals. a = 979 fluo a.u. nM$^{-1}$ (957.6, 1000). Goodness of fit: SSE = 3241. R-square = 0.995. RMSE = 13.06.

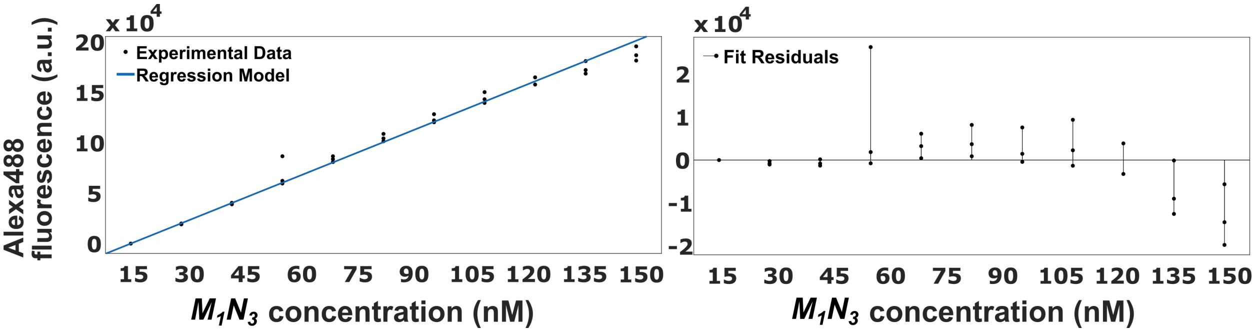


**Supplementary Figure 4: Calibration: $M_1N_3$ complex (Alexa Fluor® 488) at 25° C. Detector Gain 1261.** Three replicas; consistency shows that saturating $N_3$ is able to bind to all $M_1$. Left: Regression line. Right: Fit residuals. a = 1342 fluo a.u. nM$^{-1}$ (1317, 1367). Goodness of fit: SSE = 7893. R-square = 0.9935. RMSE = 16.5.

#### Multi-channel fluorescence calibration

Sequence-specific copying experiments involved the formation of several $M_xN_y$ dimers. Each $M_xN_y$ had a different combination of fluorophores that produced a specific fluorescence emission that was measured in six different fluorescent channels. These species had a non-negligible bleed-through in the different fluorescence channels that had to be quantified for a correct deconvolution of the measured fluorescence signal. The data collected in Supplementary Tables 5 and 6 shows, for monomers and dimer products, the coefficient of fluorescence of each species in each channel. Each entry into this calibration matrix was obtained from a single set of three concentrations of the relevant species (25, 50, and 100 nM); these data were fit assuming that fluorescence was proportional to the molecular concentration after accounting for background. We report two versions of the calibration matrix in Suppleementary Tables 5 and 6, since experimetns were performed using two different settings on the plate reader.

These experiments additionally confirmed two things. First, the quenching efficiency of each $M_xL$ complex: $M_1L_1 = 97.1 \pm 0.3$ %; $M_2L_2 = 89.3 \pm 0.7$ %; $M_3L_3 = 89.4 \pm 1$%. Second, the strands labeled with Alexa Fluor® 546 or 647 had essentially identical fluorescence in any complex that contained no other fluorophores (to within approximately 2%). In other words, $M_3N_3$ and $N_1$ could be described with a single set of parameters, as could $M_2N_3$ and $N_2$. The data presented in Supplementary Tables 5 and 6 for these complexes uses an average over these pairs.

**Supplementary Table 5: Calibration matrix 1: multi-channel coefficient of fluorescence of the species present in sequence-specific copying experiments at 25° C.** Calculated values of the coefficient of fluorescence of $M_xL$, $M_xN_y$ and $N_y$ ($a$ if $f(x) = ax$, where $f$ is fluorescence and $x$ is concentration) for 6 different fluorescence channels. These values were used for all multichannel experiments, except iteration 1 of the sequence-specific dimerization experiments. Unlabelled $N_3$ showed no fluorescence above the background. **Bold:** Main channel used to quantify the species in each row. Blue = Alexa Fluor® 488 (EX: 488/14 nm; EM: 535/30 nm), Gain = 1261; Green = Alexa Fluor® 546 (EX: 540/20 nm; EM: 590/30 nm) Gain = 1304; Red = Alexa Fluor® 647 (EX: 625/30 nm; EM: 680/30 nm) Gain = 1896; B/G = FRET signal of Alexa Fluor® 488 and 546 (EX: 488/14 nm; EM: 590/30 nm), Gain = 1461; G/R = FRET signal of Alexa Fluor® 488 and 647 (EX: 488/14 nm; EM: 670/30 nm), Gain = 2259 and B/R = FRET signal of Alexa Fluor® 546 and 647 (EX: 540/20 nm; EM: 680/30 nm), Gain = 2257. Fitted data is collected in the data repository accompanying this publication.

| Species | Blue | Green | Red | B/G | G/R | B/R |
|---|---|---|---|---|---|---|
| $M_1N_1$ | 410 | 0.71 | 625 | 88 | 198 | **1363** |
| $M_1N_2$ | 200 | 679 | 0.17 | **1229** | 131 | 251 |
| $M_1N_3$ | **1332** | 1.19 | 0.05 | 287 | 0.89 | 40 |
| $M_2N_1$ | 4.1 | 260 | 684 | 63 | **1289** | 246 |
| $M_2N_2$ | 10 | **965** | 0.04 | 235 | 193 | 54 |
| $M_2N_3$ **or** $N_2$ | 11 | **924** | 0.003 | 224 | 174 | 49 |
| $M_3N_1$ | 1.3 | 0.28 | **1020** | 0.54 | 310 | 48 |
| $M_3N_2$ | 1.9 | 88 | 599 | 23 | **1327** | 250 |
| $M_3N_3$ **or** $N_1$ | 0.72 | 0.29 | **680** | 0.35 | 200 | 29 |
| $M_1L_1$ | **39** | 0.08 | 0.11 | 8.8 | 0.23 | 1.4 |
| $M_2L_2$ | 1.5 | **97** | 0.03 | 24 | 19 | 5.3 |
| $M_3L_3$ | 0.52 | 0.05 | **73** | 0.15 | 23 | 3.7 |

**Supplementary Table 6: Calibration matrix 2: multi-channel coefficient of fluorescence (fluorescence emission per nM) of the species present in sequence-specific copying experiments at 25° C.** Calculated values of the coefficient of fluorescence of $M_xL$, $M_xN_y$ and $N_y$ ($a$ if $F(x) = ax$, where $F$ is fluorescence and $x$ is concentration) for 6 different fluorescence channels. This calibration is used for the 1$^{st}$ iteration of the sequence-specific dimerization experiment (Supplementary Figure 22. Unlabelled $N_3$ showed no fluorescence above the background. **Bold:** Main channel used to quantify the species in each row. Blue = Alexa Fluor® 488 (EX: 488/14 nm; EM: 535/30 nm), Gain = 1508; Green = Alexa Fluor® 546 (EX: 540/20 nm; EM: 590/30 nm) Gain = 1552; Red = Alexa Fluor® 647 (EX: 625/30 nm; EM: 680/30 nm) Gain = 2169; B/G = FRET signal of Alexa Fluor® 488 and 546 (EX: 488/14 nm; EM: 590/30 nm), Gain = 1733; G/R = FRET signal of Alexa Fluor® 488 and 647 (EX: 488/14 nm; EM: 680/30 nm), Gain = 2554 and B/R = FRET signal of Alexa Fluor® 546 and 647 (EX: 540/20 nm; EM: 680/30 nm), Gain = 2607. Fitted data is collected in the data repository accompanying this publication

| Species | Blue | Green | Red | B/G | G/R | B/R |
|---|---|---|---|---|---|---|
| $M_1N_1$ | 372 | 0.96 | 820 | 77.7 | 205 | **1300** |
| $M_1N_2$ | 128 | 929 | 2.97 | **1315** | 144 | 255 |
| $M_1N_3$ | **1338** | 1.65 | 2.85 | 280 | 1.14 | 37.4 |
| $M_2N_1$ | 4.75 | 360 | 847 | 73.4 | **1281** | 244 |
| $M_2N_2$ | 13 | **1353** | 0.15 | 277 | 213 | 58.3 |
| $M_2N_3$ **or** $N_2$ | 12.8 | **1197** | 0.09 | 243 | 182 | 50 |
| $M_3N_1$ | 1.09 | 0.43 | **1358** | 0 | 323 | 47.9 |
| $M_3N_2$ | 2.16 | 153 | 781 | 32.6 | **1395** | 264 |
| $M_3N_3$ **or** $N_1$ | 0.61 | 0.43 | **900** | 0 | 211 | 29.8 |
| $M_1L_1$ | **39.9** | 0.1 | 0.13 | 9.57 | 0.14 | 1.68 |
| $M_2L_2$ | 1.65 | **140** | 0.16 | 22.4 | 19.6 | 5.8 |
| $M_3L_3$ | 0.88 | 0.11 | **97** | 0.13 | 22.9 | 3.84 |

### 3 Supplementary Note 3: Experimental procedures

Variations in both the concentration of stock oligonucleotides and pipetting and injection procedures mean that the actual concentration of strands in solution will differ from that intended in the experimental design. In some cases, we attempted to infer the actual concentration of strands in the experiment, as detailed below. Where a distinction is necessary, we refer to the "intended" concentration of the experimental design, and the "inferred" concentration apparent in experiments.

The designed experiments consisted of the measurement of reaction kinetics, alongside a set of ancillary measurements to determine reactant concentrations.[4] All experiments contained at least one negative (experimental buffer only) and one positive control per fluorescence channel measured. these controls were subjected to the same volume changes as the measured kinetics. Positive controls contained a variable concentration of fluorophore-labeled strands in the experimental buffer. Controls allowed the correction of changes in fluorescence due to environmental conditions. Data correction using the controls' signal is further described in Supplementary Note 4.

All ancillary measurements contained at least 10 data points and quantified the fluorescence baseline of a steady state in the experiment. These measurements were used to estimate the concentration of each species in the system from the fluorescence signal steady states and were obtained either before the start of the reaction or by sequentially triggering the reaction of all the species (Supplementary Figure 5). These ancillary measurements helped to diminish the uncertainty in concentrations that can arise from a range of factors, including pipetting errors. The number and nature of the ancillary measurements were experiment-contingent; however, all the different experiments followed a similar structure, outlined below. In each case, the median value obtained during an ancillary measurement was used as "the" value of the measurement.

Measurement 1. Initial fluorescence: All experiments started by mixing a quenched complex (usually $M_xL$) in experimental buffer, for a total volume of 140 µL. The measurement was made after 30 minutes of incubation at the experiment's temperature when the fluorescence value of the positive controls became stable.

Measurement 2. Additional species: For catalytic dimerization experiments that required the addition of template, 10 µL of a solution containing $T_{xy}$ was added to the well. The addition of a template made an equimolar amount of complex $M_xL$ react, increasing the fluorescence baseline.

Measurement 3. Experimental kinetics: This step involves the addition of 50 µL of a solution containing a reaction trigger species, usually $N_y$, resulting in a final reaction volume of 200 µL. In most cases, rather than perform replica experiments at identical concentrations, we used variants in which a single input species had a different concentration. These "replicas" allow a qualitative test for consistency and provide insight into the nature of the underlying reaction kinetics. We use the notation $[X, Y]$ to denote a set of concentrations in the range $X$ to $Y$ (inclusive).

The system kinetics were measured after the injection until the reaction reached a steady state. Kinetics that took more than 2 hours to reach a steady state were measured continuously for at least 50 hours after completely sealing the plate to prevent evaporation. After these measurements, the plate was incubated for up to a month to try to reach a steady state, sometimes after the addition of a low concentration of a reaction catalytic species, such as $T_{xy}$, to accelerate the approach to a steady state. The incubation was done at the experimental temperature and in dark conditions to avoid fluorophore photobleaching.

Measurement 4. Estimation of reaction trigger concentration: After the reaction reached a steady state, the resultant fluorescence plateau was recorded to infer the total amount of reaction trigger added in reactions (under the assumption that the reaction is irreversible).

Measurement 5. Saturation and quantification of the remaining species: To finish the assay, experiments were saturated with a combination of $N_y$ and $T_{xy}$. This way, all the $M_x$ monomer will be converted into $M_xN_y$ complex, facilitating its quantification. Note that the fluorescence of the different reaction intermediaries can have variations up to 30% when compared to $M_xN_y$ (Supplementary Note 2.1).

The specific procedures for each type of experiment are discussed and collected in tables in the following subsections. As explained in Supplementary Note 2, species concentration is reported at any time as the concentration if the same amount of solute was dissolved in a reaction volume of 200 µL. This volume was chosen for being the one at which kinetics were always measured.

#### 3.1 Single fluorescence channel experiments

##### 3.1.1 Simple 2nd order reactions ($M_x$ binding and monomer leak reactions)

The protocols, given in Supplementary Tables 7 and 8, do not include 'Measurement 2' as no initial $T_{xy}$ is present in these experiments.

Binding rate of $M_xL$ to $T_{xy}$ (displacement of $L$ from $M_xL$ by $T_{xy}$). We measured the kinetics of displacement of $L$ from $M_xL$ by $T_{xy}$ for every $T_{xy}$ strand used during this work and fitted a TMSD rate constant $k_t$ to the results. Experiments consisted of a solution of $M_xL$ ($[M_xL]_0 = 100$ nM) triggered by a variable concentration of a complementary $T_{xy}$ ($[T_{xy}]_0$ between 20 and 112.5 nM). The aim of testing some experimental replicas with $[T_{xy}]_0 > [M_x]_0$ was to confirm that $T_{xy}$ reacts with a 1:1 stoichiometry with $M_xL$ and discard a potential equilibration with its reversible reaction. Protocol steps are contained in Supplementary Table 7 and illustrated in Supplementary Figure 5. Additionally, Supplementary Figure 5 demonstrates that the fluorescence signal is proportional to the amount of $T_{xy}$ injected.

$M_xL$ and $N_3$ leak reactions. The kinetics of template-free leak reaction was obtained for the reaction between $M_x$ and $N_3$ using single-channel fluorescence experiments. These data were used to fit a leak reaction rate constant, $k_{\text{leak}}$. Experiments consisted of a solution of $M_xL$ ($[M_xL]_0 = 100$ nM) triggered by $N_3$ ($[N_3]_0 = [40, 90]$ nM). During 'Measurement 4', 5 nM of $T_{x3}$ was injected for the reaction to reach completion. The estimated signal produced from the extra $T_{x3}$ was later removed to estimate $[N_3]_0$. Protocol steps are contained in Supplementary Table 8.

**Supplementary Table 7: Detailed experimental procedure for $M_x$-binding characterisation.** Several trajectories for each $T_{xy}$ condition and their controls were triggered by mechanical injection and measured simultaneously. The triggering species was a solution of $T_{xy}$ with a complementary primary toehold. Since no $T_{xy}$ was initially present in the experiment, no 'Measurement 2' was needed.

| Steps | Addition | Purpose | Expected concentrations in 200 µL (nm) | Total volume (µL) |
|---|---|---|---|---|
| **Measurement 1** | 10 µL $M_xL$ (2 µM) + 140 µL Buffer | Estimate residual fluorescence baseline | $M_xL = 100$ | 150 |
| **Measurement 3** | [50,10] µL $T_{xy}$ (450 nM) + [0,40] µL Buffer | Experimental kinetics | Evolve with time | 200 |
| **Measurement 4** | - | Estimate $[T_{xy}]_0$ | $M_xL$ = [77.5,0]; $M_1T_{xy}$ = [22.5,100]; Lock:[22.5,100] | 200 |
| **Measurement 5** | 5 µL each of $N_3$ and $T_{x3}$ (10 µM) | Estimate $[M_xL]_0$ | $MN_{x3} = 100$; $L = 100$; $T_{xy}$ and $N_3 \approx 200$ | 210 |

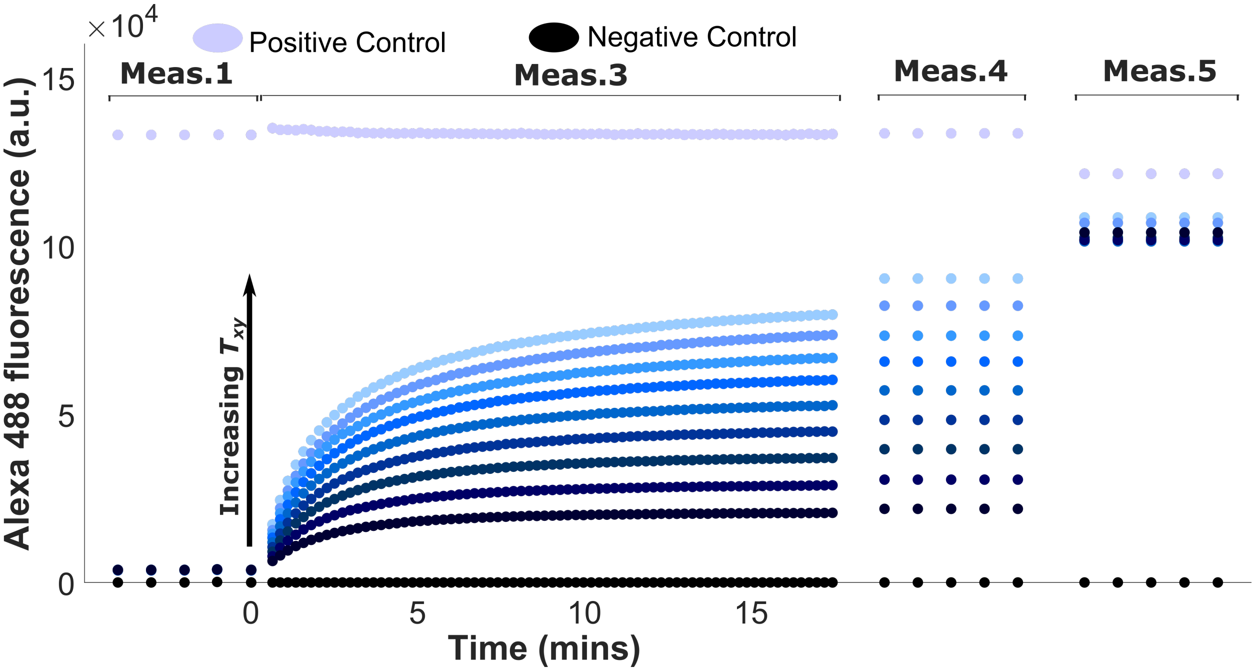


**Supplementary Figure 5: Example of $M_xL$ binding characterisation.** Raw data from the displacement of $L$ from the $M_1L$ complex by adding strand $T_{12}$ at 25° C. 'Measurement 1' contains the baseline of the experiments, considered as 0 nM of reacted $M_1L$. During 'Measurement 3', increasing amounts of $T_{12}$ demonstrate that the signal produced is proportional to the concentration of $T_{12}$. After the reaction reached a steady state, $[T_{12}]$ was estimated from 'Measurement 4'. Afterwards, the solution was saturated with $N_3$ and $T_{13}$ to transform all $M_1L$ into $M_1N_3$ (the large excess of $N_3$ was sufficient to displace $T_{12}$ from $M_1T_{12}$ by the secondary toehold within a day). Fluorescence signal increased even for saturated conditions due to the transformation of $M_1T_{12}$ into the complex $M_1N_3$. Concentrations: $[M_1L]_0 = 100$ nM, $[T_{12}]_0$ = [101.25, 22.5] nM in increments of 11.25 nM. The positive control contained around 100 nM of $M_1N_3$. The negative controls consisted of only the experimental buffer.

**Supplementary Table 8: Detailed experimental procedure for leak reaction characterisation for single-fluorescence channel monomers.** Several trajectories were triggered by mechanical injection and measured simultaneously. The triggering species was a solution of $N_3$. Since no $T_{xy}$ was initially present in the experiment, no 'Measurement 2' was needed.

| **Steps** | **Addition** | **Purpose** | **Expected concentrations in 200** µL (nM) | **Total volume** (µL) |
|---|---|---|---|---|
| **Measurement 1** | 10 µL $M_xL$ (2 µM) + 140 µL Buffer | Estimate residual fluorescence baseline | $M_xL = 100$ | 150 |
| **Measurement 3** | [45,10] µL $N_3$ (400 nM) + [5,40] µL Buffer | Experimental kinetics | Evolve with time | 200 |
| **Measurement 4** | 5 µL $T_{x3}$ (200 nM) | Estimate $[N_3 + T_{x3}]$ | $M_xL$ = [75, 5]; $M_xN_3$ = [20, 90]; $L$ = [25, 95]; $T_{x3}M_x = 5$ | 205 |
| **Measurement 5** | 5 µL each of $N_3$ and $T_{x3}$ (10 µM) | Estimate $[M_xL]_0$ | $M_xN_3 = 100$; $L = 100$; $T_{x3}$ & $N_3 \approx 200$ | 215 |

##### 3.1.2 Handhold-mediated reaction of $\boldsymbol{M_1T_{13}}$ with $\boldsymbol{N_3}$

For a subset of the $T_{13}$ variants, we studied the reaction between $M_1T_{13}$ and $N_3$. The kinetics of this reaction could be measured by harnessing the significant increase in fluorescence resulting from the formation of $M_1N_3$ from $M_1T_{13}$. Experiments consisted of a solution of preannealed $M_1T_{13}$b($u$t/$v$h) ($[M_1T_{13}]_0$ = 100 nM) triggered by $N_3$ ($[N_3]_0$ = [40, 90] nM). Protocol steps for handhold experiments are identical to the ones followed in Supplementary Table 8 except for no addition of $T_{xy}$ in 'Measurement 4', no 'Measurement 5', and exchanging $M_xL$ for $M_1T_{13}$.

##### 3.1.3 Dimerization experiments using $\boldsymbol{M_x}$, $\boldsymbol{T_{x3}}$ and $\boldsymbol{N_3}$

These experiments were performed to explore system kinetics and product inhibition as a function of the length of primary toehold and handhold domains. We considered two regimes of monomer concentration for these experiments:

1. High monomer concentration experiments consisted of a solution of $M_xL$ ($[M_xL]_0 = 100$ nM) triggered by $N_3$ ($[N_3]_0$ = 100 nM), 24 hours after adding a variable amount of $T_{x3}$ ($[T_{x3}]_0$ = [10 - 0.25] nM) into the $M_xL$ solution. These conditions were used to obtain the total turnover assays for all nine $M_xN_y$ products (Figure 3c and Supplementary Figures 16 to 18).

2. Low monomer concentration experiments consisted of a solution of $M_1L$ ($[M_1L]_0 = 10$ nM) triggered by $N_3$ ($[N_3]_0$ = 10 nM), 24 hours after adding 1 nM of $T_{13}$ into the $M_1L$ solution. These conditions were used during the turnover rate and inhibition assays for the formation of $M_1N_3$ catalyzed by $T_{13}$ (Figures 2, 3a, and Supplementary Figure 15).

Protocol steps are summarised in Supplementary Table 9 and illustrated in Supplementary Figure 6.

**Supplementary Table 9: Detailed experimental procedure to measure dimerization kinetics.** Several trajectories with different $T_{x3}$ concentrations and their controls were triggered by mechanical injection and measured simultaneously. The trigger species was $N_3$. 'Measurement 2' and 'Measurement 4' aimed to estimate the amount of $T_{x3}$ and $N_3$ in the solution, respectively. However, the estimated concentrations were highly imprecise due to the reactions never completely reaching steady-state.

| Steps | Addition | Purpose | Expected concentrations in 200 µL (nm) | Total volume (µL) |
|---|---|---|---|---|
| **Measurement 1** | 10 µL $M_xL$ (2 µM) + 130 µL Buffer | Estimate residual fluorescence baseline | $M_xL = 100$ | 140 |
| **Measurement 2** | [10,1] µL $T_{x3}$ (100 nM) + [0, 9] µL Buffer | Estimate $[T_{x3}]_0$ | $M_xL = [95, 99.75]$; $L = [5, 0.25]$; $M_xT_{x3} = [5, 0.25]$ | 150 |
| **Measurement 3** | 50 µL $N_3$ (400 nM) | Experimental kinetics | Evolve with time | 200 |
| **Measurement 4** | 5 µL $T_{x3}$ (200 nM) | Estimate $[N_3 + T_{x3}]$ | Equilibrium between $M_XN_Y$ $M_xT_{x3}$ and $T_{x3}$ | 205 |
| **Measurement 5** | 5 µL each of $N_3$ and $T_{x3}$ (10 µM) | Estimate $[M_xL]_0$ | $M_xN_3 = 100$; $L = 100$; $T_{x3}$ and $N_3 \approx 200$ | 215 |

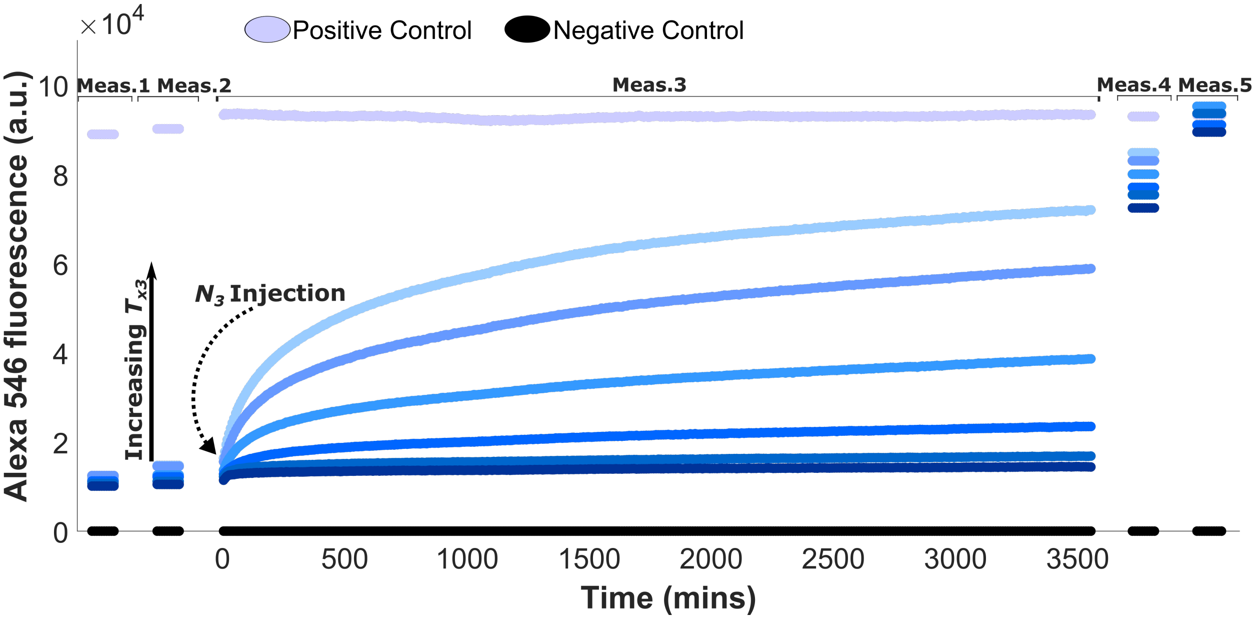


**Supplementary Figure 6: Example of dimerization experiments with $N_3$ (single fluorescent channel).** Raw data showing the displacement of $L$ from a $M_2L$ complex by addition of the strands $N_3$ and $T_{23}$ at 25° C. 'Measurement 1' contains the baseline of the experiments, which are considered as 0 nM of reacted $M_2$. During 'Measurement 2', variable concentrations of $T_{23}$ produced an increment of the baseline. After that, the complete catalytic cycle was closed by the addition of $N_3$. After measuring the kinetics for enough time, a small quantity of $T_{23}$ (5 nM) was added to all wells to promote the consumption of all $N_3$. After one day of further incubation, the new baseline wass measured in 'Measurement 4'. Afterwards the solution was saturated with $N_3$ and $T_{23}$ to transform all $M_2L$ and quantify the total amount of $M_2L$ in 'Measurement 5'. Concentrations: $[M_2L]_0 = 100$ nM, $[T_{23}]_0 = [0.25, 0.5, 1.25, 2.5, 3.75$ and $5$ nM$]$, $[N_3]_0 = 100$ nM. The positive control contained around 100 nM of $M_2N_3$. The negative controls consisted of only the experimental buffer.

Product inhibition testing. The step-by-step protocol for experiments testing product inhibition are given in Supplementary Table 10. When studying product inhibition, the experiment included a variable initial concentration of annealed $M_1$(Unlab.)$N_3$. This unlabeled product was added before triggering the reaction kinetics with $N_3$ during 'Measurement 3'. The function of the non-fluorescent $M_1N_3$ dimer was to inhibit the consumption of the fluorescent $M_1L$ complex via product inhibition, without producing a high fluorescent background that would have hindered the interpretation of the kinetics. The non-fluorescent $M_1N_3$ was annealed with a 10% excess of $N_3$ over $M_1$ (Unlab.) to avoid sequestration of the $N_3$ strands intended to drive the reaction. Due to the high concentration of inhibitory species in the experiment, $N_3$ quantification during 'Measurement 4' was unreliable. $[N_3]_0$ was instead assumed to be the intended concentration plus the 10% excess resultant from $M_1$(Unlab.)$N_3$ annealing.

Inhibition experiments tested the kinetics with a low monomer concentration ($[M_1L]_0 =$ 10 nM; $[N_3]_0 = 10$ nM; $[T_{13}\ (ut/vh)]_0 = 1$ nM). For each ($ut/v$h) for the template $T_{1,3}$, eight different unlabeled $[M_1N_3]_0$ conditions were tested: 0, 2.5, 5, 10, 15, 25, 50 and 100 nM. A subset of $ut/v$h was also tested at higher concentrations ($[M_1L]_0 = 100$ nM; $[N_3]_0 = 100$ nM; $[T_{13}\ (ut/vh)]_0 =$ 2,5, 5 and 10 nM; $[M_1N_3]_0 =$ 0, 50, 100, 200 and 300 nM).

**Supplementary Table 10: Detailed experimental procedure to test product inhibition.** Several trajectories with different unlabeled $[M_1N_3]_0$ concentrations and their controls were triggered by mechanical injection and measured simultaneously. The trigger species was $N_3$. 'Measurement 2' aimed to estimate the amount of $T_{x3}$ in the solution respectively. Due to the excess of product in the solution, no 'Measurement 4' could be included. This table describes experiments at a 10 nM-scale.

| Steps | Addition | Purpose | Expected concentrations in 200 µL (nm) | Total volume (µL) |
|---|---|---|---|---|
| **Measurement 1** | 10 µL $M_xL$ (200 nM ) + 130 µL Buffer | Estimate residual fluorescence baseline | $M_xL = 10$ | 140 |
| **Measurement 2** | 10 µL $T_{x3}$ (20 nM) | Estimate $[T_{x3}]_0$ | $M_xL = 9$; $L = 1$; $M_xT_{x3} = 1$ | 150 |
| **Measurement 3** | 40 µL $N_3$ (50 nM) 10 µL $M_1N_3$ ([0, 2] µM) | Experimental kinetics | Evolve with time | 200 |
| **Measurement 5** | 5 µL each of $N_3$ and $T_{x3}$ (10 µM) | Estimate $[M_xL]_0$ | $M_xN_3 = 10$; $L = 10$; $T_{x3}$ and $N_3 \approx 200$ | 210 |

#### 3.2 Multichannel fluorescence experiments

This section describes experiments that require measuring the direct fluorescence of, and FRET between, several fluorophores. The experiments in question are those in which both $M_x$ and $N_y$ are labeled.

##### 3.2.1 Leak and dimerization experiments with labeled $N_1$ or $N_2$

The experiments explored the kinetics of systems consuming monomers $N_1$ and $N_2$. The objective, sequence of measurements, and other DNA strands involved in these experiments were similar to their single-fluorophore version involving $N_3$; the main difference is that fluorescence measurements were performed in multiple channels. Measurements 1 to 3 included measurements in up to three fluorescent channels simultaneously, depending on which fluorophores were present: a channel for $M_xL$, a channel for $N_y$, and a channel to follow the FRET signal of the resultant $M_xN_y$ product.

'Measurements 4 and 5' were removed from the experimental protocol. The inferred $[N_y]_0$ could be quantified from its signal just after injection in 'Measurement 3'. 'Measurement 5' was removed due to the difficulty of estimating $M_xL$ through saturation of the system with labeled $N_y$. If labeled $N_y$ were used to saturate the experiment, the significant increase of signal would interfere with the quantification of the product FRET signal. On the other hand, adding $N_3$ would produce, besides the assay product, an undetermined amount of product $M_xN_3$, impeding the quantification of $M_xL$. Instead, since all experiments were performed at high $M_xL$ concentration, the quenched signal of the complex in 'Measurement 1' was used to infer its initial concentration.

Protocol steps are summarised in Supplementary Table 11 and a multifluorophore dimerization experiment is illustrated in Supplementary Figure 7.

**Supplementary Table 11: Detailed experimental procedure for dimerization kinetics with labeled $N_y$.** Several trajectories with different $T_{xy}$ concentrations and their controls were triggered by mechanical injection and measured simultaneously. The trigger species was a labeled $N_y$ strand. During these experiments, 'Measurement 1' was used to estimate the amount of $M_xL$, while the concentration of $N_y$ was extracted from the initial values of 'Measurement 3'.

| Steps | Addition | Purpose | Expected concentrations in 200 µL (nm) | Total volume (µL) |
|---|---|---|---|---|
| **Measurement 1** | 10 µL $M_xL$ (2 µM ) + 130 µL Buffer | Estimate $[M_xL]_0$ | $M_xL = 100$ | 140 |
| **Measurement 2** | [10,0.5] µL $T_{xy}$ (100 nM) | Estimate $[T_{xy}]_0$ This step is not used for leak reactions | $M_xL$ = [95, 99.75]; $L$ = [5,0.25]; $M_xT_{xy}$ = [5,0.25] | 150 |
| **Measurement 3** | 50 µL $N_y$ ([400] nM) | Experimental kinetics and estimate $[N_y]_0$ | Evolve with time | 200 |

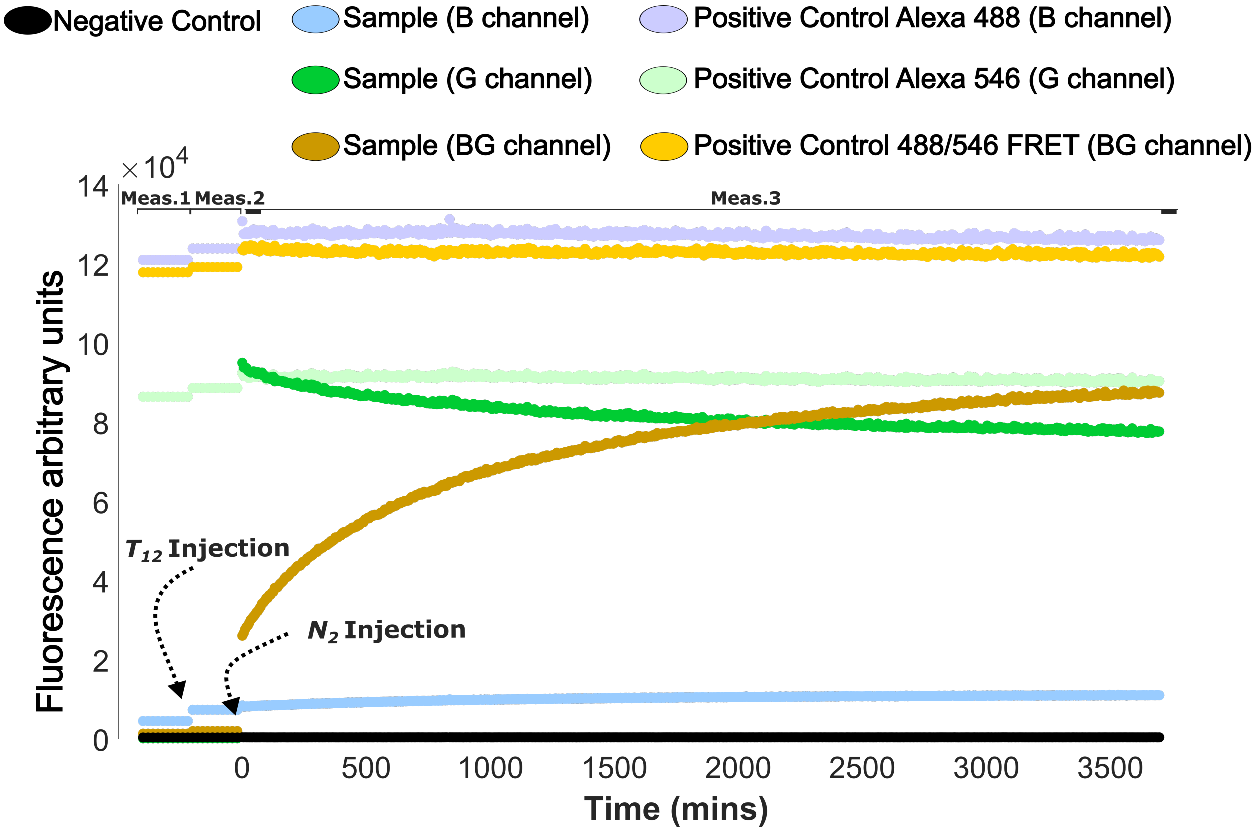


**Supplementary Figure 7: Example of dimerization experiments with labeled $N_y$ (multichannel fluorescence experiment).** Raw data from the reaction resulting from triggering the release of $L$ from $M_1L_1$ complexes by adding $N_2$ and $T_{12}$ at 25° C. 'Measurement 1' contains the baseline of the experiments, considered as 0 nM of reacted $M_1$. This baseline used to estimate $[M_1L]_0$. During 'Measurement 2', adding $T_{12}$ produces an increment of the baseline that can be quantified. After that, the complete catalytic cycle is closed by adding $N_2$. The resultant product $M_1N_2$ produces a FRET signal used to monitor the kinetics. $[N_2]_0$ is estimated from the green signal at the beginning of this kinetic trace. Concentrations: $[M_1L]_0$ = 100 nM, $[T_{12}]_0 = 5$ nM], $[N_2]_0 = 100$ nM. The positive controls contained around 100 nM of $M_1N_3$, $N_2$, or an equimolar mix of both for FRET positive control. The negative controls consisted of only the experimental buffer.

##### 3.2.2 Sequence-specific copying by templated dimerization

The objective of these experiments was both to qualitatively confirm the specificity of templated catalytic assembly and to provide a quantitative estimate of yields. The experiments started with 140 µL containing the three $M_xL$ complexes ($M_{1-3}L$) at a concentration of 100 nM to which 10 µL of solution containing $T_{xy}$ was added, for a final concentration of either 5 nM or 10 nM. The reaction was triggered by adding 50 µL of a solution containing the three species of $N_y$ ($N_{1-3}$) for a final concentration of 100 nM or 75 nM. Three sets of experiments with slightly different initial concentrations were used but in all cases $[M_xL]_0 \sim [N_y]_0 \gg [T_{xy}]_0$ (refer to Figure 5 and Supplementary Figures 22 and 24 for specific conditions). Each set of experiments contained simultaneously nine systems of interest (one per template), a leak control with a solution containing only the monomers at the same concentration (Supplementary Figure 26), and nine positive controls each containing approximately 75 nM of one of the possible $M_XN_y$ products. For each experiment, fluorescence was monitored in the 6 channels listed in Supplementary Table 6; we discuss the deconvolution of this data in Supplementary Note 4.2. The detailed protocol is described in Supplementary Table 12. An example raw data set of is shown in Supplementary Figure 8.

**Supplementary Table 12: Detailed experimental procedure for sequence-specific copying by templated dimerization.** For each set, ten wells, each containing all $M_xL$ and a single $T_{xy}$ (or none, in negative control) were triggered by mechanical injection of a solution with all $N_y$ and measured simultaneously in six fluorescence channels. During these experiments, 'Measurement 1' was used to estimate the amount of $M_xL$, while the concentration of $N_y$ was extracted from the initial values of 'Measurement 3'.

| **Steps** | **Addition** | **Purpose** | **Expected concentrations in 200** µL **(nm)** | **Total volume** (µL) |
|---|---|---|---|---|
| **Measurement 1** | 10 µL $M_{1-3}L$ (2 µM ) + 110 µL Buffer | Estimate $[M_xL]_0$ | $M_{1-3}L = 100$ | 140 |
| **Measurement 2** | [10] µL $T_{xy}$ ([100-200] nM) | Estimate $[T_{xy}]_0$ | $M_xL$ = [90, 95] (non-complementary $M_xL$ = 100); $L$ = [10, 5]; $M_xT_{xy}$ = [5, 0.5] | 150 |
| **Measurement 3** | 50 µL $N_{1-3}$ ([400-300] nM) | Experimental kinetics and estimate $[N_{1-3}]_0$ | Evolve with time | 200 |

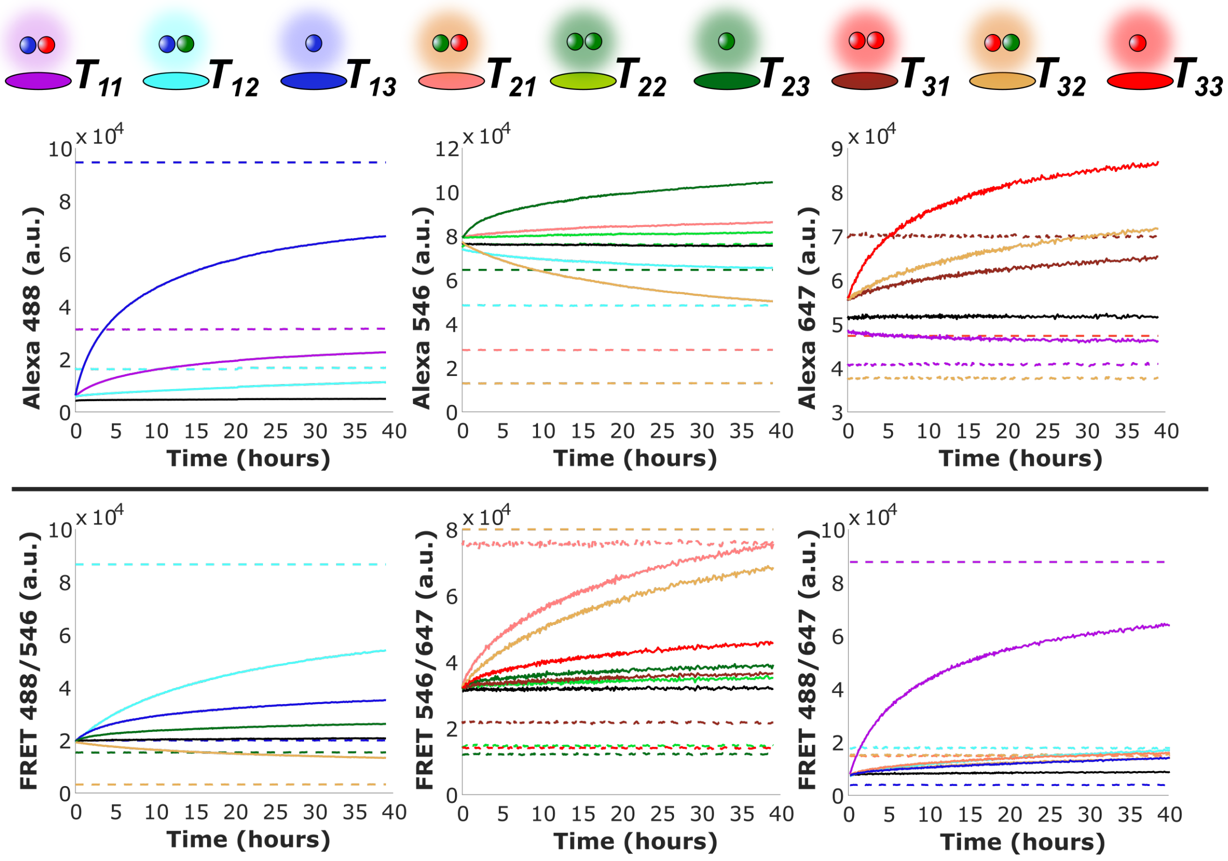


**Supplementary Figure 8: Raw data showing the kinetics of the sequence-specific copying by templated dimerization.** Fluorescence signal produced in the six measured fluorescence channels for the nine reactions catalyzed by different templates $T_{xy}$. Only data showing a significant deviation from the leak control is plotted, for clarity. This data is deconvoluted (Supplementary Note 4.2) using the fluorescence calibration matrix in Supplementary Table 5 to obtain estimates of the concentrations of products. This raw data corresponds to the 3rd iteration, with processed data shown in Figure 5 of the main text. Legend: The kinetics of each $T_{xy}$ experiment at the different fluorescence channels are represented by the colors shown in the ovals. The 'halo' of each fluorophore dimer above the ovals indicates the channel at which higher fluorescence contribution is expected for their intended product $M_xN_y$ (Blue, Alexa 488; green, Alexa 546; red, Alexa 647; cyan, FRET 488/546; orange, FRET 546/648; purple, FRET 488/648). Dashed lines: Fluorescence produced by the $M_xN_y$ control corresponding to each template. This control may appear below the kinetics signal due to the background resulting from unconsumed $N_y$ in the kinetic measurements. Black lines: Fluorescence value of the template-free leak control. Trajectories not represented followed the same trend as the leak control. Intended concentrations: Each $[M_xL]_0 = 100$ nM, each $[N_y]_0 = 75$ nM, $[T_{xy}]_0 = 5$ nM.

### 4 Supplementary Note 4: Data processing

We now discuss how raw fluorescent signals are converted into inferred molecular concentration. Fluorescence signals were processed following a method similar to Cabello *et al.*[4] Subsequent subsections describe how the signal is transformed in each particular set of experiments.

#### 4.1 Single fluorescence channel experiments

##### 4.1.1 Fluorescence coefficient variability

During the recording of the single fluorophore experiments, the bulb of the plate reader had irregular behavior and had to be replaced. Therefore, the absolute calibration reported in Supplementary Note 2 couldn't be used directly to infer concentrations in all cases. Instead, where necessary, we assumed that the initial concentration of $M_x$ (present either in $M_xL$ of $M_xT_{xy}$ complexes) was exactly equal to the intended concentration when averaged over all experiments performed in parallel. We then used the ancillary measurement of total $M_x$ concentration to estimate a coefficient of fluorescence for that complex in the experiment in question. In general, this coefficient differed slightly from the one estimated in Supplementary Note 2. Our second assumption was that all fluorescent species had their coefficient of fluorescence scaled by the same factor relative to the values reported in Supplementary Note 2. In practice, the implied deviation in fluorescence coefficient was typically 10% or less, and the later results (performed with the new bulb) had coefficients consistent with the calibration in Supplementary Note 2 (performed with the new bulb).

##### 4.1.2 Inferring molecular concentrations during displacement of $L$ from $M_xL$ by $T_{x3}$

Let $F_i$ be the fluorescence at timepoint $i$ during 'measurement 3' for a given reaction, having subtracted the fluorescence of the negative control (buffer only). We first rescale this fluorescence to remove environmental fluctuations evident in the positive control. We do so by rescaling by $F_1^+/F_i^+$, where $F_i^+$ is the positive control signal at time $i$ and $F_1^+$ is the positive control recorded during measurement 1. This $F_i' = (F_1^+/F_i^+)F_i$ is then rescaled again to give a fractional conversion into $M_xT_{x3}$

$$\frac{[M_xT_{x3}]_i}{[M_xL]_0} = \frac{F_i' - F_1}{\left(F'_{\mathrm{T,max}} - F_1\right)}. \tag{1}$$

Here $F_1$ is the residual fluorescence during 'Measurement 1' due to the quenched monomers, and $F'_{\mathrm{T,max}}$ is a putative maximal fluorescence if all of the $M_x$ strands were bound to the template. In practice, such a state is not obtained in the experiment; instead, in 'Measurement 5' all monomers are converted to $M_xN_3$. Let $F'_{\mathrm{N,max}}$ be the (positive control rescaled) fluorescence recorded during 'Measurement 5'. Then

$$[M_xT_{x3}]_i = [M_xL]_0 \frac{F_i' - F_1}{\left(F'_{\mathrm{N,max}} \frac{a_{M_xT_{x3}}}{a_{M_xN_3}} - F_1\right)} = \frac{a_{M_xN_3}}{a_{M_xT_{x3}}} [M_xL]_0 \frac{F_i' - F_1}{\left(F'_{\mathrm{N,max}} - F_1 \frac{a_{M_xN_3}}{a_{M_xT_{x3}}}\right)}, \tag{2}$$

where $\frac{a_{M_xN_3}}{a_{M_xT_{x3}}}$ is the ratio of fluorescence coefficients for the two complexes. As outlined above, we assume $\frac{a_{M_1N_3}}{a_{M_1T_{13}}}$ has the value given in Supplementary Section 2.1 (1.37). $\frac{a_{M_xN_3}}{a_{M_xT_{x3}}}$ for $x = 2, 3$ are assumed to be 1, as in those cases the fluorophores showed no variation due to binding with unlabeled strands.

It is assumed that the mean of $F'_{\mathrm{N,max}}$ for all reactions involving the same monomer $M_x$ corresponds to the fluorescence of the intended 100 nM of $M_xN_3$. The inferred value of $[M_xL]_0$ for each reaction was then calculated by multiplying 100 nM by the value of $F'_{\mathrm{N,max}}$ recorded in that reaction, divided by this mean fluorescence. The total concentration of template, $[T_{x3}]_0$, was inferred from the value recorded during measurement 4, $F_4'$, by replacing $F_i'$ with $F_4'$ in Supplementary Equation 2, and assuming $[T_{x3}]_0 = [M_xT_{x3}]_4$.

###### 4.1.3 Inferring molecular concentrations during the leak reaction between $M_xL$ and $N_3$

Let $F_i$ be the fluorescence at timepoint $i$ during 'measurement 3' for a given reaction, having subtracted the fluorescence of the negative control (buffer only). We first rescale this fluorescence to remove environmental fluctuations evident in the positive control. We do so by rescaling by $F_1^+/F_i^+$, where $F_i^+$ is the positive control signal at time $i$ and $F_1^+$ is the positive control recorded during measurement 1. This $F_i' = (F_1^+/F_i^+)F_i$ is then rescaled again to give a fractional conversion into $M_xN_3$

$$\frac{[M_xN_3]_i}{[M_xL]_0} = \frac{F_i' - F_1}{(F'_{\text{max}} - F_1)}. \tag{3}$$

Here $F_1$ is the residual fluorescence during 'Measurement 1' due to the quenched monomers, and $F'_{\text{max}}$ is the maximal fluorescence observed during 'Measurement 5', when all monomers $M_xL$ are converted into $M_xN_3$.

It is assumed that the mean of $F'_{\text{max}}$ for all reactions involving the same variant of monomer $M_x$ corresponds to the fluorescence of the intended 100 nM of $M_xN_3$. The inferred value of $[M_xL]_0$ for each reaction was then calculated by multiplying 100 nM by the value of $F'_{\text{max}}$ in that reaction, divided by this mean fluorescence. The inferred value of $[N_3]_0$ was extracted from 'Measurement 4'.

###### 4.1.4 Inferring molecular concentrations during HMSD between $M_1T_{13}$ and $N_3$

Let $F_i$ be the fluorescence at timepoint $i$ during 'Measurement 3' for a given reaction, having subtracted the fluorescence of the negative control (buffer only). In a slight abuse of notation, we will use $F_1$, $F_2$ etc. to refer to specific values of fluorescence obtained in 'Measurement 1', 'Measurement 2' etc., and use $F_i$ for arbitrary time points during 'Measurement 3'.

We first rescale $F_i$ to remove environmental fluctuations evident in the positive control. We do so by rescaling by $F_1^+/F_i^+$, where $F_i^+$ is the positive control signal at time $i$ and $F_1^+$ is the value of the positive control recorded during 'Measurement 1'. This $F_i' = (F_1^+/F_i^+)F_i$ is then rescaled again to give a fractional conversion into $M_xN_3$

$$\frac{[M_1N_3]_i}{[M_1T_{13}]_0} = \frac{F_i' - F_1}{\left(F'_{\text{N,max}} - F_1\right)}. \tag{4}$$

Here $F_1$ is the residual fluorescence during 'Measurement 1' due to the template-bound monomers, and $F'_{\text{N,max}}$ is the putative maximal fluorescence if all of the $M_x$ strands were bound to $N_3$. In practice, we infer $F'_{\mathrm{N,max}}$ from $F_1$, using $F'_{\mathrm{N,max}} = \frac{a_{M_1N_3}}{a_{M_1T_{13}}} F_1$ and the value of $\frac{a_{M_1N_3}}{a_{M_1T_{13}}} = 1.37$ from Supplementary Section 2.1. Thus

$$[M_1N_3]_i = [M_1T_{13}]_0 \frac{F'_i - F_1}{\left(F_1 \frac{a_{M_1N_3}}{a_{M_1T_{13}}} - F_1\right)}. \tag{5}$$

It is assumed that the mean of $F_1$ for all reactions involving the same variant of $M_1$ corresponds to the fluorescence of the intended 100 nM of $M_1T_{13}$. The inferred value of $[M_1T_{13}]_0$ for each reaction was then calculated by multiplying 100 nM by the value of $F_1$ for that reaction, divided by this mean fluorescence. The total concentration of $N_3$, $[N_3]_0$, was calculated assuming $[N_3]_0 = [M_1N_3]_4$, the inferred concentration during measurement 4. $[M_1N_3]_4$ was inferred from Supplementary Equation 5 with the fluorescence recorded in measurement 4, $F'_4$, replacing $F'_i$.

###### 4.1.5 Inferring molecular concentrations for templated dimerization of $M_xL$ and $N_3$

The processing of the fluorescence to obtain inferred concentration time series is illustrated in Supplementary Figure 9. Let $F_i$ be the fluorescence at timepoint $i$ during 'measurement 3' for a given reaction, having subtracted the fluorescence of the negative control (buffer only). We first rescale this fluorescence to remove environmental fluctuations evident in the positive control. We do so by rescaling by $F_1^+/F_i^+$, where $F_i^+$ is the positive control signal at time $i$ and $F_1^+$ is the positive control recorded during measurement 1. This $F'_i = (F_1^+/F_i^+)F_i$ is assumed to be the sum of the fluorescence from three species: the product $M_xN_3$ with fluorescence $F_i^*$, the template-bound monomer $M_xT_{x3}$ and the initial monomer $M_xL$.

For simplicity, we assume that during the reaction kinetics (measurement 3) all template strands are bound to $M_x$. Thus the contribution of $M_xT_{x3}$ to the fluorescent signal should be constant, $F'_T$. Finally, the contribution of $M_xL$ will be equal to the initial fluorescence recorded in measurement 1, $F_1$, minus the contribution to that total of the monomers that have been converted into $M_xT_{xy}$ or $M_xN_3$.

$$F'_i = F^*_i + F'_T + \left(F_1 - \frac{a_{M_xL}}{a_{M_xN_3}}F^*_i - \frac{a_{M_xL}}{a_{M_xT_{x3}}}F'_T\right). \tag{6}$$

In this expression, $\frac{a_{M_xL}}{a_{M_xN_3}}F^*_i$ estimates the fluorescence contribution of the $M_xL$ complexes that have been converted into $M_xN_3$, and $\frac{a_{M_xL}}{a_{M_xT_{x3}}}F'_T$ the fluorescence of $M_xL$ complexes that have been converted into $M_xT_{xy}$. Rearranging,

$$[M_xN_3]_i = \frac{F^*_i}{a_{M_xN_3}} = \frac{F'_i - F'_T\left(1 - \frac{a_{M_xL}}{a_{M_xT_{x3}}}\right) - F_1}{a_{M_xN_3} - a_{M_xL}}. \tag{7}$$

This expression can be simplified using the ancillary measurements. Taking $F'_{\max}$ as the value recorded in the final measurement 5, when all $M_x$ is in $M_xN_3$ complexes and hence $[M_xN_3] = [M_xL]_0$,

$$[M_xN_3]_i = [M_xL]_0\frac{F'_i - F'_T\left(1 - \frac{a_{M_xL}}{a_{M_xT_{x3}}}\right) - F_1}{F'_{\max} - F_1}. \tag{8}$$

Continuing in this vein, the value recorded during measurement 2 ($F'_2$) is assumed to arise from a concentration of monomer-bound templates equal to the injected concentration, $[T_{xy}]_0$, plus the contribution of the remaining $M_xL$ complexes. Thus

$$F'_2 = F'_T + F_1\left(1 - \frac{a_{M_xL}}{a_{M_xT_{x3}}}F'_T\right), \tag{9}$$

or

$$F'_T\left(1 - \frac{a_{M_xL}}{a_{M_xT_{x3}}}\right) = F_1 - F'_2. \tag{10}$$

We therefore infer a product yield of

$$[M_xN_3]_i = [M_xL]_0\frac{F'_i - F'_2}{F'_{\max} - F_1}. \tag{11}$$

The inferred value of $[M_xL]_0$ for each reaction was calculated by multiplying 100 nM by the value of $F_1$ for that reaction, divided by this mean of all values of $F_1$ for the same input (effectively assuming that the mean concentration of $[M_xL]$ over all reactions involving that variant of $M_x$ is equal to the intended value). The injected template concentration, $[T_{xy}]_0$, was inferred as

$$[T_{xy}]_0 = \frac{F'_2 - F_1}{a_{M_xT_{x3}} - a_{M_xL}} = [M_xL]_0\frac{F'_2 - F_1}{F'_{\max}\frac{a_{M_xT_{x3}}}{a_{M_xN_3}} - F_1}, \tag{12}$$

Using the same assumptions as in the initial displacement of $L$ from $M_xL$ by $T_{xy}$, $\frac{a_{M_1T_{13}}}{a_{M_1N_3}} = 1/1.37$ and $\frac{a_{M_xT_3}}{a_{M_xN_3}} = 1$ for $x = 2, 3$. However, 'Measurement 2', from which $T_{x3}$ is estimated was highly imprecise, forcing the use of the intended $[T_{x3}]$, rather than a value obtained from 'Measurement 2', during subsequent analysis. Results from 'Measurement 2' were only considered as quality control to check the signal increase as increasing amounts of $[T_{x3}]_0$ were included in the experiment. The low precision during $[T_{x3}]_0$ quantification may have resulted from slow reaction rates and the large relative concentration of $M_xL$, potentially including some wrongly annealed complexes that mask the signal induced by $T_{x3}$ addition.

$[N_3]_0$ was estimated from transforming the signal in 'Measurement 5' using Supplementary Equation 11 and subtracting the signal expected from 5 nM of $T_{x3}$.

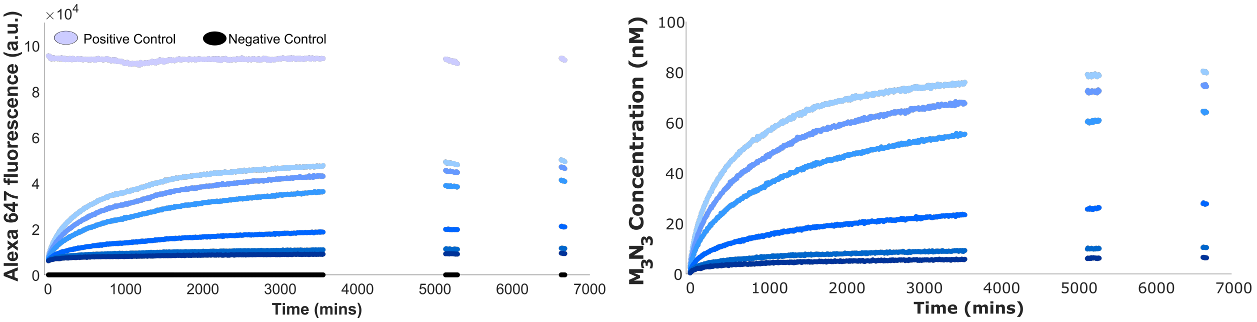


**Supplementary Figure 9: Example of the result of fluorescence data processing for single fluorophore experiments.** The kinetics contain six traces of the catalytic dimerization experiment of $M_3L$ and $N_3$ mediated by $T_{33}$. This case had a particularly non-uniform positive control signal. **Left)** Raw data. During long experiments, plate misalignment can produce bumps in the fluorescence that tampers the fitting process. These artifacts were corrected using the positive control, which has fluorescence intensity fluctuations proportional to those in the samples. **Right)** After the fluorescence processing pipeline. After correcting the fluorescence data it was transformed to nM. $[T_{33}]_0 = [0.25, 0.5, 1.25, 2.5, 3.75, 5]$ nM, $[N_3]_0 = 100$ nM.

#### 4.2 Multichannel fluorescence experiments

Multichannel fluorescent experiments include both assays of a single pair of dye-labeled monomers and their complementary template and assays of sequence-specific dimerization (Supplementary Tables 11 and 12). The data from both classes of experiments were processed in the same way, with only the final fitting to infer concentrations being distinct. In all cases, fittings were performed with the MATLAB R2019a Optimization Toolbox.[2]

In each experiment, a number of fluorescence/FRET channels (ranging from 3 to 6) were monitored. Let $F_i^a$ be the signal observed in channel $a$ at timepoint $i$ of 'Measurement 3', having subtracted the buffer-only negative control. As in Supplementary Section 4.1, we rescale according to $F_i'^{\alpha} = (F_1^{\alpha+}/F_i^{\alpha+})F_i^{\alpha}$, where $F_1^{\alpha+}$ is the positive control signal recorded during 'Measurement 1' in channel $a$ and $F_i^{\alpha+}$ being the positive control at timepoint $i$ during 'Measurement 3'.

In all cases, we consider the increase in fluorescence relative to the value measured during 'Measurement 2', $F_2'^{\alpha}$, as entirely resultant from the injection of labeled $N_y$ and conversion of $M_xL$ and $N_y$ into products $M_xN_y$; as in Supplementary Note 4.1 the contribution of complexes involving the template is assumed to be constant during 'Measurement 3' (and equal to the value obtained during 'Measurement 2'). Thus within channel $\alpha$

$$\Delta F_i^{\prime,\alpha} = \sum_y \left( [N_y]_0 - \sum_x [M_xN_y]_i \right) a_N^{\alpha} + \sum_{x,y} [M_xN_y]_i \left( a_{M_xN_y}^{\alpha} - \alpha_{M_xL}^{a} \right), \qquad (13)$$

where $\Delta F_i^{\prime,\alpha} = F_i^{\prime,\alpha} - F_2^{\prime,\alpha}$, $a_Z^{\alpha}$ is the coefficient of fluorescence of species $Z$ in channel $\alpha$, as collected in Supplementary Table 6. We estimate $[M_xN_y]_i$ at each timepoint $i$ by fitting Supplementary Equation 13 to the data.

##### 4.2.1 Leak and dimerization experiments with labeled $N_1$ or $N_2$

During these assays, up to three fluorescent channels were monitored, corresponding to the fluorophore on $M_x$, the fluorophore on $N_y$, and the FRET channel for $M_xN_y$ (for $M_2N_2$ and $M_3N_1$, the FRET channel doesn't exist as the fluorophores are identical). With a single monomer of each type, Supplementary Equation 13 reduces to

$$\Delta F_i^{\prime,\alpha} = ([N_y]_0 - [M_xN_y]_i)\, a_N^{\alpha} + [M_xN_y]_i \left( a_{M_xN_y}^{\alpha} - \alpha_{M_xL}^{a} \right). \qquad (14)$$

$[M_xL]_0$ was inferred by dividing the signal obtained in its dominant channel during 'Measurement 1' by the fluorescence coefficient: $[M_xL]_0 = F_1^{\alpha}/a_{M_xL}^{\alpha}$. $[N_y]_0$ was inferred by assuming $[M_xN_y] = 0$ immediately after the injection that starts 'Measurement 3'. The increase in fluorescence of the first timepoint of 'Measurement 3' relative to 'Measurement 2' ($\Delta F_3^{\prime,\alpha}$)

in the channel most relevant to $N_3$ was then used to infer $[N_y]_0$ via $[N_y]_0 = (\Delta F_3^{\prime,\alpha})/a^{\alpha}_{N_y}$.

$[M_xN_y]$ at subsequent timepoints was inferred by fitting $[M_xN_y]_i$ to the observed values of $\Delta F_i^{\prime,\alpha}$ by minimizing the base-10 logarithm of the mean square error:

$$\log_{10}\left(\frac{\sum_{\alpha=1}^{3}(\Delta F^{\prime,\alpha}_{i,\text{meas}} - \Delta F^{\prime,\alpha}_{i,\text{pred}})^2}{w_\alpha}\right), \tag{15}$$

which quantifies the difference between the observed signal and the signal predicted using Supplementary Equation 14 in the various channels. Here, $\Delta F^{\prime,\alpha}_{i,\text{meas}}$ is the measured increase in fluorescence in channel $\alpha$ at time step $i$ relative to 'Measurement 2', and $\Delta F^{\prime,\alpha}_{i,\text{pred}}$ is the increase predicted using Supplementary Equation 14. To reduce the impact of calibration and assumption-originated errors, the fluorescence channels were weighted differently using a factor $w_\alpha$ to prioritize error reduction in the FRET-measuring channel signal. The fit was performed chronologically through the timepoints, with the initial condition for each timepoint being $[M_xN_y]_{i-1}$, the value obtained at the previous timepoint.

###### 4.2.2 Sequence-specific copying by templated dimerization

The procedure is similar to inferring the concentrations of a single product, except that all six FRET/fluorescence channels were monitored in each experiment, and the aim is to extract concentrations for all nine products $M_xN_y$. In practice, it is essentially impossible to infer $M_2N_2$ because $\alpha_{N_2} + \alpha_{M_2} \approx \alpha_{M_2N_2}$ in the channel in which it generates a strong signal. Formation of $M_3N_1$, which also contains two identical fluorophores (Alexa Fluor 647), however, results in a detectable change in fluorescence, albeit with a low signal-to-noise ratio. We therefore neglect the $[M_2N_2]$ terms in Supplementary Equation 13, and fit only the remaining eight values of $[M_xN_y]$ to the observed data.

Fitting proceeded as follows:

1. From the signal of 'Measurement 1' we estimated $[M_1L]_0$, $[M_2L]_0$ and $[M_3L]_0$ by fitting the signal of the 6 fluorescence channels, using as $a^{\alpha}_{M_xL}$ their corresponding values from the fluorescent coefficient calibration matrix (Supplementary Table 5 or 6). We fit

initial concentrations to the fluorescence data by minimizing:

$$\log_{10}\left(\frac{\sum_{\alpha=1}^{6}(F^{\alpha}_{1,\text{meas}} - F^{\alpha}_{1,\text{pred}})^2}{6}\right), \tag{16}$$

given $F^{\alpha}_{1,\text{pred}} = \sum_x [M_x L]_0 a^{\alpha}_{M_x L}$, with initial estimates for $[M_x L]_0$ given by their intended concentration. Due to the low signal of the quenched $[M_x L]_0$, the quantification of this species was more prone to error than other fluorescent species.

2. We inferred $[N_1]_0$ and $[N_2]_0$ from the first timepoint of 'Measurement 3' by minimizing

$$\log_{10}\left(\frac{\sum_{\alpha=1}^{6}(F'^{\alpha}_{3,\text{meas}} - \Delta F'^{\alpha}_{3,\text{pred}})^2}{6}\right), \tag{17}$$

given $\Delta F'^{,\alpha}_{3\text{pred}} = \sum_{y=1,2}[N_y]_0 a^{\alpha}_{N_y}$, and using the intended concentrations as initial conditions of the fit. Since $[N_3]_0$ is not labeled, but it was injected in a mix along the other two monomers, its concentration for each well was assumed to be the average of the fitted $[N_1]_0$ and $[N_2]_0$.

3. We iteratively fitted $[M_x N_y]_i$ at each successive timepoint $i$ to minimise

$$\log_{10}\left(\frac{\sum_{\alpha=1}^{6}(\Delta F'^{\alpha}_{i,\text{meas}} - \Delta F'^{\alpha}_{i,\text{pred}})^2}{6} + 5000\frac{\sum_{x,y\neq 2,2}([M_x N_y]_i - [M_x N_y]_{i-1})^2}{8}\right). \tag{18}$$

Here, $\Delta F'^{,\alpha}_{i,\text{meas}}$ is the measured increase in fluorescence in channel $\alpha$ at time step $i$ of 'Measurement 3' relative to 'Measurement 2', and $\Delta F'^{,\alpha}_{i,\text{pred}}$ is the increase predicted using Supplementary Equation 13, given the estimated $[M_x N_y]_i$ at that timepoint. $[M_x N_y]_{i-1}$ are the concentrations estimated at the previous time step. $[M_x N_y]_{i-1}$ was also used as the initial condition for the fitting of $[M_x N_y]_i$, except for the dimer that matches the template. The initial condition for that matching dimer was chosen by incrementing $[M_x N_y]_i$ sufficiently to match the increase in fluorescence in the most relevant fluorescence/FRET channel.

Since there are eight variables to fit six signals, a unique optimal fit cannot be guaranteed. However, using the iterative procedure outlined above that penalizes unphysical rapid concentration fluctuations, it was possible to obtain a well-behaved fit to the fluorescent signals.

### 5 Supplementary Note 5: Fitted models

We fit the molecular concentrations, inferred as described in Supplementary Note 4, to simple kinetic models.

Binding of $M_xL$ to $T_{xy}$. Binding of $M_xL$ to $T_{xy}$ (or displacement of the lock strand $L$ from $M_xL$ by $T_{xy}$) is fitted using an irreversible reaction:

$$\mathrm{M}_x\mathrm{L} + \mathrm{T_{xy}} \xrightarrow{\mathrm{k}_t} \mathrm{M}_x\mathrm{T_{xy}}. \tag{19}$$

The corresponding ODE is

$$\frac{d[M_xT_{xy}]}{dt} = k_t[M_xL][T_{xy}], \tag{20}$$

using initial conditions $[M_xT_{xy}]_0 = 0$ and $[M_xL]_0$ and $[T_{xy}]_0$ inferred from the data as outlined in Supplementary Note 4.

Leak reaction between free-in-solution $M_xL$ and $N_y$. Reaction between monomers in the absence of $T_{xy}$ is fitted using an irreversible reaction:

$$\mathrm{M}_x\mathrm{L} + \mathrm{N}_y \xrightarrow{\mathrm{k_{leak}}} \mathrm{M}_x\mathrm{N}_y. \tag{21}$$

The corresponding ODE is

$$\frac{d[M_xN_y]}{dt} = k_{\mathrm{leak}}[M_xL][N_y], \tag{22}$$

with initial conditions $[M_xN_y]_0 = 0$ and $[M_xL]_0$ and $[N_y]_0$ inferred from the data as outlined in Supplementary Note 4.

Handhold-mediated strand displacement of $T_{xy}$ by $N_y$. The displacement of $T_{xy}$ from the $M_xT_{xy}$ complex is fitted using an irreversible reaction:

$$\mathrm{M}_x\mathrm{T_{xy}} + \mathrm{N}_y \xrightarrow{\mathrm{k}_h} \mathrm{M}_x\mathrm{N}_y/\mathrm{T_{xy}}. \tag{23}$$

With '/' denoting the handhold-mediated binding of $M_xN_y$ to $T_{xy}$. The corresponding ODE is

$$\frac{d[M_xN_y/T_{xy}]}{dt} = k_h[M_xT_{xy}][N_y], \tag{24}$$

with initial conditions $[M_xN_y/T_{xy}]_0 = 0$ and $[M_xT_{xy}]_0$ and $[N_y]_0$ inferred from the data as outlined in Supplementary Note 4.

Initial turnover frequency for catalytic dimerisation of $M_xL$ and $N_y$ by $T_{xy}$. We assume that at short times (less than two hours for 10 nM $M_xL$-scale reactions and thirty minutes for 100 nM $M_xL$-scale reaction) the concentration of reacted $M_xL$ molecules (inferred as outlined in Supplementary Note 4) is linear with time

$$[M_xL]_{\text{reacted}} = V_0t + c. \tag{25}$$

Here, $V_0$ is the initial rate of the reaction, which is divided by the intended amount of template in the reaction to calculate the turnover frequency (TOF).

### 6 Supplementary Note 6: General fitting procedure for the estimation of rate constants

All fits were performed, using MATLAB® R2022b 'Optimization Toolbox', on the data transformed as described in Supplementary Section 4.

#### 6.1 Fits to obtain $k_t$, $k_h$ and $k_{\text{leak}}$

Data from individual reactions were fitted to the analytical solutions of the ODE models (Supplementary Equations 19, 23 and 21),

$$[X]_t = [\text{Trig}]_0 \left( \frac{1 - e^{k(t+t_0)([\text{Trig}]_0 - [X^*]_0)}}{1 - \frac{[Trig.]_0}{[X^*]_0} e^{k(t+t_0)([\text{Trig}]_0 - [X^*]_0)}} \right), \tag{26}$$

with $X$ being the reaction-produced fluorescence species, $X^*$ the initially quenched species, $k$ the 2<sup>nd</sup> order kinetic rate of the reaction and Trig the species at the lower concentration used to trigger the reaction, e.g., $N_{xy}$ in HMSD reactions. We used the minimization function *fminsearch* to minimize the base-10 logarithm of the weighted mean squared errors of the fitting

$$\log_{10}(\text{ mean weighted error }) = \log_{10} \frac{\sum_t \left([X]_{\text{exp}}(t) - [X]_{\text{model}}(t)\right)^2}{[X]_{\text{model}}(t)}, \tag{27}$$

where $[X]_{\text{exp}}(t)$ is the inferred concentration of product at time $t$ of the fluorescent trace, and $[X]_{\text{model}}(t)$ is the total concentration predicted by the model at time $t$. The error was weighted at each point with $[X]_{\text{model}}(t)$ to account for the Poisson distribution of the photomultiplier detection error.

We treated the rate constant $k$ as the main fitting parameter. The initial guess for $k$ ($k_{\text{init}}$) was inferred from the time at which the reaction reached 90% completion:

$$k_{\text{init}} = \frac{\ln\left(10 - 9\frac{[\text{Trig.}]_0}{[X]_0}\right)}{t_{90\%}\left([X]_0 - [\text{Trig}]_0\right)}. \tag{28}$$

In addition, in the fits to Supplementary Equation 26, the initial measurement time ($t_0$) and $[\text{Trig}]_0$ were treated as secondary parameters to be fitted. The initial guess for $t_0$ ($t_{0\_\text{init}}$)

was obtained from the cutting point with the x-axis of projection of the initial rate of the reaction. This projection was a linear fit calculated using initial data from before the reaction had reached 50% completion, where the trajectories are in a pseudo-linear regime.

Approximate values of $[\mathrm{Trig}]_0$ were inferred as side products from the processing of experimental data outlined in Supplementary Notes 3 and 4, typically using the steady-state plateau in 'Measurement 4'. These approximate values were then used as initial conditions for fitting $[\mathrm{Trig}]_0$ as part of identifying rate constants $k$. For kinetics that did not reach at least 90% completion, the trigger's concentration was only allowed to fluctuate ± 10% from the value inferred from 'Measurement 4'. The percentage of reaction completion was determined by comparing the last value of the recorded kinetics in 'Measurement 3' with 'Measurement 4'. Consequently, for leak reactions, which never reach completion in the measured time, $[N_y]_0$ is always only allowed to fluctuate ± 10%.

##### 6.1.1 Results for $M_xL$ binding to $T_{xy}$ (estimating $k_t$)

The fitted $k_t$ values are reported in Supplementary Table 13. Fitted trajectories are shown in Supplementary Figures 10 and 11. The inferred rate constants presented some variability between different tested $T_{xy}$ strands that shared a $t$ sequence. This difference could perhaps be explained by experimental error or the partial secondary structure of the $T_{xy}$ strands.

**Supplementary Table 13: Fitted TMSD rate constant $k_t$.** Data obtained using procedures described in Supplementary Table 7 and Supplementary Note 6. The reactant strands used, at 25° C, were all $T_{xy}$ strands with 8 nt $h$ and 6 nt primary toehold unless stated otherwise, (Supplementary Table 1) with their complementary $M_x$ (Supplementary Table 2). Mean of all fitted $k_t$ for six-nucleotide toehold variants of $M_1 = 9.1 \pm 0.3 \times 10^4$ M$^{-1}$ s$^{-1}$; for $M_2 = 4.3 \pm 0.3 \times 10^4$ M$^{-1}$ s$^{-1}$; for $M_3 = 2.33 \pm 0.12 \times 10^4$ M$^{-1}$s.$^{-1}$

| **Condition (x/y) ($u$t/$v$h)** | **Individual experiments $k_t$ x $10^5$ (M$^{-1}$ s$^{-1}$)** | | | | | | | | | **Mean $k_t$ x $10^5$ ± SEM (M$^{-1}$ s$^{-1}$)** |
|---|---|---|---|---|---|---|---|---|---|---|
| $T_{11}$ | 0.99 | 0.9 | 0.84 | 0.79 | 0.75 | 0.75 | 0.74 | 0.75 | - | 0.81 ± 0.03 |
| $T_{12}$ | 0.885 | 0.9 | 0.979 | 0.995 | 0.985 | 0.943 | 0.921 | 0.916 | 0.858 | 0.931 ± 0.016 |
| $T_{13}$ | 1.1 | 1.1 | 1.12 | 1.21 | 0.92 | 1.03 | 0.6 | 0.71 | 0.92 | 0.97 ± 0.07 |
| $T_{21}$ | 0.24 | 0.233 | 0.235 | 0.234 | 0.24 | 0.245 | 0.256 | 0.278 | 0.347 | 0.256 ± 0.012 |
| $T_{22}$ | 0.501 | 0.473 | 0.486 | 0.486 | 0.492 | 0.503 | 0.5 | 0.526 | 0.57 | 0.504 ± 0.01 |
| $T_{23}$ | 0.671 | 0.545 | 0.553 | 0.519 | 0.497 | 0.52 | 0.535 | 0.508 | 0.586 | 0.548 ± 0.018 |
| $T_{31}$ | 0.1674 | 0.1699 | 0.1729 | 0.1741 | 0.1756 | 0.1739 | 0.1758 | 0.1785 | 0.1829 | 0.1746 ± 0.0015 |
| $T_{32}$ | 0.3165 | 0.3082 | 0.3196 | 0.3105 | 0.3147 | 0.3151 | 0.3127 | 0.3144 | 0.3172 | 0.3143 ± 0.0012 |
| $T_{33}$ | 0.222 | 0.209 | 0.207 | 0.208 | 0.21 | 0.209 | 0.221 | 0.217 | 0.207 | 0.212 ± 0.002 |
| $T_{13}$ **(4t/8h)** | 0.0113 | 0.011 | 0.01148 | 0.0111 | - | - | - | - | - | 0.01122 ± 0.00011 |
| $T_{13}$ **(5t/8h)** | 0.0449 | 0.0444 | 0.0442 | 0.0456 | - | - | - | - | - | 0.0448 ± 0.0003 |
| $T_{13}$ **(7t/8h)** | 9.6 | 10.7 | 11.2 | 10.8 | 10.5 | 10.4 | - | - | - | 10.5 ± 0.2 |
| $T_{13}$ **(8t/8h)** | 13.7 | 15.4 | 18.5 | 16.4 | 16.1 | 16.1 | - | - | - | 16 ± 0.6 |

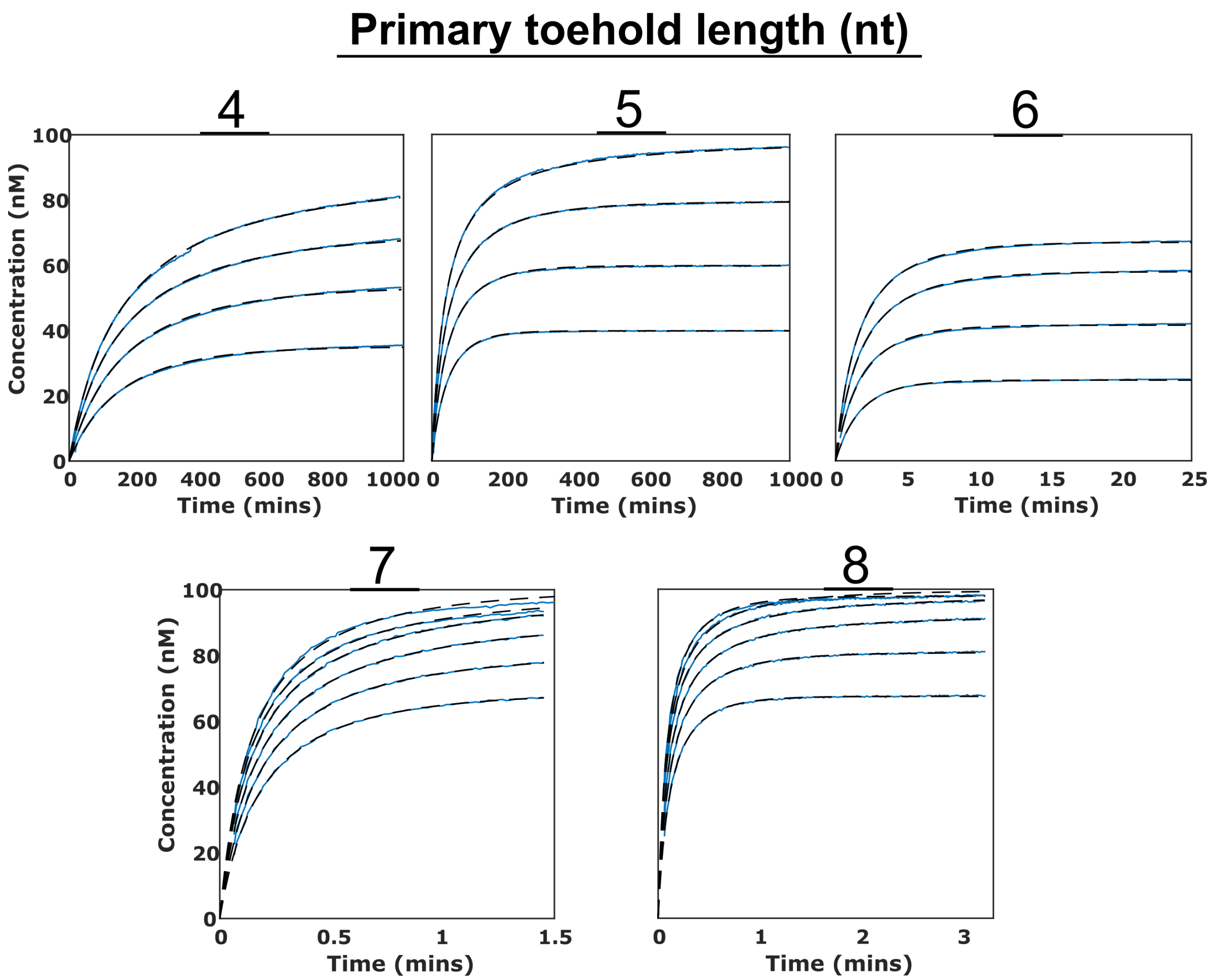


**Supplementary Figure 10: Fits of $k_t$ for different primary toehold lengths $t$.** The kinetics display the concentration of displaced $M_1L$ with time. Blue: Experimental data. Dashed line: Fitted model. Intended initial concentrations: $[M_1L]_0 = 100$ nM, $[T_{xy}]_0 =$ variable.

##### 6.1.2 Results for leak reaction rates $k_{leak}$ for $M_xL$ and $N_y$

Of these systems, only $M_1L$ with $N_3$ or $N_1$ were assayed with replicas as independent experiments. The rest of monomer combinations were individual controls accompanying their corresponding templated reactions. Due to the subtle fluorescence changes when producing $M_2N_2$ and $M_3N_1$ from their constituents, no leak signal could be detected and hence no $k_{\text{leak}}$ could be fitted. For fitting $k_{\text{leak}}$, the initial 1000 or 2000 minutes of the reaction were ignored to fit the representative kinetics at later time, and thus neglecting the initial leak due to malformed structures.

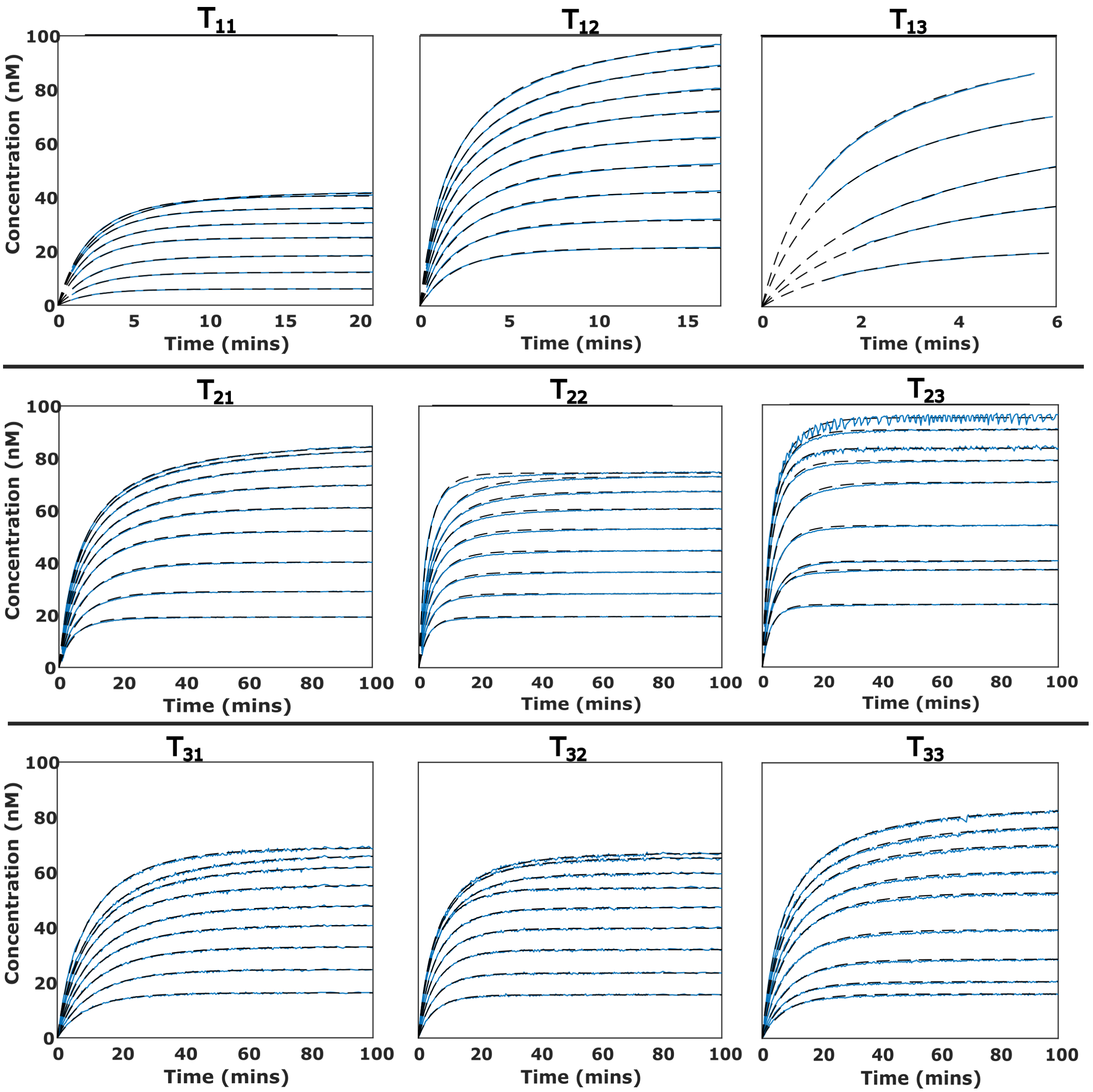


**Supplementary Figure 11: Fits of $k_t$ for different competing templates.** The kinetics display the concentration of displaced $M_xL$ with time. Blue: Experimental data. Dashed line: Fitted model. Intended concentrations: $[M_xL]_0$ = 100 nM, $[T_{xy}]_0$ = variable.

All fitted $k_{\text{leak}}$ are collected in Supplementary Table 14. Fitted trajectories are shown in Supplementary Figure 12. The obtained $k_{\text{leak}}$ values are very slow in comparison with the overall templated reaction, having a value that ranges from the $10^{-1}$ to 1 $M^{-1}$ $s^{-1}$. Additionally, $k_{\text{leak}}$ seems more dependent on $M_xL$ than $N_y$, with averages for each $M_x$ being: $M_1 = 0.32 \pm 0.05$ $M^{-1}$ $s^{-1}$; $M_2 = 1.1 \pm 0.2$ $M^{-1}$ $s^{-1}$; $M_3 = 0.56 \pm 0.013$ $M^{-1}$ $s^{-1}$.

**Supplementary Table 14: Fitted leak reaction rate $k_{leak}$.** Data obtained using procedures described in Supplementary Table 8 and Supplementary Note 6. The reactant complexes used were $M_xL$ (Supplementary Table 2) triggered by different $N_y$ strands (Supplementary Table 3). All reported experiments were performed at 25$^0$C. Mean of fitted $k_{\text{leak}}$ for $M_1L = 0.32 \pm 0.05$ M$^{-1}$ s$^{-1}$; $M_2L = 1.1 \pm 0.2$ M$^{-1}$ s$^{-1}$; $M_3L = 0.56 \pm 0.013$ M$^{-1}$ s$^{-1}$ .

| **Condition (x/y)** | **Individual experiments $k_{leak}$ (M$^{-1}$ s$^{-1}$)** | | | | | **Mean $k_{leak}$ ± SEM (M$^{-1}$ s$^{-1}$)** |
|---|---|---|---|---|---|---|
| $M_1N_1$ | 0.17 | 0.31 | 0.65 | 0.22 | - | 0.34 ± 0.1 |
| $M_1N_2$ | 0.25 | - | - | - | - | 0.25 |
| $M_1N_3$ | 0.22 | 0.25 | 0.32 | 0.56 | 0.27 | 0.32 ± 0.06 |
| $M_2N_1$ | 1.2 | - | - | - | - | 1.2 |
| $M_2N_3$ | 0.9 | - | - | - | - | 0.9 |
| $M_3N_2$ | 0.55 | - | - | - | - | 0.55 |
| $M_3N_3$ | 0.57 | - | - | - | - | 0.57 |

##### 6.1.3 Results for handhold-mediated strand displacement reaction rate $k_h$ for the reaction of $M_1T_{13}$ with $N_3$

All fitted $k_h$ are collected in Supplementary Table 15. Fitted trajectories are shown in Supplementary Figures 13 and 14. The fitted experiments had a lower signal-to-noise ratio and were faster than experiments that started with fully quenched monomers, leading to $k_h$ values with significant uncertainty. However, the results show a clear trend of increasing $k_h$ with handhold length. Additionally, several primary toehold lengths were tested with handhold $8h$, to test if the fluorescence recovery was correlated with the detachment of the product from $T_{xy}$. No importantt differences in fluorescence recovery were observed between experiments.

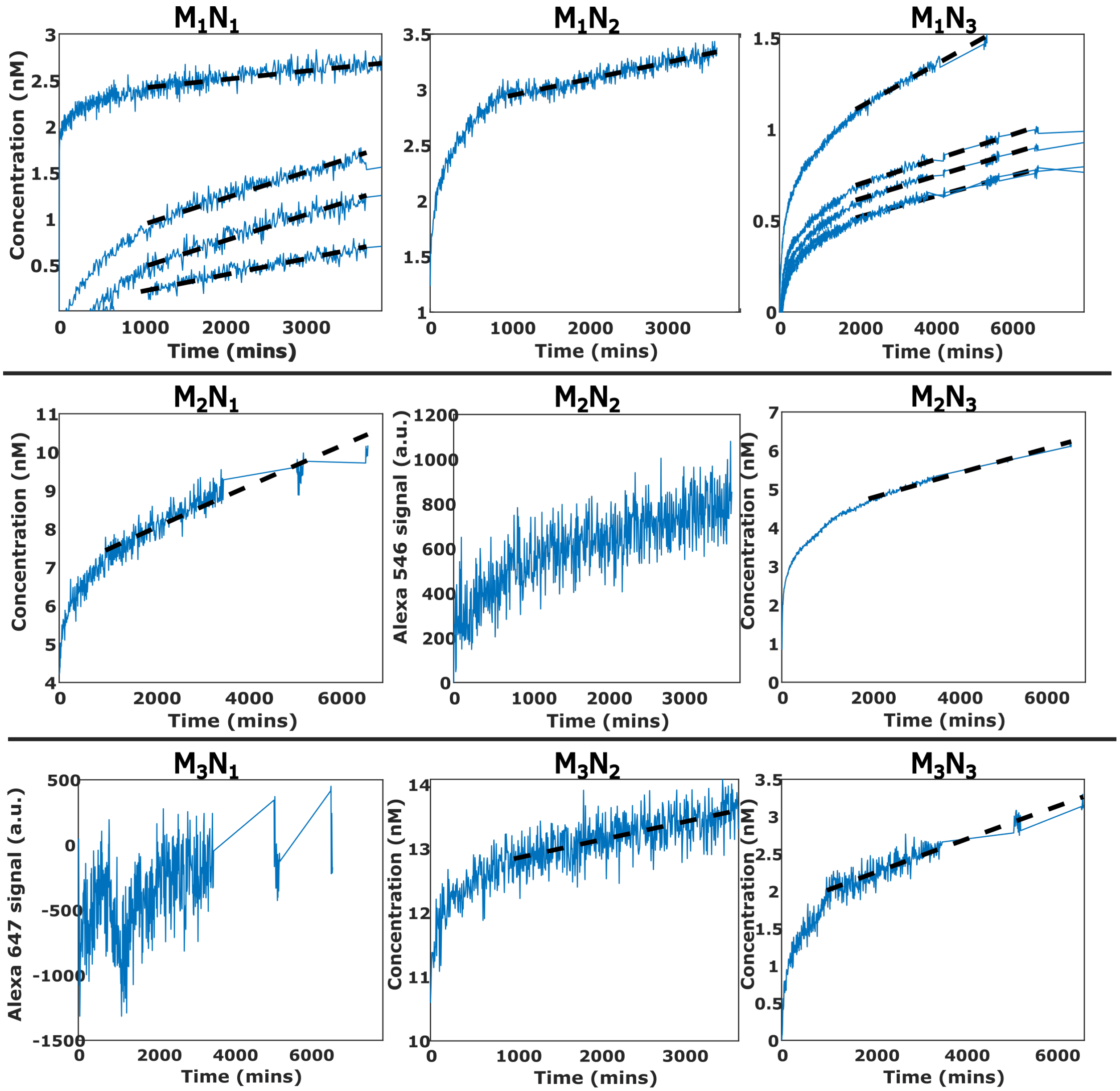


**Supplementary Figure 12: Fits of $k_{\text{leak}}$ for different combination of monomers.** Kinetics display the concentration of displaced $M_xL$ with time, except in $M_2N_2$ and $M_3N_1$ where the processed values of fluorescence are reported. Blue: Experimental data. Dashed line: Fitted model. Intended concentrations: $[M_xL]_0 = 100$ nM, $[N_y]_0 = 100$ nM (or 100, 80, 60 and 40 nM for $M_1N_3$). $M_1N_1$ concentrations were: 100/100, 100/100, 150/150 and 50/150 ($M_x/N_y$ nM).

**Supplementary Table 15: Fitted $k_h$.** Data obtained using procedures described in Supplementary Table 8 and Supplementary Note 6. The reactant complexes used were $M_1T_{13}$ (Supplementary Tables 1 and 2) triggered by $N_3$ strands (Supplementary Table 3). Fitting results with excessive error, which were not considered for the mean calculation in the final solumn, are shown in italics. Mean of fitted $k_h$ for the values considered reliable: 6 nt handhold: 5.9 ± 0.6 x $10^4$ $M^{-1}$ $s^{-1}$; 7 nt: 5.2 ± 0.4 x $10^4$ $M^{-1}$ $s^{-1}$; 8 nt: 8 ± 0.5 x $10^4$ $M^{-1}$ $s^{-1}$; 9 nt: 2.26 ± 0.1 x $10^5$ $M^{-1}$ $s^{-1}$; 10 nt: 5.6 ± 0.8 x $10^5$ $M^{-1}$ $s^{-1}$

| **Condition ($ut/vh$)** | **Individual experiments $k_h$ x $10^4$ ($M^{-1}$ $s^{-1}$)** | | | | | | **Mean $k_h$ x $10^4$ ± SEM ($M^{-1}$ $s^{-1}$)** |
|---|---|---|---|---|---|---|---|
| **6/6** | *16.3* | *15* | *12.3* | *10.3* | 6.5 | 5.3 | 5.9 ± 0.6 |
| **6/7** | *17.3* | *15.7* | *12.4* | 5.5 | 4.3 | 5.9 | 5.2 ± 0.4 |
| **4/8** | 3.2 | 3.58 | 3.3 | 3.75 | 4.06 | 3.86 | 3.63 ± 0.13 |
| **5/8** | 10.4 | 8.8 | 8.6 | 8.1 | 8.5 | 7.9 | 8.7 ± 0.4 |
| **6/8** | 7.14 | 7.28 | 6.74 | 6.82 | 7.41 | 7.86 | 7.21 ± 0.17 |
| **7/8** | 9.8 | 9.5 | 9.5 | 10.2 | 11.3 | 12.3 | 10.4 ± 0.5 |
| **8/8** | 9.7 | 9.3 | 10.2 | 11.3 | 11.2 | 6.9 | 9.8 ± 0.7 |
| **6/9** | 22.2 | 23.5 | 21.1 | 26.3 | 19 | 23.4 | 22.6 ± 1 |
| **6/10** | 64 | 66 | *10* | *17* | *26* | 40 | 56 ± 8 |

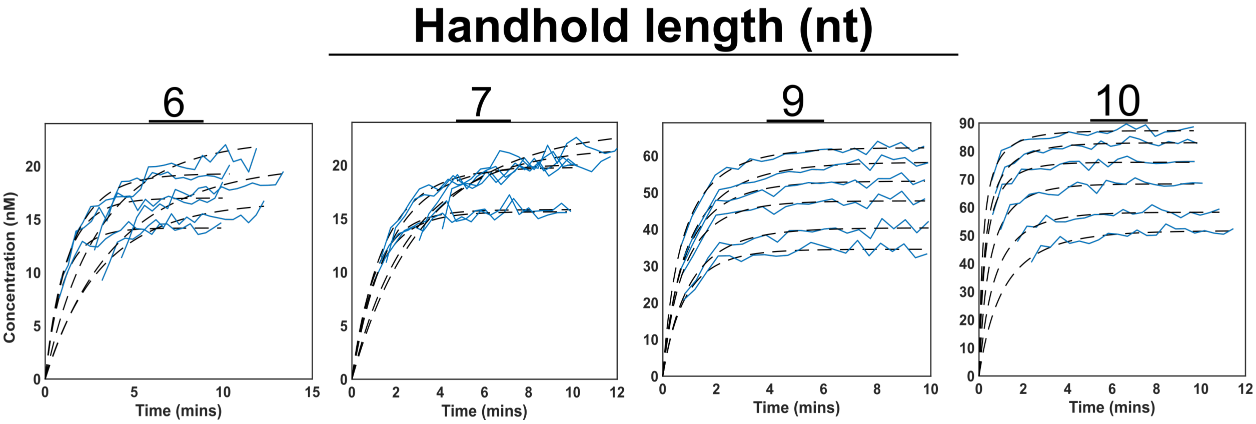


**Supplementary Figure 13: Fits of $k_h$ for different handhold lengths.** Kinetics display the concentration of formed $M_1N_3$ with time. Due to the low signal-to-noise ratio and swiftness of the reaction, most of the kinetics were almost completed when starting to read. Blue: Experimental data. Dashed line: Fitted model. Intended concentrations: $[M_1T_{13}]_0 = 100$ nM, variable concentration for $N_3$.

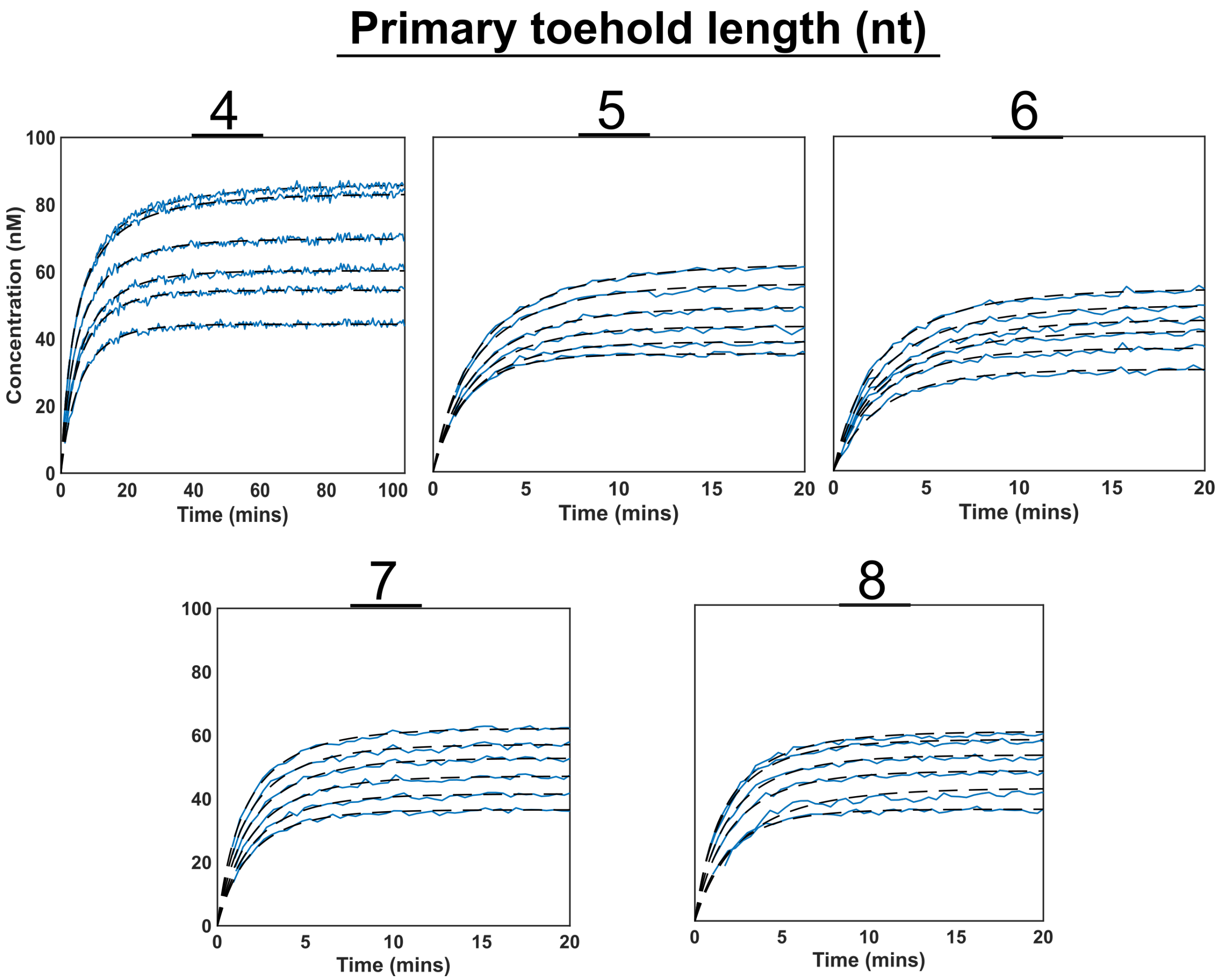


**Supplementary Figure 14: Primary toehold length doesn't affect the HMSD kinetics.** Fitting of 8 nt handhold experiments with different $t$ lengths. Kinetics display the concentration of formed $MN_xL$ with time. Blue: Experimental data. Dashed line: Fitted model. Intended concentrations: $[M_1T_{13}]_0 = 100$ nM, variable concentration for $N_3$.

#### 6.2 Fitting of turnover frequency for catalytic dimerization of $M_1L$ and $N_3$ mediated by $T_{13}$ with and without poduct inhibition

The first minutes of each individual reaction were fitted to Supplementary Equation 25, weighting each timepoint by the inverse of its fluorescence intensity. In this section, we report the slope of the regression line, which corresponds to the reactions' initial rate, with a confidence interval of 95%.

All fitted initial rates are collected in Supplementary Table 16. Complete trajectories are shown in Supplementary Figure 15. From the reported results, an $IC_{50}$ was estimated for each condition by simple linear interpolation between the two points nearest to half the initial rate of the reactions at $[M_1N_3]_0$=0. For example, $IC_{50}$(6t/6h) $\approx$ 70 nM, $IC_{50}$(6t7h) $\approx$ 50 nM, $IC_{50}$(6t8h) $\approx$ 11 nM, $IC_{50}$(6t9h) $\approx$ 4 nM, and $IC_{50}$(6t10h) $\approx$ 2 nM.

**Supplementary Table 16: Initial rates of monomer turnover by $T_{13}$ in the presence of $M_1N_3$ (10 nM-scale).** Data obtained using procedures described in Supplementary Table 16 and Supplementary Note 6. The reactant complexes used were $M_1L$ (Supplementary Table 1) triggered by $N_3$ strands (Supplementary Table 3) in the presence of different $T_{13}$ strands (Supplementary Table 1). Intended concentrations: $[M_1L]_0$ = 10 nM, $[N_3]_0$ 10 nM, $[T_{13}]_0$ = 1 nM, $[M_1N_3]_0$ = variable. Fitting results are reported in nM hour$^{-1}$, with a 95% confidence interval.

| **Condition ($ut/vh$)** | $[M_1N_3]_0$ **0 nM** | $[M_1N_3]_0$ **2.5 nM** | $[M_1N_3]_0$ **5 nM** | $[M_1N_3]_0$ **10 nM** | $[M_1N_3]_0$ **15 nM** | $[M_1N_3]_0$ **25 nM** | $[M_1N_3]_0$ **50 nM** | $[M_1N_3]_0$ **100 nM** |
|---|---|---|---|---|---|---|---|---|
| $T_{13}$ **(4t/6h)** | 0.024±0.004 | 0.017±0.007 | 0.016±0.005 | 0.021±0.005 | 0.023±0.004 | 0.018±0.004 | 0.021±0.005 | 0.021±0.005 |
| $T_{13}$ **(4t/7h)** | 0.025±0.005 | 0.024±0.004 | 0.021±0.005 | 0.021±0.003 | 0.02±0.004 | 0.025±0.005 | 0.021±0.005 | 0.02±0.006 |
| $T_{13}$ **(4t/8h)** | 0.046±0.006 | 0.049±0.008 | 0.056±0.008 | 0.043±0.005 | 0.054±0.009 | 0.05±0.008 | 0.051±0.008 | 0.036±0.007 |
| $T_{13}$ **(4t/9h)** | 0.043±0.007 | 0.037±0.009 | 0.035±0.008 | 0.033±0.006 | 0.029±0.007 | 0.022±0.006 | 0.061±0.014 | 0.014±0.006 |
| $T_{13}$ **(4t/10h)** | 0.05±0.007 | 0.037±0.007 | 0.035±0.007 | 0.032±0.005 | 0.033±0.006 | 0.028±0.005 | 0.017±0.009 | 0.024±0.005 |
| $T_{13}$ **(5t/6h)** | 0.027±0.004 | 0.027±0.005 | 0.026±0.005 | 0.022±0.006 | 0.018±0.005 | 0.021±0.004 | 0.021±0.004 | 0.02±0.006 |
| $T_{13}$ **(5t/7h)** | 0.028±0.005 | 0.026±0.004 | 0.029±0.005 | 0.028±0.003 | 0.028±0.005 | 0.024±0.004 | 0.026±0.005 | 0.018±0.004 |
| $T_{13}$ **(5t/8h)** | 0.134±0.012 | 0.119±0.012 | 0.129±0.014 | 0.108±0.013 | 0.099±0.01 | 0.099±0.013 | 0.089±0.012 | 0.081±0.011 |
| $T_{13}$ **(5t/9h)** | 0.105±0.008 | 0.081±0.01 | 0.082±0.01 | 0.069±0.009 | 0.049±0.009 | 0.058±0.009 | 0.031±0.008 | 0.029±0.012 |
| $T_{13}$ **(5t/10h)** | 0.092±0.006 | 0.065±0.005 | 0.062±0.006 | 0.042±0.007 | 0.043±0.006 | 0.038±0.007 | 0.032±0.007 | 0.034±0.01 |
| $T_{13}$ **(6t/6h)** | 0.063±0.004 | 0.063±0.005 | 0.056±0.004 | 0.05±0.005 | 0.049±0.006 | 0.044±0.005 | 0.037±0.005 | 0.022±0.006 |
| $T_{13}$ **(6t/7h)** | 0.096±0.005 | 0.082±0.004 | 0.08±0.003 | 0.074±0.005 | 0.066±0.004 | 0.063±0.005 | 0.046±0.005 | 0.019±0.005 |
| $T_{13}$ **(6t/8h)** | 0.622±0.009 | 0.493±0.008 | 0.417±0.009 | 0.322±0.011 | 0.261±0.013 | 0.223±0.012 | 0.155±0.014 | 0.058±0.007 |
| $T_{13}$ **(6t/9h)** | 1.01±0.02 | 0.615±0.015 | 0.453±0.013 | 0.307±0.008 | 0.208±0.006 | 0.161±0.009 | 0.09±0.009 | 0.067±0.009 |
| $T_{13}$ **(6t/10h)** | 0.369±0.013 | 0.156±0.01 | 0.101±0.007 | 0.071±0.006 | 0.064±0.011 | 0.048±0.008 | 0.043±0.01 | 0.041±0.01 |
| $T_{13}$ **(7t/6h)** | 0.058±0.003 | 0.056±0.003 | 0.056±0.004 | 0.051±0.003 | 0.048±0.003 | 0.052±0.005 | 0.045±0.003 | 0.04±0.003 |
| $T_{13}$ **(7t/7h)** | 0.172±0.004 | 0.146±0.003 | 0.129±0.004 | 0.114±0.003 | 0.098±0.004 | 0.091±0.004 | 0.075±0.003 | 0.065±0.003 |
| $T_{13}$ **(7t/8h)** | 0.432±0.006 | 0.338±0.005 | 0.281±0.005 | 0.231±0.004 | 0.19±0.004 | 0.168±0.003 | 0.11±0.003 | 0.094±0.004 |
| $T_{13}$ **(7t/9h)** | 0.272±0.008 | 0.208±0.006 | 0.167±0.005 | 0.133±0.004 | 0.112±0.005 | 0.091±0.005 | 0.066±0.003 | 0.052±0.003 |
| $T_{13}$ **(7t/10h)** | 0.09±0.004 | 0.067±0.004 | 0.06±0.004 | 0.046±0.004 | 0.048±0.004 | 0.041±0.005 | 0.038±0.005 | 0.034±0.004 |
| $T_{13}$ **(8t/6h)** | 0.054±0.004 | 0.047±0.003 | 0.053±0.003 | 0.047±0.003 | 0.049±0.004 | 0.045±0.003 | 0.038±0.004 | 0.04±0.003 |
| $T_{13}$ **(8t/7h)** | 0.16±0.004 | 0.143±0.005 | 0.122±0.003 | 0.106±0.004 | 0.092±0.004 | 0.086±0.004 | 0.068±0.005 | 0.058±0.004 |
| $T_{13}$ **(8t/8h)** | 0.194±0.004 | 0.15±0.004 | 0.119±0.004 | 0.102±0.004 | 0.086±0.004 | 0.072±0.004 | 0.061±0.004 | 0.052±0.003 |
| $T_{13}$ **(8t/9h)** | 0.114±0.005 | 0.09±0.004 | 0.066±0.004 | 0.051±0.004 | 0.047±0.004 | 0.044±0.004 | 0.032±0.004 | 0.029±0.004 |
| $T_{13}$ **(8t/10h)** | 0.043±0.004 | 0.037±0.003 | 0.037±0.003 | 0.035±0.004 | 0.03±0.004 | 0.027±0.005 | 0.028±0.004 | 0.036±0.005 |

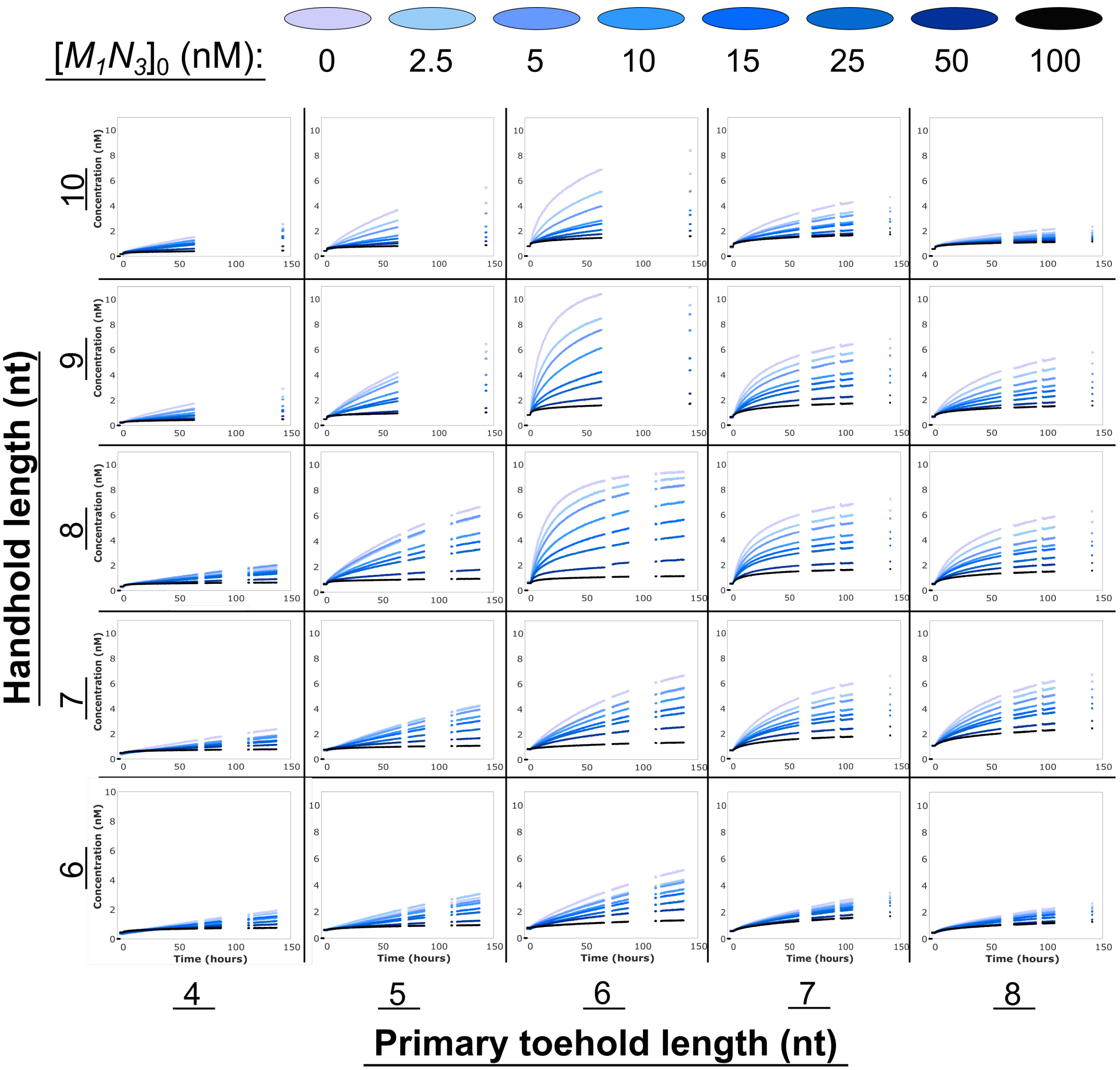


**Supplementary Figure 15: Kinetics of 10 nm-scale inhibition tests.** Reacted monomer concentration $[M_1L]$ in a system with 10 nM $N_3$, 10 nM $M_1L$, and 1 nM $T_{13}$, with variable lengths of $t$ and $h$ domains, and an initial non-fluorescent pool of products $M_1N_3$ at a range of concentrations $[M_1N_3]_0$.

##### 6.2.1 Results for $M_1L$ and $N_3$ dimerisation in the presence of $M_1N_3$, at high initial concentrations of substrate and product

To demonstrate the monomer-competitive nature of the observed product inhibition, inhibition testing was repeated, for different concentrations of $T_{13}$ (6t/8h), and the monomers at an intended concentration of 100 nM. All fitted initial rates for inhibition experiments are collected in Supplementary Table 17. Complete trajectories are shown in Supplementary Figure 16. The obtained results demonstrate that, for the three $[T_{13}]_0$ tested (2.5, 5 and 10 nM), the addition of $[M_1N_3]_0$ produced a similar reduction in their TOF. Additionally, for this new concentration regime, the estimated $IC50$ for all template concentrations is $[M_1N_3]_0 \approx [M_1L]_0$.

**Supplementary Table 17: Initial rates of monomer turnover by $T_{13}$ in the presence of $M_1N_3$ (100 nM-scale).** Data obtained using procedures described in Supplementary Table 16 and Supplementary Note 6. The reactant complexes used were $M_1L$ (Supplementary Table 1) triggered by $N_3$ strands (Supplementary Table 3) in the presence of different concentrations of $T_{13}$ (6t/8h) (Supplementary Table 1). Intended concentrations: $[M_1L]_0 = 100$ nM, $[N_3]_0$ 100 nM, $[T_{13}]_0 =$ variable, $[M_1N_3]_0 =$ variable. For easier comparison between different $[T_{13}]_0$ conditions, fitting results are reported as TOF, in hour$^{-1}$, with a 95% confidence interval.

| **Template Concentration** | $[M_1N_3]_0$ **0 nM** | $[M_1N_3]_0$ **50 nM** | $[M_1N_3]_0$ **100 nM** | $[M_1N_3]_0$ **200 nM** | $[M_1N_3]_0$ **300 nM** |
|---|---|---|---|---|---|
| 2.5 nM | 4.5 ± 0.2 | 2.62 ± 0.12 | 1.42 ± 0.08 | 1.18 ± 0.07 | 0.94 ± 0.07 |
| 5 nM | 4.5 ± 0.2 | 2.61 ± 0.09 | 2.07 ± 0.06 | 1 ± 0.03 | 0.77 ± 0.05 |
| 10 nM | 3.4 ± 1.2 | 2 ± 1 | 1.3 ± 0.7 | 0.8 ± 0.6 | 0.7 ± 0.6 |

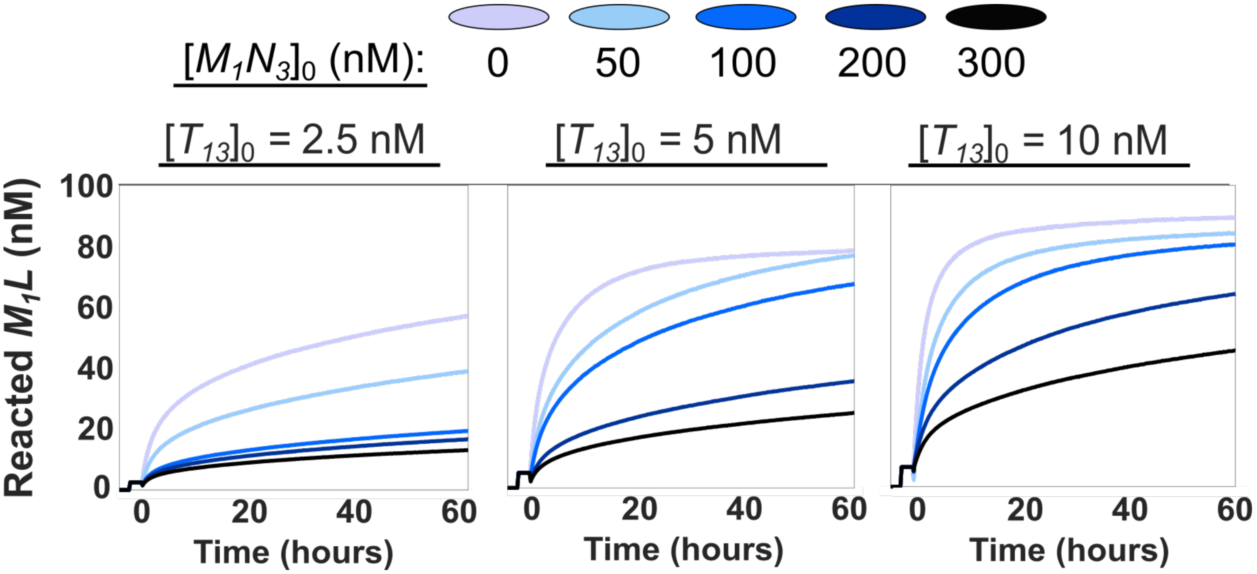


**Supplementary Figure 16: Kinetics of 100 nM-scale inhibition tests.** Reacted monomer concentration $[M_1L]$ in a system with 100 nM $N_3$, 100 nM $M_1L$, variable $T_{13}$ (6t/8h), and an initial non-fluorescent pool of products $M_1N_3$ at a range of concentrations $[M_1N_3]_0$. Results demonstrate that the proportion of TOF reduction is largely determined by the ratio of monomers/products, demonstrating the competitive nature of the product inhibition in the system.

##### 6.2.2 Characterisation of specific dimerization templates

All the different templates present in the selective dimerization assays were tested individually to confirm that they catalyzed formation of their intended products with similar rates between templates. Fitted initial rates for each template at different concentrations are collected in Supplementary Table 18. Complete trajectories are shown in Supplementary Figure 17. The initial rate of catalytic templating by $T_{31}$ was obtained by fitting for 180 minutes instead of 30 as usual, due to the low signal-to-noise ratio of the condition. Supplementary Figure 18 collects the fittings demonstrating the linear relation of initial rate and template concentration for all the different tested templates. Obtained results correlate with the $k_t$ values obtained individually for each $M_xL$ complex, with $T_{1y}$ having an overall higher TOF. All templates, except $T_{22}$, were capable of producing a discernible signal during the kinetics. The catalytic acticvity of $T_{22}$ was alternatively confirmed through PAGE electrophoresis (Figure 4b).

**Supplementary Table 18: Initial rates for monomer turnover by $T_{xy}$ used during specific dimerization assays.** Data obtained using procedures described in Supplementary Table 9 and Supplementary Note 6. The reactant complexes used were $M_xL$ (Supplementary Table 9) triggered by $N_y$ strands (Supplementary Table 3) in the presence of their complementary $T_{xy}$ strand (Supplementary Table 1). Intended concentrations: $[M_1L]_0$ = 100 nM, $[N_3]_0$= 100 nM, and $[T_{13}]_0$ = variable. Fitting results are reported in nM hour$^{-1}$, with a 95% confidence interval. For each $T_{xy}$ the TOF was estimated by a linear regression represented in Supplementary Figure 18. The results of two replicas were used to calculate the $T_{13}$ TOF (Figure 3d).

| **Template Condition** | $[T_{xy}]_0$ **5 nM** | $[T_{xy}]_0$ **3.75 nM** | $[T_{xy}]_0$ **2.5 nM** | $[T_{xy}]_0$ **1.25 nM** | $[T_{xy}]_0$ **0.5 nM** | $[T_{xy}]_0$ **0.25 nM** | **Fitted TOF (hour$^{-1}$)** |
|---|---|---|---|---|---|---|---|
| $T_{11}$ | 11.5 ± 0.6 | 8.8 ± 0.5 | 5.6 ± 0.4 | 3.2 ± 0.2 | 1.3 ± 0.4 | 0.9 ± 0.2 | 2,24 ± 0.11 |
| $T_{12}$ | 6.6 ± 0.6 | 5.1 ± 0.5 | 3.2 ± 0.2 | 2.3 ± 0.5 | 1.3 ± 0.3 | 1.01 ± 0.19 | 1.17 ± 0.11 |
| $T_{13}$ | 19.7 ± 0.9 | 14.9 ± 0.6 | 9.9 ± 0.5 | 6.4 ± 0.3 | 2.79 ± 0.18 | 1.17 ± 0.08 | 3.6 ± 0.3 |
| $T_{21}$ | 12 ± 3 | 9 ± 2 | 4 ± 3 | 3.4 ± 0.7 | 2.1 ± 0.7 | 1.7 ± 1.3 | 2.1 ± 0.5 |
| $T_{22}$ | 0 | 0 | 0 | 0 | 0 | 0 | |
| $T_{23}$ | 18 ± 2 | 12.1 ± 1.4 | 8 ± 1.5 | 3.8 ± 0.6 | 3.4 ± 1.5 | 3.3 ± 1.2 | 3 ± 1 |
| $T_{31}$ | 3.9 ± 0.3 | 3.1 ± 0.4 | 1.42 ± 0.08 | 0 | 0 | 0 | 1 ± 1 |
| $T_{32}$ | 6.3 ± 0.9 | 4.4 ± 1.2 | 4.5 ± 0.8 | 2.1 ± 1.4 | 1.9 ± 1.2 | 1.3 ± 1 | 1 ± 0.4 |
| $T_{33}$ | 11.5 ± 1.3 | 10.9 ± 1.4 | 7.5 ± 0.8 | 4.3 ± 0.8 | 2.2 ± 0.4 | 2.2 ± 0.5 | 2.2 ± 0.5 |
| **Template Condition** | $[T_{xy}]_0$ **5 nM** | $[T_{xy}]_0$ **2.5 nM** | $[T_{xy}]_0$ **1 nM** | $[T_{xy}]_0$ **0.5 nM** | | | **Fitted TOF (hour$^{-1}$)** |
| $T_{13}$ (replica) | 18.6 ± 0.3 | 10.03 ± 0.14 | 3.85 ± 0.08 | 2.12 ± 0.06 | | | Values considered for $T_{13}$ TOF calculation above |

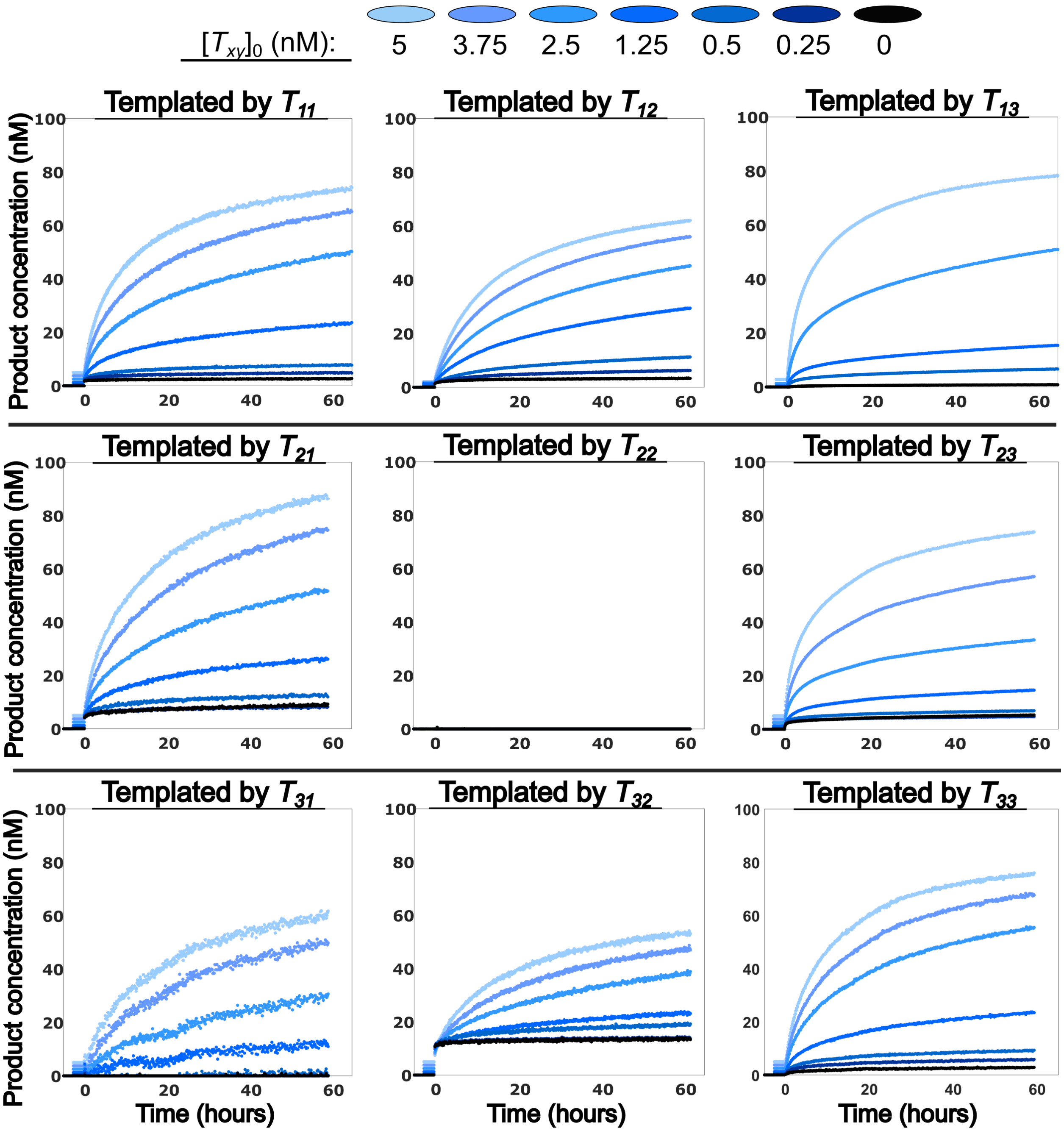


**Supplementary Figure 17: Kinetics of the $T_{xy}$-mediated reaction in isolation.** Formation of $M_xN_y$ product in a system with 100 nM of a single $N_y$, 100 nM of a single $M_xL$, and variable $[T_{xy}]_0$ as indicated by the trace color (5 nM, 3.75 nM, 2.5nM, 1.25 nM, 0.5nM and 0.25 nM). The $T_{13}$ panel contains only the kinetic traces not included in Figure 3c. The initial jump for conditions reading Green/Red FRET ($T_{21}$ and $T_{32}$) may be the result of calibration errors for species in that fluorescence channel. $T_{22}$ does not produe a detectable signal change upon dimerization.

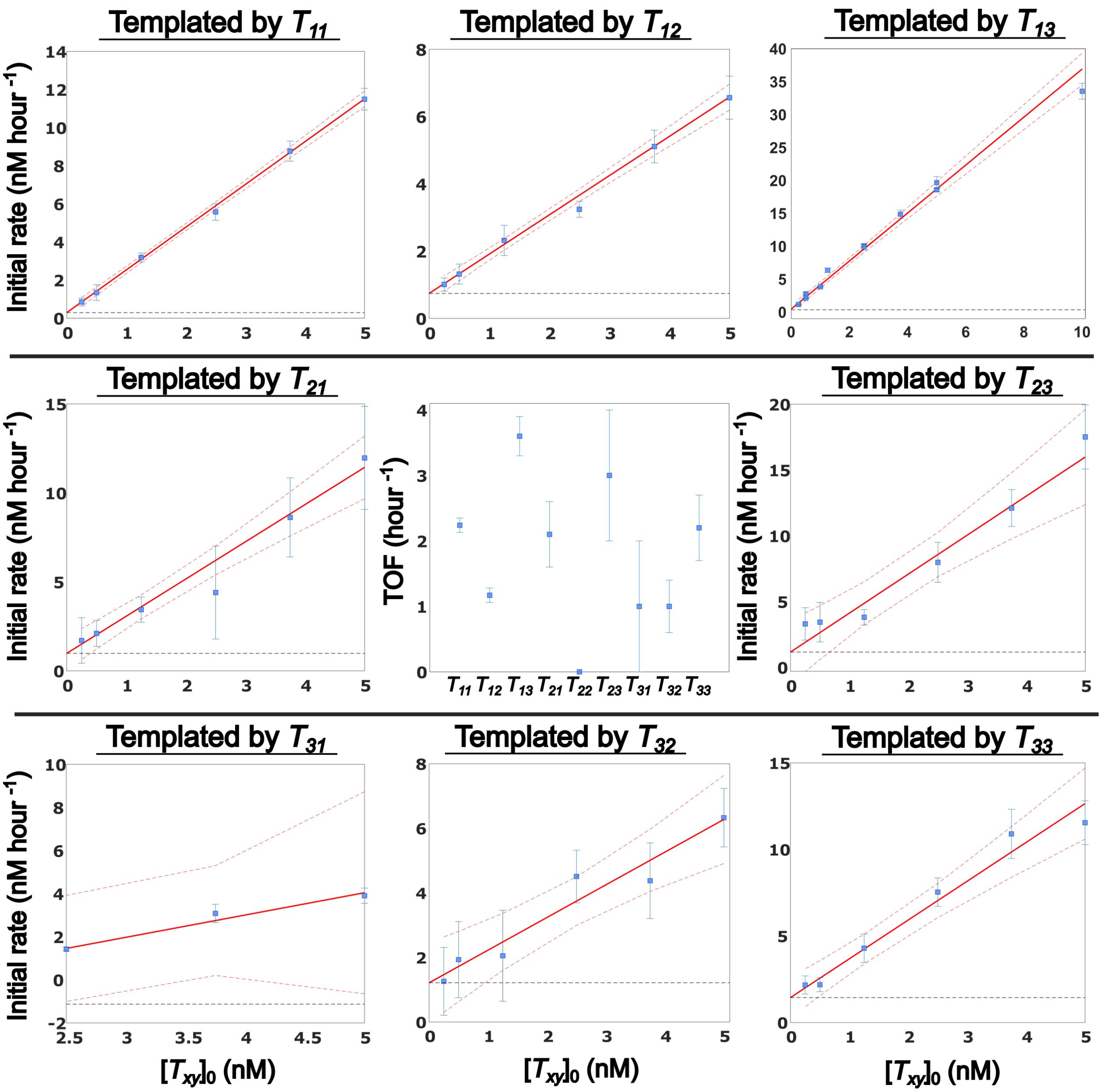


**Supplementary Figure 18: Linear fits of sequence-specific dimerization templates' initial rates to obtain their TOF.** Initial rates reported in Supplementary Table 18 were fitted to a linear model to calculate their TOFs (represented with a 95% confidence interval in the central panel). Blue: Experimental data; red line: Best linear fit with each data point inversely weighted by its error; red dashed line: Fit's 95% confidence interval; black dashed line: Fit estimated initial rate in the absence of template.

### 7 Supplementary Note 7: Putative design for longer templates

In Fig. 19, we illustrate how an HMSD-based system could – in principle – combat product inhibition for longer templates. Consider a template extended by adding more sites that look like the binding site of $M_x$ in this work prior to a final truncated binding site analogous to the binding site for monomer $N_y$ in this work. Monomers capable of recognizing these intermediate sites would have two polymerization domains, one in the forward direction and one in the backward direction, enabling them to form a long polymer.

In such a system, monomers binding to all but the final site of the template would bind stably until they have formed a polymerization bond in the forward's direction, just like $M_x$ binds stably to the template until dimerized with $N_y$ in the current work. Only the last monomer (the analog of $N_y$ in this paper) would be truncated to allow spontaneous detachment of the whole product after it is incorporated. Thus, HMSD could ,in principle, be used to overcome the worst consequences of cooperative product inhibition whilst allowing partially-formed copies to remain stably attached to the template. Crucial to this functionality is the manner in which HMSD channels free energy from the dimerization reaction into destabilizing the binding of one of the target monomers to the template. Recent theoretical work has demonstrated that a system with these properties is, *a priori*, capable of producing copies of longer templates with surprising efficacy in wide regions of parameter space.[5]

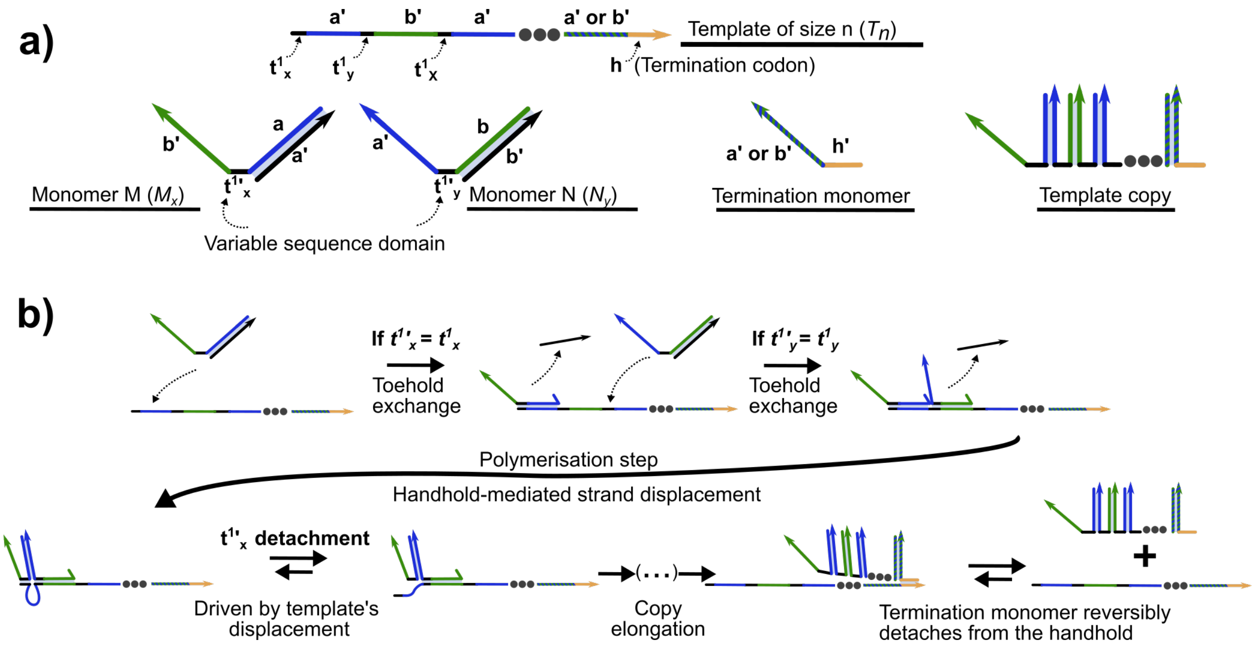


**Supplementary Figure 19: Outline design of a system that exploits HMSD to copy a longer template**. a) A long template consists of an alternating sequence of information-bearing toehold recognition domains and dimerization (or, in this case, polymerization) domains. Note that the dimerization domains themselves alternate (blue/green/blue...), to prevent self-interactions within the monomers. The final site of the template contains a truncated domain, which would play the role of a termination codon. The overall reaction would polymerize monomers of two different types (with interchanged polymerization domains) into a copy polymer with an information-bearing sequence of toehold recognition domains (shown in black here) and alternating polymerization duplexes. A truncated "termination" monomer caps the product. b) Detailed mechanism, showing iterated TMSD and HMSD reactions. Just as in the case of the dimer, HMSD would allow the product to bind only weakly to the template once the final termination monomer is incorporated. Note that, according to Juritz *et al.*,[5] such a system would actually operate best if multiple copies are bound to the template at the same time. We do not illustrate such a state for simplicity.

### 8 Supplementary Results 1: Results for alternate branch migration domain design

In this note we report a set of the experiments performed with a prototype of the dimerization mechanism, using the strands named 'v.1' (Supplementary Tables 1 to 3). All experiments described in this section followed the protocols described in Supplementary Note 3 for 'single fluorescence channel experiments'. Results were processed and fitted as described in Supplementary Sections 4.1 and 6.

The first tested version of the dimerization mechanism had strands with a branch migration domain of 28 nt and no clamps (Supplementary Figure 20a). Experiments with this version of the system proved successful as the dimerization reaction rate was completely dependent on template concentration for a variety of templates (results not included). The TOF of the template was also significantly higher than the final version of the system. In Supplementary Figure 20b we show one of the conditions tested for this prototype ($T_{xy}$(6t/8h)) which exhibits an initial TOF of 20 ± 4 hour$^{-1}$, calculated by fitting Supplementary Equation 25 to the first 10 minutes of the trajectories (Supplementary Figure 20c). This calculated initial TOF, for a monomer concentration of 60 nM, is faster than the 3.6± 0.3 hour$^{-1}$ resulting from the final system with the same toehold and handhold domain lengths at a monomer concentration of 100 nM (a 5.6 ± 1.4-fold decrease) (Figure 3d). The difference in monomer concentrations mean that this ratio of TOF is a lower bound on the relative speed of the v.1 design. The higher TOF of the v.1 system is in turn explained by its higher $k_t$ ($k_t$ for $M_1$v.1 6t = 9.6 ± 0.6 x 10$^5$ M$^{-1}$ s$^{-1}$, Supplementary Figure 21b).

However, this design was discarded as the monomers in solution suffered from a significant leak reaction, with a rate similar to that expected for a 1 nt toehold-mediated strand displacement ($k_{\text{leak}}$ for $M_1$v.1 = 27.7 ± 0.2 M$^{-1}$ s$^{-1}$, Supplementary Figure 21b). This leak reaction would have hindered the precision of the templated system in the specific dimerization regime, with a pool of six different monomers at high concentrations (Figure 5). The final version of the system reduced $k_{\text{leak}}$ for $M_1L$ monomers to 0.29 ± 0.05 $M^{-1}$ $s^{-1}$ (a 96 ± 15-fold decrease, Supplementary Table 14). A sacrifice of reaction rate in exchange for leak suppression was considered a beneficial trade-off that significantly increased the precision of the specific dimerization templating.

We hypothesized that the relatively high $k_{\text{leak}}$ arose from the partial complementarity of $M_x$'s primary toehold domain with $N_y$'s handhold. A leak reaction test with a handhold-less $N_3$ ($N_3$ v.1 (0h)) resulted, however, in only a reduction by half of $k_{\text{leak}}$ ($k_{\text{leak}}$ for strand $N$v.1 (0h) = 11.4 ± 0.7 $M^{-1}$ $s^{-1}$). That this $k_{\text{leak}}$ was still high revealed that the main source of the leak was $M_xL$ binding stability and a possible invasion from the fluorophore-labeled end of the complex. Aiming to reduce $k_{\text{leak}}$, the final version of the dimerization mechanism included 1 nt clamps at 3' and 5' of $L$ strands. In addition, the branch migration domain length was increased to 41 nt to ensure binding stability between $M_x$ and $L$, even in the presence of the two mismatches (Figure 1d).

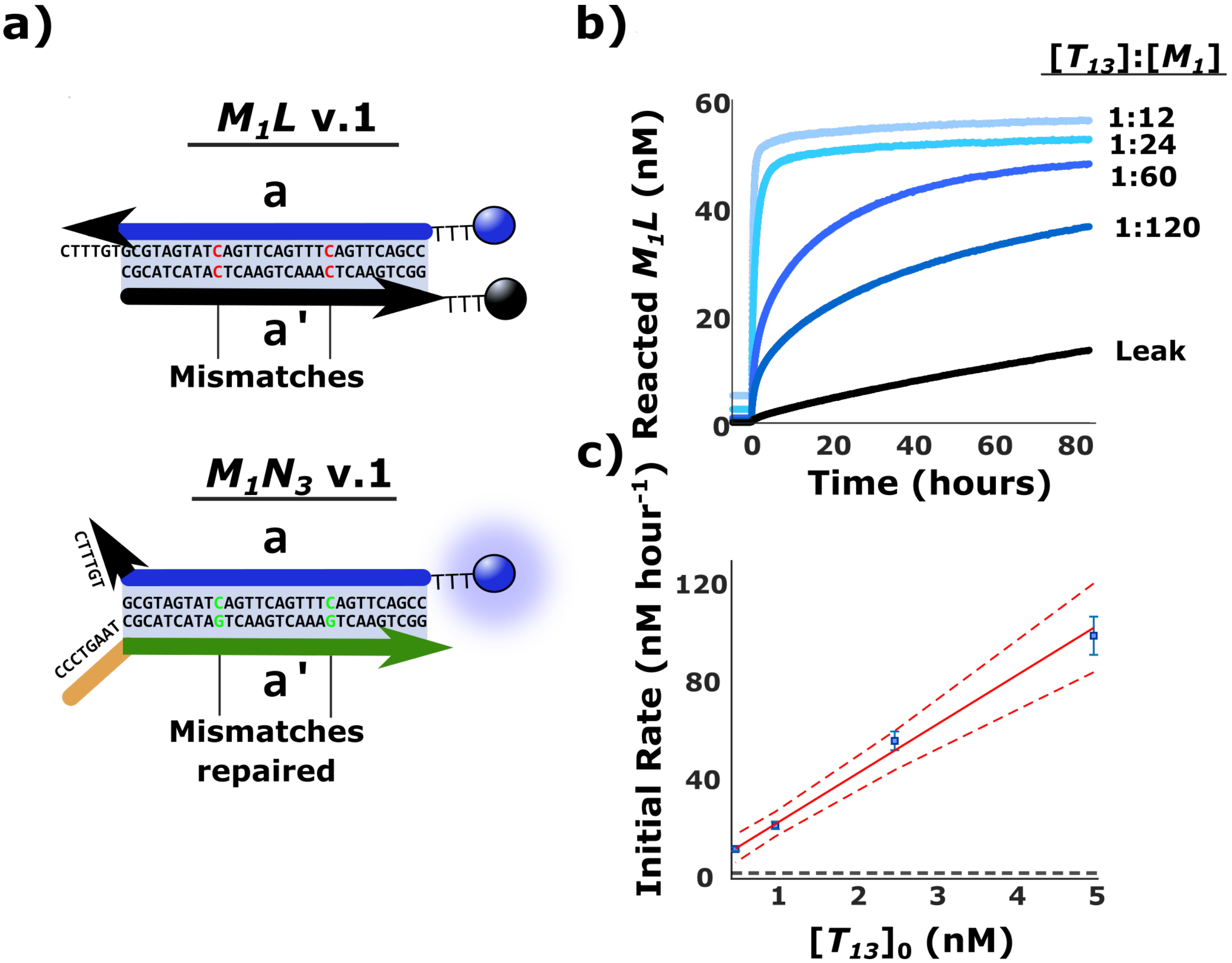


**Supplementary Figure 20: Performance of the clamp-free version of the dimerization system.** a) Detail of v.1 $M_xL$ and $M_xN_y$ complexes. These complexes had a shorter branch dimerization domain $a$ of 24 nt and lacked 1 nt clamps in 3' and 5' extremes of the $L$ strand. Two mismatched base pairs in $M_xL$'s $a$ were still included as a hidden thermodynamic drive, ensuring that dimerization is thermodynamically favored. b) Turnover of $M_1L$ consumption as inferred from fluorescence data for v.1 strands. This simplified version of the dimerization system had a higher TOF but a high $k_{\text{leak}}$. Intended concentrations: $[M_1L \text{ v.1}]_0$ = 60 nM; $[N_3 \text{ v.1}]_0$ = 66 nM; $[T_{13} \text{ v.1}]_0$ = [5, 2.5 1, 0.5 and 0 nM]. c) TOF fitting from the initial rate of reaction from traces in the panel. Blue: experimental data; red dashed line: 95% confidence interval of the fit; red line: fit (TOF = 20 ± 4 hour$^{-1}$); black dashed line: estimated initial rate in the absence of template = 2 ± 2 nM hour$^{-1}$.

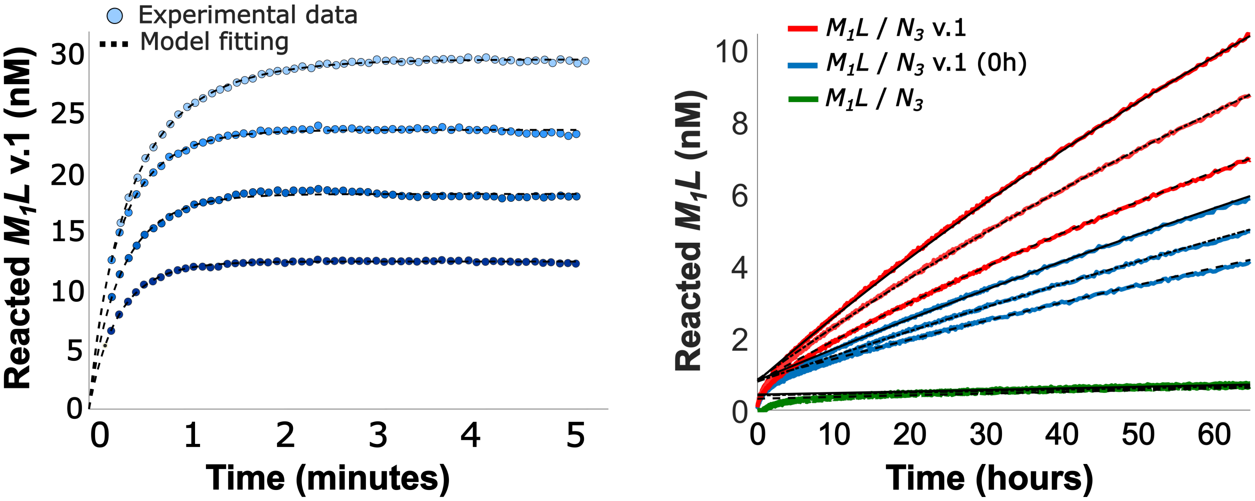


**Supplementary Figure 21: Exploring reaction kinetics for the original version of the displacement domain**. a) Monomer turnover inferred from fluorescence resulting from the binding reaction between $M_1L$ v.1 complex and $T_{13}$ v.1 (6t). The reaction proceeds with a higher rate ($k_t$ = 9.6 ± 0.6 x $10^5$ $M^{-1}$ $s^{-1}$) than its final version counterpart (Supplementary Figure 11). Concentrations: $[M_1L$ v.1$]_0$ = 60 nM; $[T_{13}$ v.1$]_0$ = [30, 24, 18 and 12 nM]. b) Fluorescence resulting from the leak reaction between several designs of $M_1L$ and $N_3$. The removal of handhold from $N_3$ v.1 resulted in a halving of $k_{\text{leak}}$ ($k_{\text{leak}}$ for $M_1N_3$v.1 = 27.7 ± 0.2 $M^{-1}$ $s^{-1}$; $k_{\text{leak}}$ for $N$v.1 (0h) = 11.4 ± 0.7 $M^{-1}$ $s^{-1}$). The addition of clamps and other design constraints in the DNA sequence of $M_1L$ reduced $k_{\text{leak}}$ by two orders of magnitude ($M_1N_3$). Conditions: The form of the model fitting line indicates $[N]_0$: – or 30 nM; -.- for 24 nM; and - - for 18 nM. For v.1 experiments: $[M_1L$ v.1$]_0$ = 60 nM. Conditions for final version: $[M_1L]_0$ = 100 nM.

### 9 Supplementary Results 2: Product yield during sequence-specific dimerization experiments

#### 9.1 Iterations of the experiment

We performed three iterations of the sequence-specific dimerization, as described in Supplementary Table 12 with different proportions of $M_xL$, $N_y$ and $T_{xy}$ (results reported in Figures 4 and 5 and Supplementary Figures 22 to 26). Data was processed as described in Supplementary Section 4. For each iteration, the formation of the $M_xN_y$ products was confirmed through PAGE. The results from the first and second iterations of the experiment, although demonstrating the specific formation of the template-specific products, exhibited appreciable formation of unintended products at long timescales for $T_{13}$. This formation of unintended products was apparently due to conversion of the $M_1N_3$ product onto alternatives $M_1N_x$. This interconvesion represents a slow relaxation towards equilibrium between the $M_1$-containing products.

We were able to largely eliminate this interconversion in iteration 3, after reducing both the concentration of $T_{xy}$ and $N_y$ monomers in the solution. In particular, we found that effective precision, given the possibility of interconversion, increased significantly when $M_x > N_y$. The nature of the mechanism for equilibrium relaxation of the products $M_xN_y$ is further discussed in Supplementary Results 3.

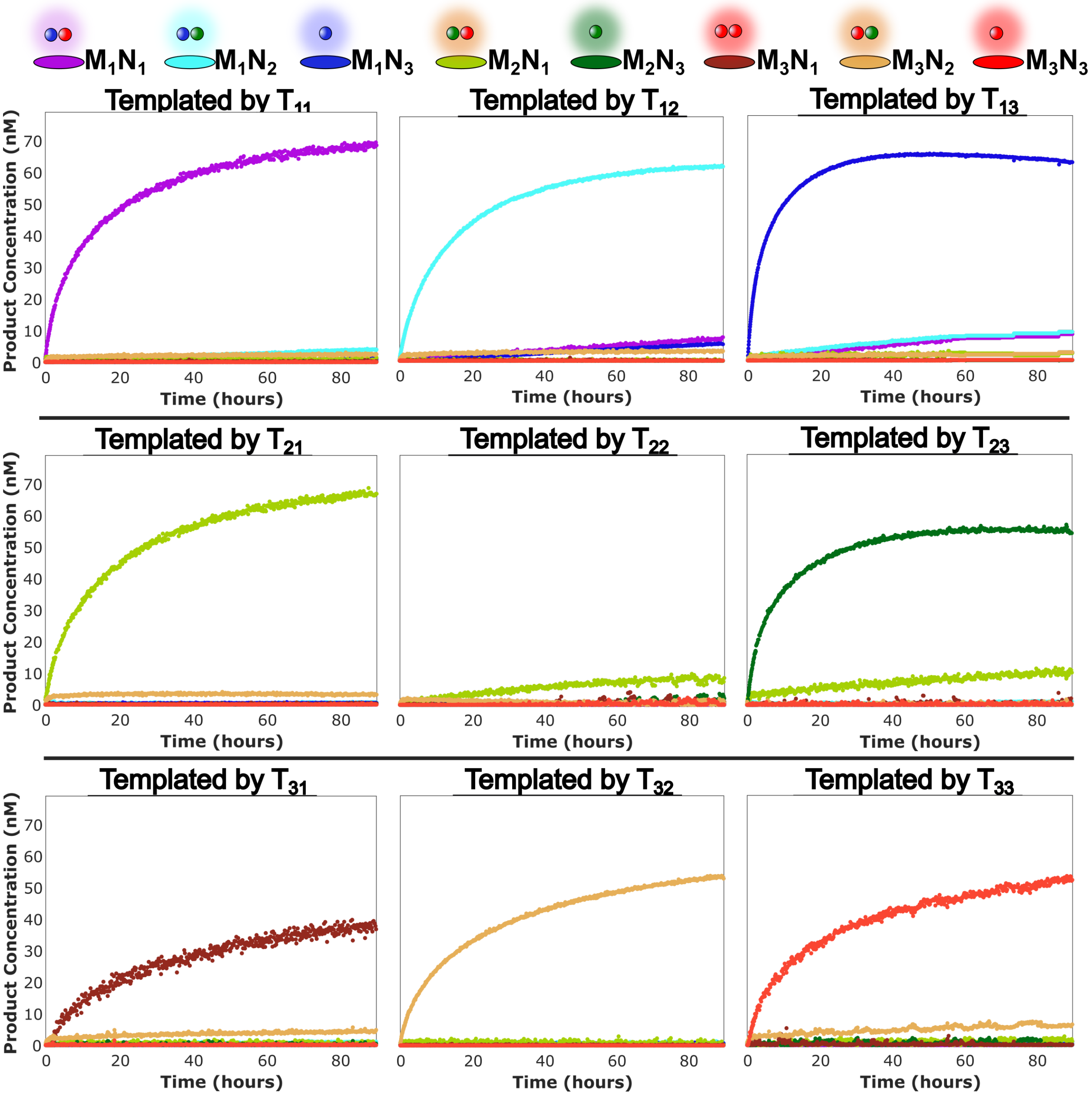


**Supplementary Figure 22: Kinetics of first iteration of sequence-specific dimerization experiment.** For each template, we report product concentrations for all species apart from $M_2N_2$, as inferred using the approach outlined in Section 4. This iteration showed moderate conversion of product $M_1N_3$ into $M_1N1$ and $M_1N_2$. Inferred concentrations of monomers (average of all wells): $[M_1L]_0 = 81$ nM; $[M_2L]_0 = 88$ nM; $[M_3L]_0 = 78$ nM; $[N_1]_0 = 93$ nM; $[N_2]_0 = 99$ nM; $[N_3]_0 = 96$ nM (average of $[N_1]_0$ and $[N_2]_0$). Intended $[T_{xy}]_0 = 5$ nM; intended $[M_3L]_0 = [N_y]_0 = 100$ nM

.

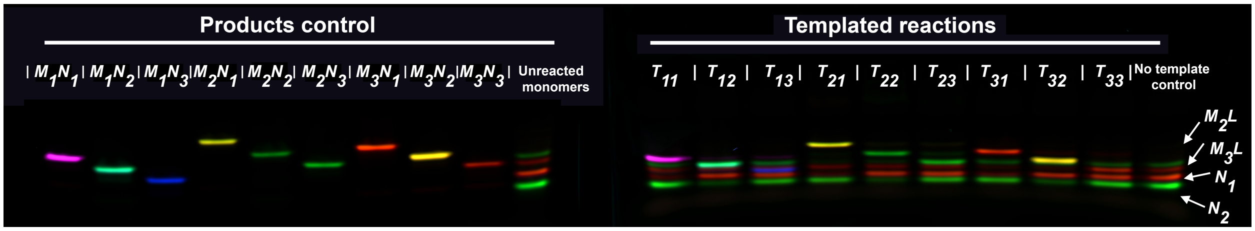


**Supplementary Figure 23: PAGE of first iteration of sequence-specific dimerization experiment.** Fluorescence emission gel electrophoresis of the products of the sequence-specific templating assay, in which a low intended concentration of a single $T_{xy}$ (5 nM) is combined with an intended concentration of 100 nM of each $M_xL$ monomer and 100 nM of each $N_y$ monomer. The annealed controls have the same concentrations of $M_x$ and $N_y$. The gel was run after a 90 hours of the experiment. Observed products are consistent with the intended $M_xN_y$, alongside bands corresponding to the unreacted monomers. False colors: blue, Alexa 488; green, Alexa 546; red, Alexa 647; cyan, FRET 488/546; yellow, FRET 546/648; purple, FRET 488/648)

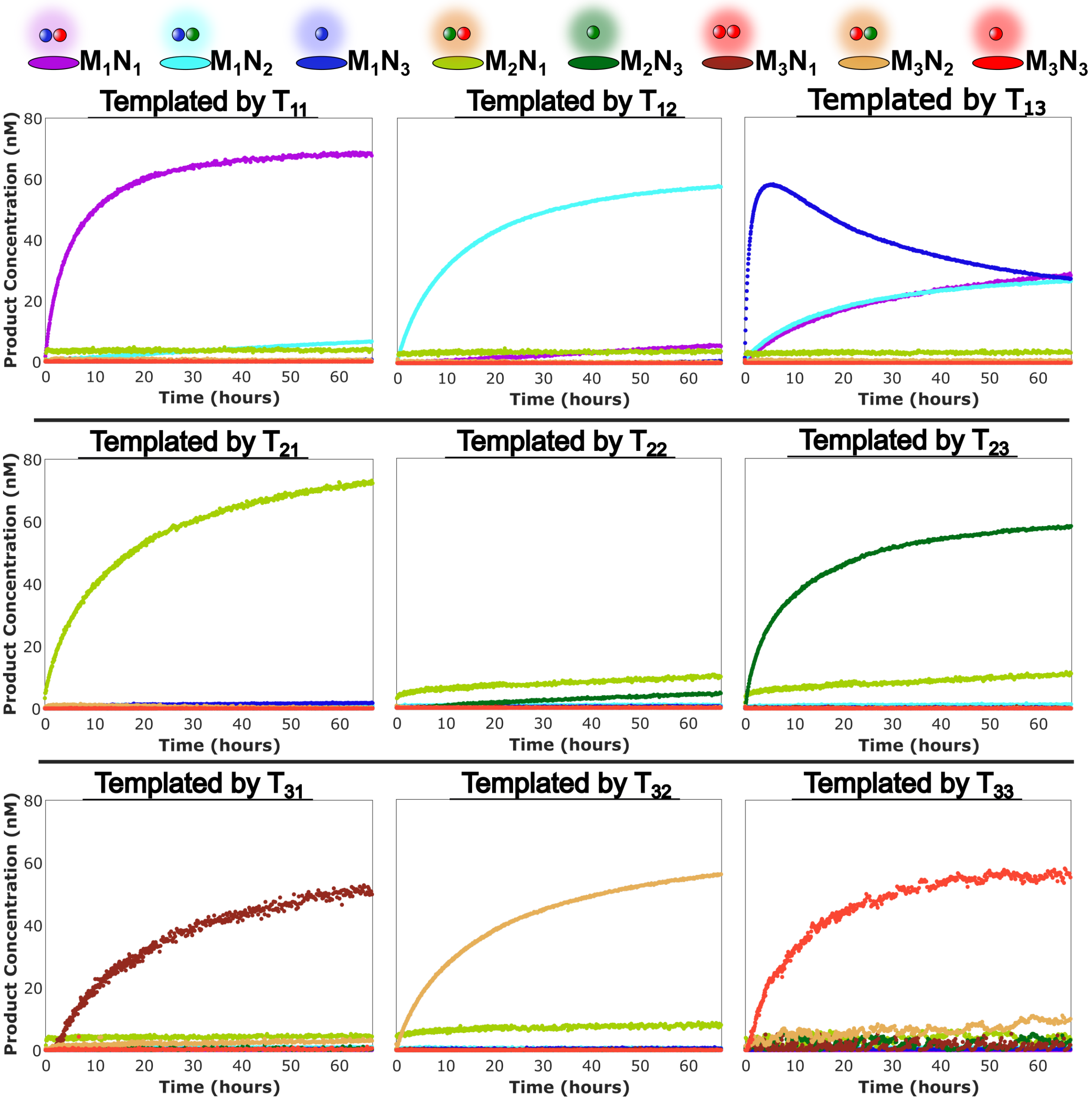


**Supplementary Figure 24: Kinetics of second iteration of sequence-specific dimerization experiment.** For each template, we report product concentrations for all species apart from $M_2N_2$, as inferred using the approach outlined in Section 4. This iteration, with a higher concentration of template, showed a clear tendency towards interconversion of the three $M_1$-containing products in the presence of $T_{13}$. Inferred concentrations of monomers (average of all wells): $[M_1L]_0 = 85$ nM; $[M_2L]_0 = 106$ nM; $[M_3L]_0 = 83$ nM; $[N_1]_0 = 95$ nM; $[N_2]_0 = 95$ nM; $[N_3]_0 = 95$ nM (average of $[N_1]_0$ and $[N_2]_0$); Intended $[T_{xy}]_0 = 10$ nM; intended $[M_3L]_0 = [N_y]_0 = 100$ nM

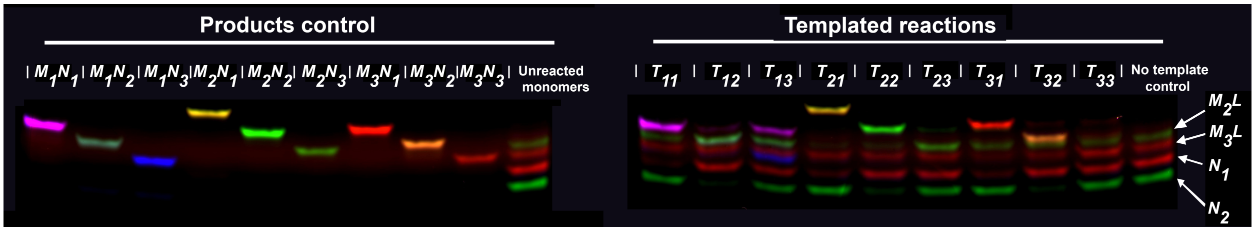


**Supplementary Figure 25: PAGE of second iteration of sequence-specific dimerization experiment.** Fluorescence emission gel electrophoresis of the products of the sequence-specific templating assay, in which a low concentration of a single $T_{xy}$ (intended concentration 10 nM) is combined with an intended concentration of 100 nM of each $M_xL$ monomer and 100 nM of each $N_y$ monomer. Annealed control has the same concentrations of $M_x$ and $N_y$. The gel was run after a 70 hours of the experiment. The condition templated by $T_{13}$ produced similar concentrations of all $M_1$-containing products. Some formation of $M_1N_2$ is also detected for $T_{11}$, $M_2N_2$ for $T_{23}$ and $M_3N_2$ for $T_{33}$. False colors: blue, Alexa 488; green, Alexa 546; red, Alexa 647; cyan, FRET 488/546; yellow, FRET 546/648; purple, FRET 488/648)

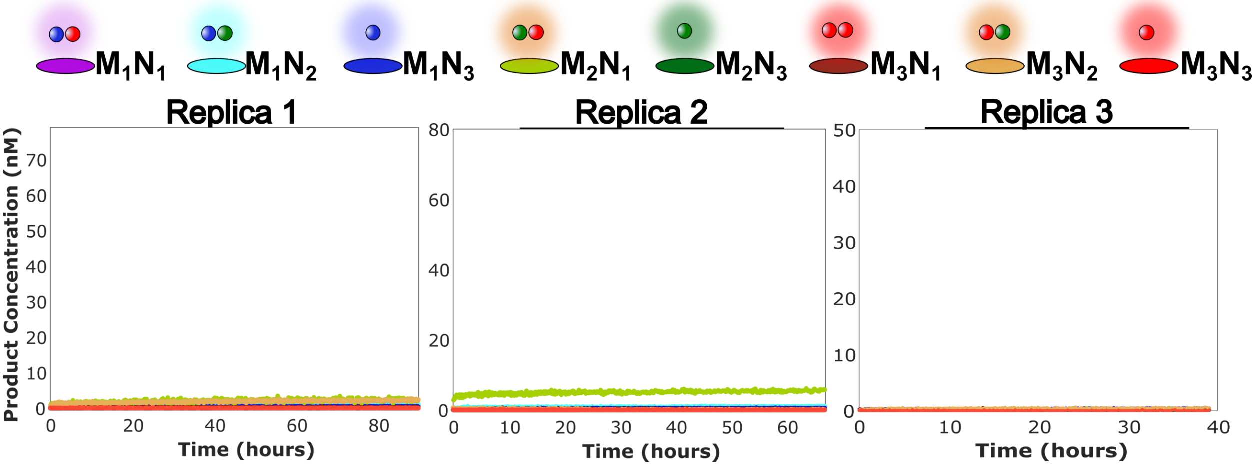


**Supplementary Figure 26: Negative controls for specific dimerization experiments.** Wells containing the same monomer concentrations as the experimental kinetics in their respective iteration in the absence of $T_{xy}$. The results demonstrate that the reaction of the monomers in the absence of $T_{xy}$ is negligible. It also suggests that the $M_2N_1$ baseline in the second iteration may be due to a quantification error, rather than an avctual reaction.

#### 9.2 Quantification of information propagation by templating

In order to estimate information propagation, we calculated the relative yield of each product in the three iterations of the sequence-specific copying experiments (Supplementary Figures 22, 24 and Figure 5 of the main text) after 24 hours of the experiment. Concentrations were estimated from the average of 8 timepoints, around 24 hours. Since the $M_2N_2$ signal could not be detected, for each template other than $T_{2y}$, $[M_2N_2]_{24h}$ was considered as the average of $[M_2N_1]_{24h}$ and $[M_2N_3]_{24h}$. For conditions using a $T_{2y}$ template, $[M_2N_2]_{24h}$ was assumed to be equal to the production of the untemplated $M_2N_y$. In the case of $T_{22}$, $[M_2N_2]_{24h}$ was assumed to be the average of $[M_2N_1]_{24h}$ and $[M_2N_3]_{24h}$ when templated by their respective templates. We justify this approach by noting that the gels did not show $M_2N_2$ as behaving significantly differently from $M_2N_1$ or $M_2N_3$.

To interpret these yields in terms of information, we first calculate the probability of selecting each product dimer from the pool of dimers produced by a template, $P(M_xN_y|T_{vw})$, based on the relative concentrations. These conditional probabilities are reported in Supplementary Tables 19 to 21. We then calculate the mutual information between product and template (in bits) as

$$I = \sum_{xy,vw} \left( P(M_xN_y, T_{vw}) \log_2 \left[ \frac{P(M_xN_y, T_{vw})}{P(M_xN_y)P(T_{vw})} \right] \right), \tag{29}$$

where $P(M_xN_y, T_{vw}) = P(T_{vw})P(M_xN_y|T_{vw})$, $P(M_xN_y) = \sum_{vw} P(M_xN_y, T_{vw})$ and assuming that templates are selected from a uniform distribution $P(T_{vw}) = 1/9$. Values for the mutual information are given in the captions to Supplementary Tables 19 to 21.

**Supplementary Table 19: Templating precision of first specific-dimerization iteration.** Product concentrations are expressed as the percentage of product per template, based on the experiment in Supplementary Figure 22. **Bold**: intended product for used template; *italics*: numbers relying on estimation of $[M_2N_2]_{24h}$. Information content: 2.31 bits out of a possible maximum of 3.17 bits.

| | $M_1N_1$ | $M_1N_2$ | $M_1N_3$ | $M_2N_1$ | $M_2N_2$ | $M_2N_3$ | $M_3N_1$ | $M_3N_2$ | $M_3N_3$ |
|---|---|---|---|---|---|---|---|---|---|
| $T_{11}$ | **91.6** | 2.67 | 0.06 | 1.12 | *0.62* | 0.11 | 0.2 | 3.47 | 0.14 |
| $T_{12}$ | 3.54 | **88.36** | 2.51 | 0.24 | *0.15* | 0.05 | 0.13 | 4.96 | 0.06 |
| $T_{13}$ | 4.58 | 5.92 | **83.35** | 2.35 | *1.22* | 0.09 | 0.08 | 2.39 | 0.01 |
| $T_{21}$ | 0.02 | 0.78 | 0.55 | **91.73** | *0.05* | 0.05 | 0.21 | 6.51 | 0.1 |
| $T_{22}$ | *0.16* | *0.98* | *0.67* | *11.7* | ***83.23*** | *0.42* | *1.27* | *0.7* | *0.88* |
| $T_{23}$ | 0.01 | 0.73 | 0.32 | 9.57 | *9.57* | **79.18** | 0.22 | 0.24 | 0.16 |
| $T_{31}$ | 0.29 | 2.16 | 1.09 | 2.8 | *1.56* | 0.32 | **80.87** | 10.58 | 0.33 |
| $T_{32}$ | 0.34 | 1.43 | 0.93 | 2.89 | *1.46* | 0.04 | 0.23 | **92.68** | 0 |
| $T_{33}$ | 0.06 | 1.5 | 0.65 | 1.99 | *1.92* | 1.86 | 1.04 | 9.55 | **81.43** |

**Supplementary Table 20: Templating precision of second specific-dimerization iteration.** Product concentrations are expressed as the percentage of product per template, based on the experiment in Supplementary Figure 24. **Bold**: intended product for used template; *italics*: numbers relying on estimation of $[M_2N_2]_{24h}$. Information content: 2.00 bits out of a possible maximum of 3.17 bits.

| | $M_1N_1$ | $M_1N_2$ | $M_1N_3$ | $M_2N_1$ | $M_2N_2$ | $M_2N_3$ | $M_3N_1$ | $M_3N_2$ | $M_3N_3$ |
|---|---|---|---|---|---|---|---|---|---|
| $T_{11}$ | **86.65** | 3.98 | 0.02 | 5.31 | *2.75* | 0.19 | 0.16 | 0.89 | 0.05 |
| $T_{12}$ | 3.54 | **85.29** | 0.05 | 6.61 | *3.4* | 0.19 | 0.28 | 0.6 | 0.04 |
| $T_{13}$ | 21.68 | 22.56 | **48.93** | 3.81 | *1.93* | 0.06 | 0.16 | 0.81 | 0.05 |
| $T_{21}$ | 0.12 | 1.47 | 2.11 | **94.42** | *0.18* | 0.18 | 0.26 | 1.25 | 0.02 |
| $T_{22}$ | *0.11* | *1.4* | *0.54* | *12.49* | ***82.05*** | *3.2* | *0.15* | *0.04* | *0.02* |
| $T_{23}$ | 0.002 | 1.56 | 0.22 | 12.04 | *12.04* | **73.84** | 0.17 | 0.04 | 0.09 |
| $T_{31}$ | 0 | 1.78 | 0.72 | 8.88 | *4.82* | 0.75 | **77.76** | 4.77 | 0.5 |
| $T_{32}$ | 0.05 | 0.98 | 0.51 | 13.68 | *6.89* | 0.1 | 0.21 | **77.52** | 0.07 |
| $T_{33}$ | 0 | 1.21 | 0.42 | 4.57 | *4.72* | 4.86 | 1.61 | 9.86 | **72.76** |

**Supplementary Table 21: Templating precision of third specific-dimerization iteration.** Product concentrations are expressed as the percentage of product per template, based on the experiment in Figure 5 of the main text. **Bold**: intended product for used template; *italics*: numbers relying on estimation of $[M_2N_2]_{24h}$. Information content: 2.46 bits out of a possible maximum of 3.17 bits.

| | $M_1N_1$ | $M_1N_2$ | $M_1N_3$ | $M_2N_1$ | $M_2N_2$ | $M_2N_3$ | $M_3N_1$ | $M_3N_2$ | $M_3N_3$ |
|---|---|---|---|---|---|---|---|---|---|
| $T_{11}$ | **94.5** | 1.64 | 0.03 | 0.36 | *0.32* | 0.27 | 0.32 | 2.4 | 0.16 |
| $T_{12}$ | 4.29 | **89.16** | 0.03 | 0.91 | *0.69* | 0.47 | 0.45 | 3.92 | 0.08 |
| $T_{13}$ | 4.66 | 3.69 | **87.49** | 0.48 | *0.26* | 0.04 | 0.22 | 3.07 | 0.08 |
| $T_{21}$ | 0.41 | 0.75 | 1.03 | **94.12** | *0.1* | 0.1 | 0.11 | 3.31 | 0.07 |
| $T_{22}$ | *0.68* | *0.34* | *1.28* | *8.6* | ***87.45*** | *0.19* | *0.25* | *1.14* | *0.06* |
| $T_{23}$ | 0.37 | 0.33 | 1.27 | 9.46 | *9.46* | **78.54** | 0.4 | 0.05 | 0.12 |
| $T_{31}$ | 0.33 | 0.62 | 1.67 | 0.25 | *0.22* | 0.19 | **86.94** | 8.68 | 1.11 |
| $T_{32}$ | 0.71 | 0.87 | 1.54 | 2.79 | *1.45* | 0.11 | 0.37 | **92.14** | 0.02 |
| $T_{33}$ | 0.32 | 0.3 | 0.8 | 0.11 | *0.14* | 0.17 | 0.96 | 9.8 | **87.39** |

### 10 Supplementary Results 3: Mechanism for product interconversion

To explain the conversion of $M_{13}$ into $M_{12}$ and $M_{11}$ observed in Supplementary Figures 22 and 24, we hypothesized that $T_{xy}$, as well as templating the formation of $M_xN_y$, can also increase the rate of relaxation of the dimer distribution towards equilibrium by favoring the interconversion between products that share the same $M_x$.

It is possible for product $M_xN_y$ to rebind to $T_{xy}$, in the process releasing $N_y$ (Supplementary Figure 27a,c). Particularly for reaction regimes where non-template-complementary $N_{y'}$ are at a higher concentration than the template-complementary $N_y$, e.g, after several days of reaction, $T_{xy}M_x$ can then react through a slow TMSD with non-templated $N_{y'}$ (Supplementary Figure 27c). This process would result in the $T_{xy}$-mediated destruction of the initially templated product, and its conversion into $M_xN_{y'}$. The addition of an excess of $M_xL$ over $N_y$ can prevent this destruction mechanism by having the templates preferentially blocked by occupied by $M_x$ when the supply of new monomers runs out. Since there is no intended thermodynamic bias towards one product or another, regardless of which template is present, we would expect all systems to exhibit this relaxation towards equilibrium on sufficiently long-time scales.

We now explore this hypothesis for the cause of product interconversion.

#### 10.1 $t$-complementarity between $T_{xy}$ and $M_xN_y$ is necessary for product destruction

We tested the kinetics of product destruction in several experiments where an initial concentration of $M_1N_3$ (Measurement 1) is mixed with $T_{xy}$ (Measurement 2) and $N_1$ (Measurement 3 - kinetics). Product concentrations were inferred using fluorescence matrix 5 and processed as described in Supplementary Note 4.1. Supplementary Figure 27b shows the results of a set of experiments with intended concentrations of $[M_1N_3]_0 = 50$ nM, $[N_1]_0 = 150$ nM and different $T_{xy}$ with variable concentrations. The results demonstrate both that interconversion requires a template and that this interconversion is $T_{xy}$-dependent. In particular, it mainly relies on complementarity with the primary toehold.

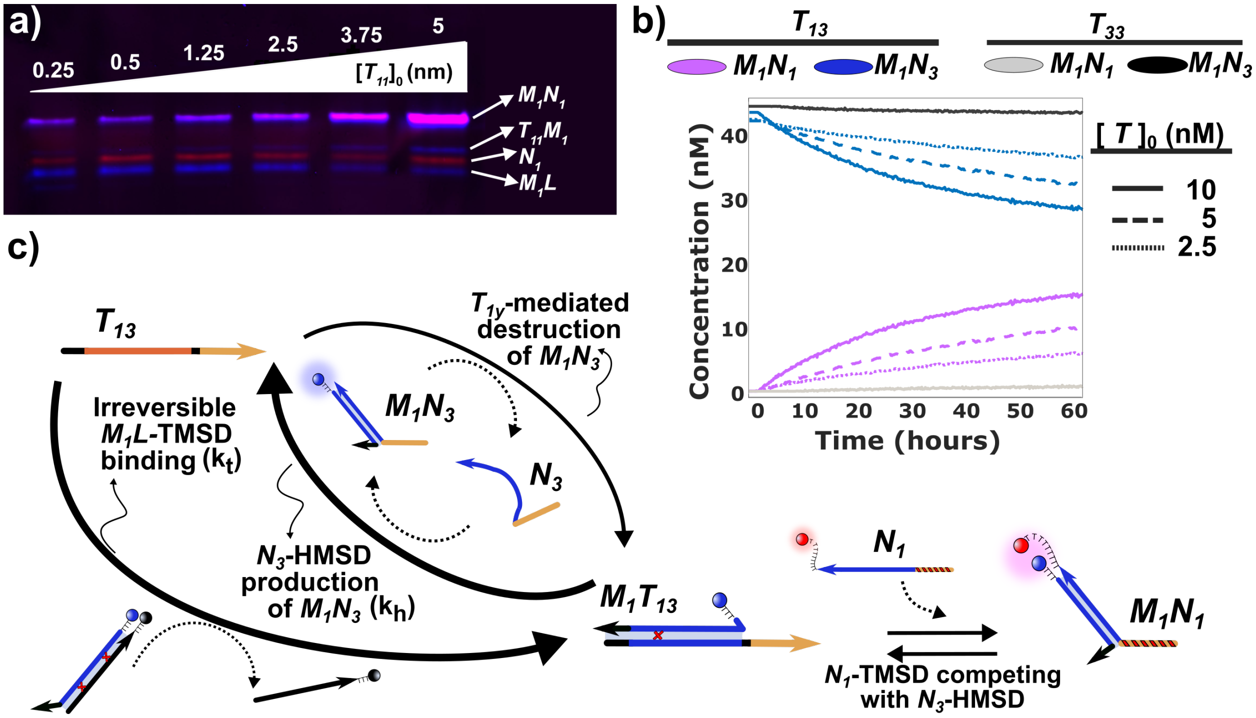


**Supplementary Figure 27: Template-mediated interconversion of products.** a) PAGE of the catalytic production of $M_1N_1$ after over 100 hours of reaction ($T_{11}$ kinetics as shown in Supplementary Figure 17). Even after several days of reaction, the consumption of monomers was not complete. However, the gel demonstrates that the inhibition mechanism is not just the binding of $M_xN_y$ to $T_{xy}$. Rather than being inhibited by the expected three-stranded complex ($M_1N_1/T_{11}$), $T_{11}$ only had $M_1$ attached, hinting at the ability of $T_{11}$ to break apart, rather than just bind to, $M_1N_1$. Intended concentrations: $[M_1 - L]_0 = 100$ nM, $[N_1]_0 = 100$ nM, $[T_{11}]_0 = [0.25, 5]$ nM. b) The template-mediated destruction of $M_xN_y$ needs a $t$-complementary $T_{xy}$. A mixture of $M_1N_3$ and $N_1$ slowly interconverts into $M_1N_1$ in the presence of the product-complementary template $T_{13}$. The rate of interconversion is proportional to the amount of template in the mixture. A non-handhold complementary template ($T_{12}$) has just a slightly slower rate (data not shown). However, removing $t$ complementarity ($T_{33}$) makes the template inert; interconversion is slow, without observable differences in the rate of interconversion for 2.5 or 10 nM of the template. Intended concentrations: $[M_1N_3]_0 = 50$ nM, $[N_1]_0 = 150$, $T_{xy} =$ variable. c) Proposed mechanism for $M_1N_3$ destruction given the experimental evidence. During the dimerization reaction, the produced $M_1N_3$ is in competition with $M_1L$ to reattach to the template. This reattachment is mediated by a $t$-dependent TMSD. An excess of $M_1L$ over the product would prevent the binding of $M_1N_3$ back to the template. While the template is occupied ($T_{13}M_1$), unspecific dimers can form slowly through a secondary toehold-mediated TMSD. The formation of unspecific products through this method becomes relatively more prevalent as the handhold-complementary $N_y$ gets consumed.

#### 10.2 Secondary toehold-mediated TMSD in the absence of handhold-complementarity is $M_xN_y$-destruction limiting factor

To further understand the dynamics of non-templated $M_xN_y$ formation we tested the rate of reaction between the $T_{xy}M_x$ complex and non-handhold complementary $N_y$ (with rate constant $k_{t2}$). An initial concentration of $M_xL$ (Measurement 1) is mixed with $T_{xy}$ (Measurement 2) and a non-handhold complementary $N_y$ (Measurement 3 - kinetics). Product concentrations were inferred as described in Supplementary Section 4.1. An approximate value of $k_{t2}$ was estimated by assuming that the complex $T_{xy}M_x$ is in a steady state during the linear regime of the kinetics fitted (Supplementary Figure 28):

$$k_{t2} = \frac{1}{[N_y]_0[T_{xy}]_0}\left(\frac{1}{\frac{d[M_xN_y]}{dt}} - \frac{1}{[T_{xy}]_0[M_x]_0k_t}\right), \tag{30}$$

with $k_{t2}$ being the rate of the secondary toehold-mediated TMSD; $\frac{d[M_xN_y]}{dt}$ the slope of the linear fit to the kinetics; $[X]_0$ the intended (or estimated in the case of Supplementary Figure 28b) initial concentrations of the reagents; and $k_t$ assumed to be the average $k_t$ for the $M_xL$ used (Supplementary Note 6). The obtained results demonstrate that the TMSD reaction mediated by the secondary toehold is over three orders of magnitude slower than the templated HMSD of the correct monomer. For $T_{x2}M_x$ reacting with $N_3$, $k_{t2}$ with $M_1 = 1.21 \pm 0.06$ M$^{-1}$ s$^{-1}$, with $M_2 = 8.78 \pm 0.04$ M$^{-1}$ s$^{-1}$ and with $M_3 = 2.12 \pm 0.02$ M$^{-1}$ s$^{-1}$(Supplementary Figure 28a). For $T_{12}M_1$ reacting with $N_1$, $k_{t2} = 53$ M$^{-1}$ s$^{-1}$ (Supplementary Figure 28b). We note in passing that the faster invasion of $N_1$ and $N_2$ than $N_3$ is consistent with interactions between the fluorophore labels – absent in $N_3$ – stabilizing both the products $M_xN_y$ and enhancing the toehold-mediated reaction.

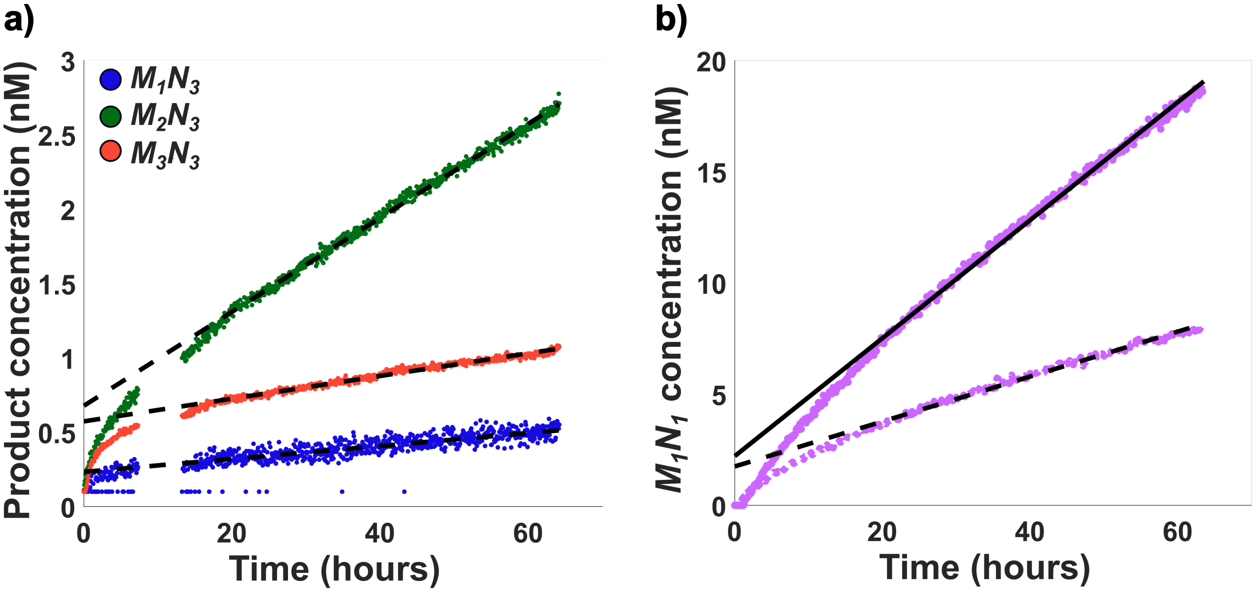


**Supplementary Figure 28: Secondary toehold mediated TMSD reaction without handhold complementarity.** a) TMSD reaction of $T_{x2}M_x$ with $N_3$ in the absence of handhold complementarity. In the absence of handhold interaction, the 2 nt secondary toehold can still mediate a slow displacement reaction. Intended concentrations: $M_{1-3}L$ = 100 nM; $N_3$ = 100 nM, $T_{x2}$ = 10 nM. b) TMSD reaction of $T_{12}M_1$ with $N_1$ in the absence of handhold complementarity. The presence of a fluorophore nearby the secondary toehold domain of $N_y$ increases the rate of untemplated TMSD by one order of magnitude relative to $N_3$. This fluorophore-induced slightly increased stability explains the prevalence of double-labeled incorrect products. Intended concentrations: $T_{12}$ = 10 nM, $M_1L$ = 150 nM and $N_1$ = 100 nM (solid fit line) or 50 nM (dashed fit line).

### 11 Supplementary Results 4: Displacement of $L$ from $M_1L$ by $T_{13}$ is effectively irreversible

To confirm whether $M$-type monomers bind irreversibly to the template, we ran the dimerization reaction (Supplementary Table 9) with varying amounts of additional lock (Supplementary Figure 29). Experiments consisted of a solution with intended concentrations of $[M_1L]_0 = 50$ nM, $[N_3]_0 = 40$ nM, $[T_{13}$ (6t/8h)$]_0 = 10$ nmol dm$^{-3}$ and a variable $[L]_0$ of either 60, 120, 300 or 1000 nM. The control demonstrates that the accumulation of $L$ strand does not produce significant differences between the traces, even for a ratio $[L]_0$:$[T_{xy}]_0 = 100$. The results confirms that $L$ is effectively inert after it has been displaced. Is therefore concluded that premature release of $M_1$ from $T_{xy}$ by $L$ is negligible.

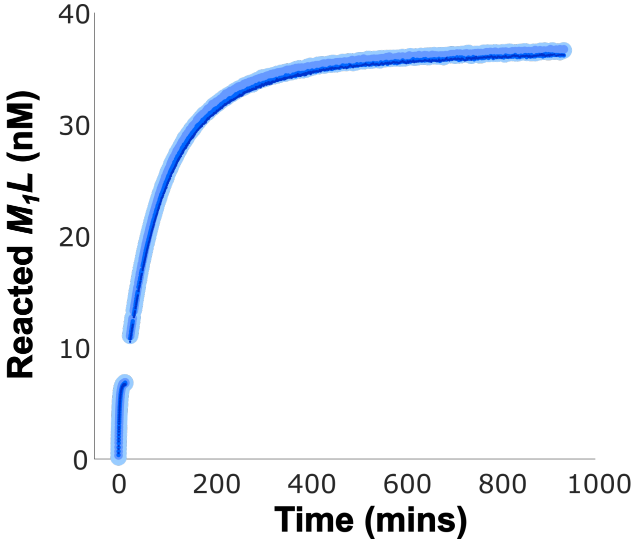


**Supplementary Figure 29: Evidence of the effective irreversibility of the displacement of $L$ from $M_1L$ by $T_{13}$.** Displacement of $L$ from $M_1L$ by $T_{13}$, carried out with background lock concentrations of 60, 120, 300 or 1000 nM. According to the experiment, the lock strand is effectively inert, given that the four replicas exhibit the same kinetics during both sets of kinetics recorded: 1. the initial displacement of $M_1L$ by $T_{13}$, and 2. the addition of $N_3$ to start the catalytic dimerization. To help identify the four near-identical traces, increasing $[L]_0$ conditions are represented with gradually smaller and darker data points. Intended initial conditions: $[M_1L]_0 = 50$ nM, $[T_{13}$ (6t/8h)$]_0 = 10$ nmol dm$^{-3}$, $[N_3]_0 = 40$ nM.

### References

(1) SP&A Application Laboratory, T. F. S. Microplate-Based Pathlength Technical Notes Correction Method for Photometric DNA Quantification. 2015.

(2) The MathWorks, I. Curve Fitting Toolbox. 2020.

(3) Zhang, D. Y.; Winfree, E. Control of DNA strand displacement kinetics using toehold exchange. *J. Am. Chem. Soc.* **2009**, *131*, 17303–17314.

(4) Cabello-Garcia, J.; Bae, W.; Stan V, G.-B.; Ouldridge, T. E. Handhold-Mediated Strand Displacement: A Nucleic Acid Based Mechanism for Generating Far-from-Equilibrium Assemblies through Templated Reactions. *ACS Nano* **2021**, *15*, 3272–3283.

(5) Juritz, J.; Poulton, J. M.; Ouldridge, T. E. Minimal mechanism for cyclic templating of length-controlled copolymers under isothermal conditions. *J. Chem. Phys.* **2022**, *156*, 074103.
